# Gene duplications and phylogenomic conflict underlie major pulses of phenotypic evolution in gymnosperms

**DOI:** 10.1101/2021.03.13.435279

**Authors:** Gregory W. Stull, Xiao-Jian Qu, Caroline Parins-Fukuchi, Ying-Ying Yang, Jun-Bo Yang, Zhi-Yun Yang, Yi Hu, Hong Ma, Pamela S. Soltis, Douglas E. Soltis, De-Zhu Li, Stephen A. Smith, Ting-Shuang Yi

**Author notes:** Co-first author (Gregory W. Stull, Xiao-Jian Qu).

## Abstract

Inferring the intrinsic and extrinsic drivers of species diversification and phenotypic disparity across the Tree of Life is a major challenge in evolutionary biology. In green plants, polyploidy (or whole-genome duplication, WGD) is known to play a major role in microevolution and speciation^1^, but the extent to which WGD has shaped macroevolutionary patterns of diversification and phenotypic innovation across plant phylogeny remains an open question. Here we examine the relationship of various facets of genomic evolution—including gene and genome duplication, genome size, and chromosome number—with macroevolutionary patterns of phenotypic innovation, species diversification, and climatic occupancy in gymnosperms. We show that genomic changes, such as WGD and genome-size shifts, underlie the origins of most major extant gymnosperm clades, and notably our results support an ancestral WGD in the gymnosperm lineage. Spikes of gene duplication typically coincide with major spikes of phenotypic innovation, while increased rates of phenotypic evolution are typically found at nodes with high gene-tree conflict, representing historic population-level dynamics during speciation. Most shifts in gymnosperm diversification since the rise of angiosperms are decoupled from putative WGDs and instead are associated with increased rates of climatic occupancy evolution, particularly in cooler and/or more arid climatic conditions, suggesting that ecological opportunity, especially in the later Cenozoic, and environmental heterogeneity have driven a resurgence of gymnosperm diversification. Our study provides critical insight on the processes underlying diversification and phenotypic evolution in gymnosperms, with important broader implications for the major drivers of both micro- and macroevolution in plants.

## Main Text

Connecting microevolutionary processes with emergent macroevolutionary patterns of phenotypic disparity and species diversification is one of the grand challenges of evolutionary biology. In green plants, polyploidy is a common phenomenon in many lineages^1,2^ that has long been considered a major evolutionary force^3,4^, unique in its ability to generate manifold phenotypic novelties at the population level and in its potential to drive long-term evolutionary trends. However, despite its prevalence and clear microevolutionary importance, its macroevolutionary significance remains unclear and, in major clades such as gymnosperms, understudied. For example, analyses of the relationship between WGD and diversification have produced conflicting results^5,6,7,8^, and no studies to our knowledge have examined whether spikes in gene duplication tend to directly coincide with bursts of phenotypic novelty on a macroevolutionary scale, although clearly gene duplications have facilitated the origins of numerous key traits in plants^9,10,11^. It is possible that other processes — e.g., major fluctuations in population size during rapid speciation events, in concert with extrinsic factors such as climate change or ecological opportunity—may contribute more fundamentally to major phases of morphological innovation and diversification^12,13^. More realistically, however, these various phenomena may interact in complex ways to shape macroevolutionary patterns in plants and, more broadly, across the Tree of Life, and detailed case studies are needed to disentangle their relative contributions to the origins of phenotypic and taxonomic diversity.

Here we examine potential drivers of phenotypic innovation and species diversification in seed plants (=*Spermatophyta*^14^), with a focus on *Acrogymnospermae*^14^, the most ancient subclade of living spermatophytes, including all extant gymnosperms (conifers, cycads, Gnetales, and *Ginkgo biloba*) as well now-extinct groups such as Cordaitales^15^. For the past ~300 million years, seed plants have occupied center stage in terrestrial ecosystems^16^, constituting the most dominant plant clade in terms of both biomass and diversity and forming myriad ecological relationships with other organisms across the Tree of Life, such as bacteria, fungi, insects, and vertebrates^17^. Reconstructing the diversification of seed plants is therefore essential for a better understanding of the history of terrestrial ecosystems across the latter half of the Phanerozoic. Many aspects of seed plant phylogeny remain poorly known or contentious^18^, including the phylogenetic placements of numerous extinct lineages (e.g., Bennettitales and Caytoniales) with respect to living gymnosperms (*Acrogymnospermae*) and flowering plants (*Angiospermae*^14^). Nevertheless, based on the extensive fossil record of seed plants^19^ and phylogenetic studies of living representatives *Acrogymnospermae*^20,21^ (which comprise ca. 1100 species), extant gymnosperm lineages clearly exhibit a complex history of ancient radiations, major extinctions, extraordinary stasis, and recent diversification, but the correlates and causes of major phases of gymnosperm evolution remain understudied. Given that recent studies^22–25^ have challenged the traditional notion^26,27^ that WGD is rare or insignificant in gymnosperms, a focused investigation on the prevalence and macroevolutionary significance of polyploidy in gymnosperms is timely.

We synthesize phylogenomic analyses, trait reconstructions, and examinations of extensive comparative data on a comprehensive phylogeny of gymnosperms to understand major processes that have shaped phenotypic innovation, species diversification, and climatic evolution in this clade. Specifically, we analyze a novel transcriptome dataset including representatives of nearly all extant genera of gymnosperms to identify hotspots of genomic conflict and gene duplication across gymnosperm phylogeny, which we relate to reconstructed bursts of phenotypic innovation and major shifts in species diversification and climatic evolution.

### Gymnosperm phylogenomics, polyploidy, and genome evolution

Based on phylogenomic analysis of our transcriptome dataset (including 790 inferred orthologs across 121 ingroup species, 77 of which were newly sequenced for this study; Supplementary Table 1), we recovered a well-resolved and highly supported species tree for gymnosperms (Fig. 1). We then took a multifaceted approach for WGD inference to circumvent the limitations of some methods and datatypes^28^. Using the species-tree phylogeny and the inferred orthogroups, we performed gene-tree reconciliation to identify hotspots of gene duplication supporting WGD (Fig. 1; Methods), and compared these results against synonymous substitution (*Ks*) plots (Extended Data Figs. 1 and 2; Supplementary Fig. 1; data deposit), as well as reconstructions of chromosome number (Supplementary Fig. 2) and genome size (Extended Data Fig. 3) across a broader sampling of gymnosperm species from our supermatrix phylogeny (Methods). Particular attention was paid to branches with previously inferred (and debated) instances of WGD in gymnosperms, such as the gymnosperm root, Pinaceae, and Cupressaceae^22,29–32^.

**Figure 1.**
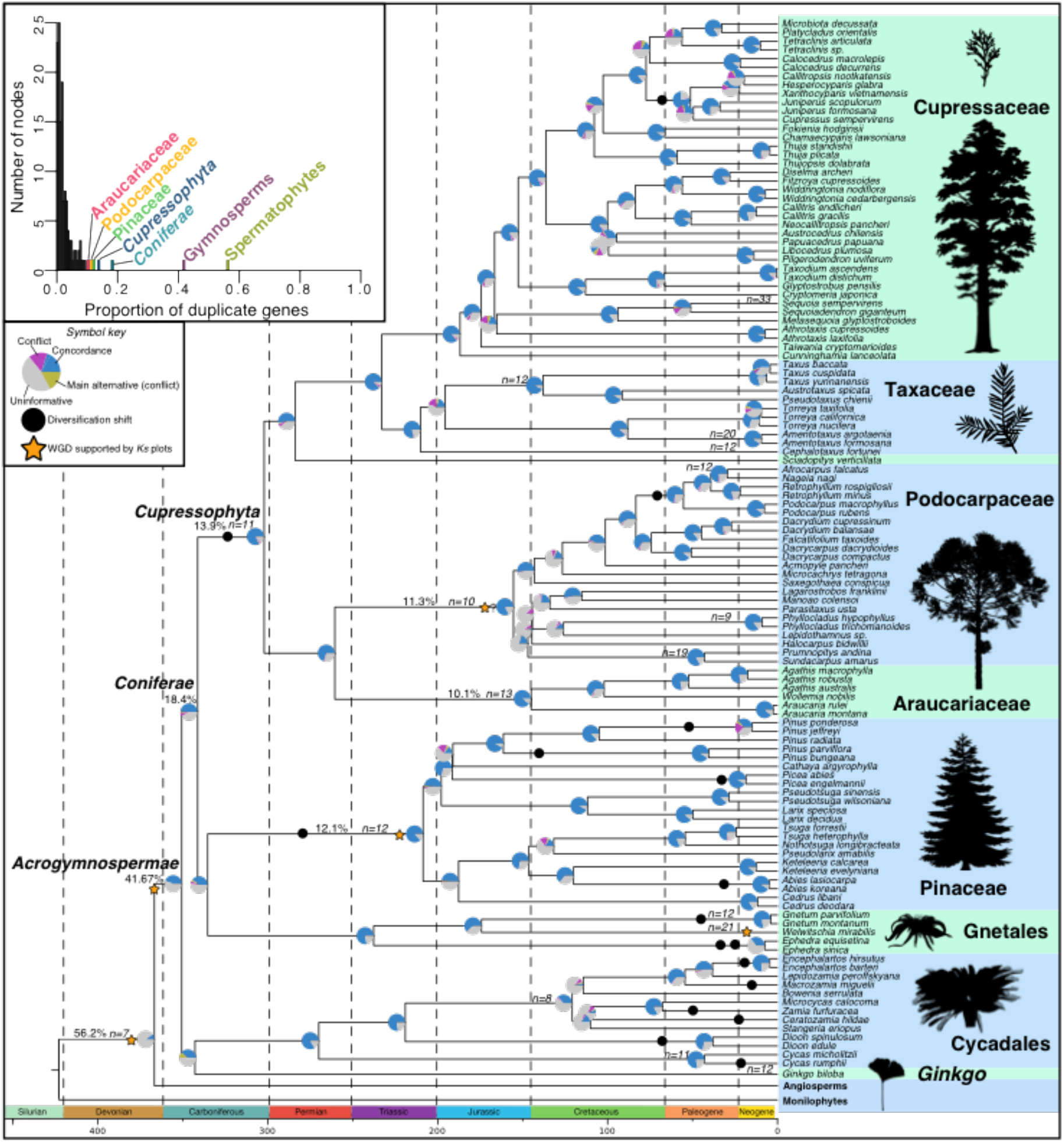
Transcriptome species tree showing major genomic events. Species tree inferred from the 790 gene trees using ASTRAL, scaled to time by fixing the nodes to the ages inferred in the supermatrix dating analysis. Pie charts at each node show the proportion of gene trees in concordance (blue), in conflict (chartreuse = dominant alternative; magenta = other conflicting topologies), or without information for that particular bipartition (grey). Nodes with the highest percentages of duplicate genes (indicative of WGD) are highlighted in the inset plot in the top left corner; the percentages of duplicate genes inferred for these outlier nodes are also displayed on the tree. Orange stars indicated WGD events supported by *Ks* plots; uncertainty regarding the phylogenetic placement(s) of such events is indicated with a question mark. Changes in the haploid chromosome number (based on the ChromEvol analyses; Methods) are indicated on the phylogeny. Black circles indicate diversification rate shifts (inferred using BAMM; Methods). Plant silhouettes were obtained from PhyloPic (http://phylopic.org); credit is acknowledged for the silhouettes of *Araucaria angustifolia* (LeonardoG and T. Michael Keesey); *Zamites* (Robert Gay), which closely resembles the architecture of cycads; and *Ginkgo* (Joe Schneid and T. Michael Keesey). The *Taxus* silhouette was generated from a line drawing of *Taxus brevifolia* obtained from the website of Natural Resources Canada, Canadian Forest Service (https://tidcf.nrcan.gc.ca/en/trees/factsheet/288).

We inferred elevated levels of gene duplication (Fig. 1) in the branches subtending seed plants (56.2 % = percentage of orthogroups showing a duplication), gymnosperms (41.7 %), *Coniferae*^14^ (including Gnetales, 18.8 %), Pinaceae (12.1 %), *Cupressophyta*^14^ (i.e., Araucariaceae + Podocarpaceae + Sciadopityaceae + Taxaceae + Cupressaceae; 13.9 %), Podocarpaceae (11.3 %), and Araucariaceae (10.1 %). Most of these branches also show at least one of the following features: shared *Ks* peaks, inferred changes in chromosome number, or extreme shifts in genome size (Fig. 1; Extended Data Figs. 1–3; Supplementary Fig. 1): seed plants (shared *Ks* peak), *Acrogymospermae* (shared *Ks* peak), *Cupressophyta* (*n* = 7 to *n* = 11), Pinaceae (shared *Ks* peak; *n* = 7 to *n* = 12), Podocarpaceae (*n* = 11 to *n* = 10; major C-value decrease), and Araucariaceae (*n* = 11 to *n* = 13). Cycads and *Ginkgo* show clear *Ks* spikes around 0.8, previously thought to represent a potential WGD event unique to this clade^30^, but these paralog *Ks* peaks are deeper than the ortholog *Ks* peak for gymnosperms as a whole (Extended Data Fig. 1), suggesting instead that these peaks represent a gymnosperm-wide WGD event. This putative gymnosperm peak is also evident in *Ks* plots of conifers, but less prominently and at a *Ks* range of around 1.0 to 1.5. We suggest these *Ks* differences between ‘ginkads’ and conifers might be due to differences in rates of molecular evolution or differences in the timing of diploidization in each lineage if they diverged prior to the diploidization of the polyploid gymnosperm ancestor.

Ancestral chromosomal changes were also observed in Cycadaceae (*n* = 7 to *n* = 11), Zamiaceae (*n* = 7 to *n* = 8), and Gnetaceae (*n* = 7 to *n* = 12), as well as within Cupressaceae, Podocarpaceae, and the genera of Gnetales, due to more recent paleopolyploidy or neopolyploidy^24,25,33^ (Supplementary Fig. 2). Regarding genome size, rate shifts (inferred using BAMM) and/or major ‘jumps’ (i.e., extreme changes in reconstructed ancestor to descendant C-values) in genome size evolution (Methods) were inferred at the origins of several major clades, including abrupt genome downsizing in Podocarpaceae (as noted above), Gnetales, and Cycadaceae and rate shifts in Zamiaceae (excluding *Dioon*), Taxaceae + Cupressaceae, and Ephedraceae; jumps or rate shifts were also inferred in numerous more recent clades (Extended Data Fig. 3). See Supplementary Information for additional results and discussion of gymnosperm genome evolution, including more detailed comparisons of our results with those of previous studies on WGD in gymnosperms^22,29,30,32^.

### Genomic correlates of diversification and phenotypic evolution

Our diversification analysis (Supplementary Fig. 3), based our dated supermatrix phylogeny of extant gymnosperms (including 82% species representation), recovered 17 major shifts, only two of which correspond directly to branches with possible WGDs: *Cupressophyta* and Pinaceae (Fig. 1). Seven other diversification shifts corresponded to major jumps or rate shifts in genome size evolution or changes in chromosome number (Fig. 1, Extended Data Fig. 3, and Supplementary Information). Although most diversification shifts (15/17) were not associated with putative WGDs, it is possible that major extinctions have obscured instances of diversification shifts associated with WGD in deep time. On the other hand, extinction of depauperate branches separating a WGD event and a diversification shift could also lead to their erroneous association. With these considerations in mind, we argue at least that WGD has played a minimal role in driving the more geologically recent diversification shifts detected here. Our results also align with recent studies showing an inconsistent relationship between WGD and diversification^5,6,7^. We suggest that in cases where WGD is associated with diversification, it is a product of WGD mediating changes in traits (such as key innovations^9^) or life history that influence population structuring, with diversification only being a secondary consequence^7^. This is exemplified by Pinaceae, where a possible ancestral WGD is associated with not only a shift in diversification (Fig. 1) but also a major spike in morphological innovation (Fig. 2) and a rate shift in climatic evolution (see below).

**Figure 2.**
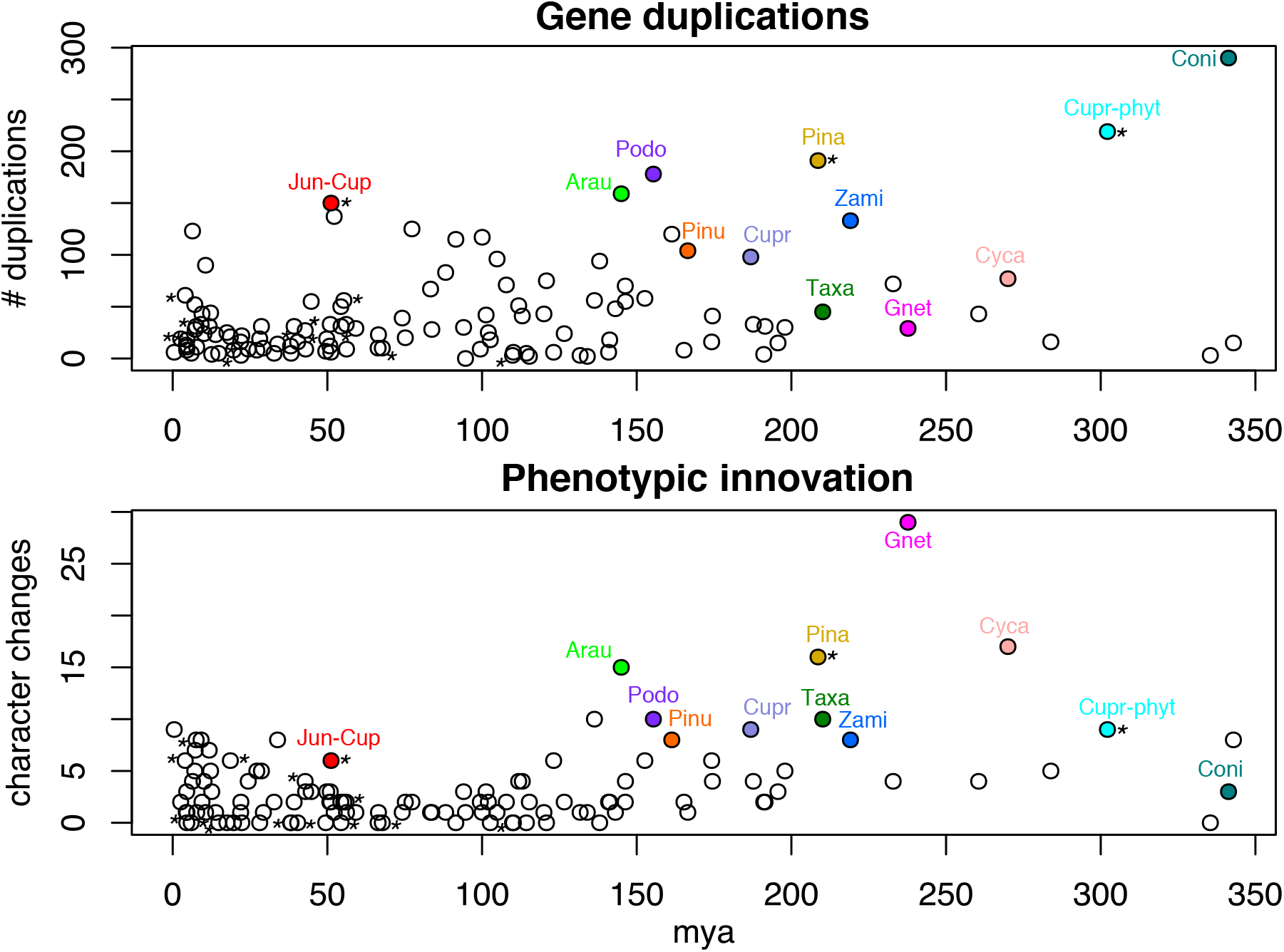
Gene duplications vs. phenotypic innovation. Plots juxtaposing levels of gene duplication and phenotypic innovation (i.e., the number of state changes) at corresponding branches across gymnosperm phylogeny (*n* = 119). The x-axis represents time in millions of years. Nodes with an inferred diversification shift are marked with an asterisk.

To examine the general relationship of gene duplications and genomic (gene-tree) conflict with levels of phenotypic innovation and rates of phenotypic evolution, we analyzed a newly compiled trait dataset for gymnosperms (available in the data deposit) comprising 148 phenotypic characters, with sampling matched to the transcriptome phylogeny (Supplementary Information). Sub- and neo-functionalization of duplicate genes are considered the primary means by which polyploidy can lead to phenotypic novelty^34,35^, and therefore an examination of the direct correspondence of gene duplication numbers to levels of phenotypic novelty at nodes across gymnosperm phylogeny serves as a means of assessing the potential contribution of large-scale (WGD) and small-scale genomic events to macroevolutionary patterns of phenotypic evolution. Genomic conflict, on the other hand, records information about population-level processes during speciation events—e.g., rapid population expansions or fragmentation, resulting in incomplete lineage sorting (ILS), or introgression—that might also play a role in rapid acquisition of phenotypic novelties^36^. Thus, these represent two different, albeit not mutually exclusive, processes that might shape macroevolutionary patterns of morphological evolution in plants. We reconstructed levels of phenotypic innovation or novelty (defined here simply as state changes along a branch) and rates (state changes/time) for each branch in the phylogeny^13^ and plotted these against corresponding levels of gene duplication and gene-tree conflict (Figs. 2 and 3). Linear regression and permutation tests were also performed to evaluate the statistical strength of these relationships (Methods).

**Figure 3.**
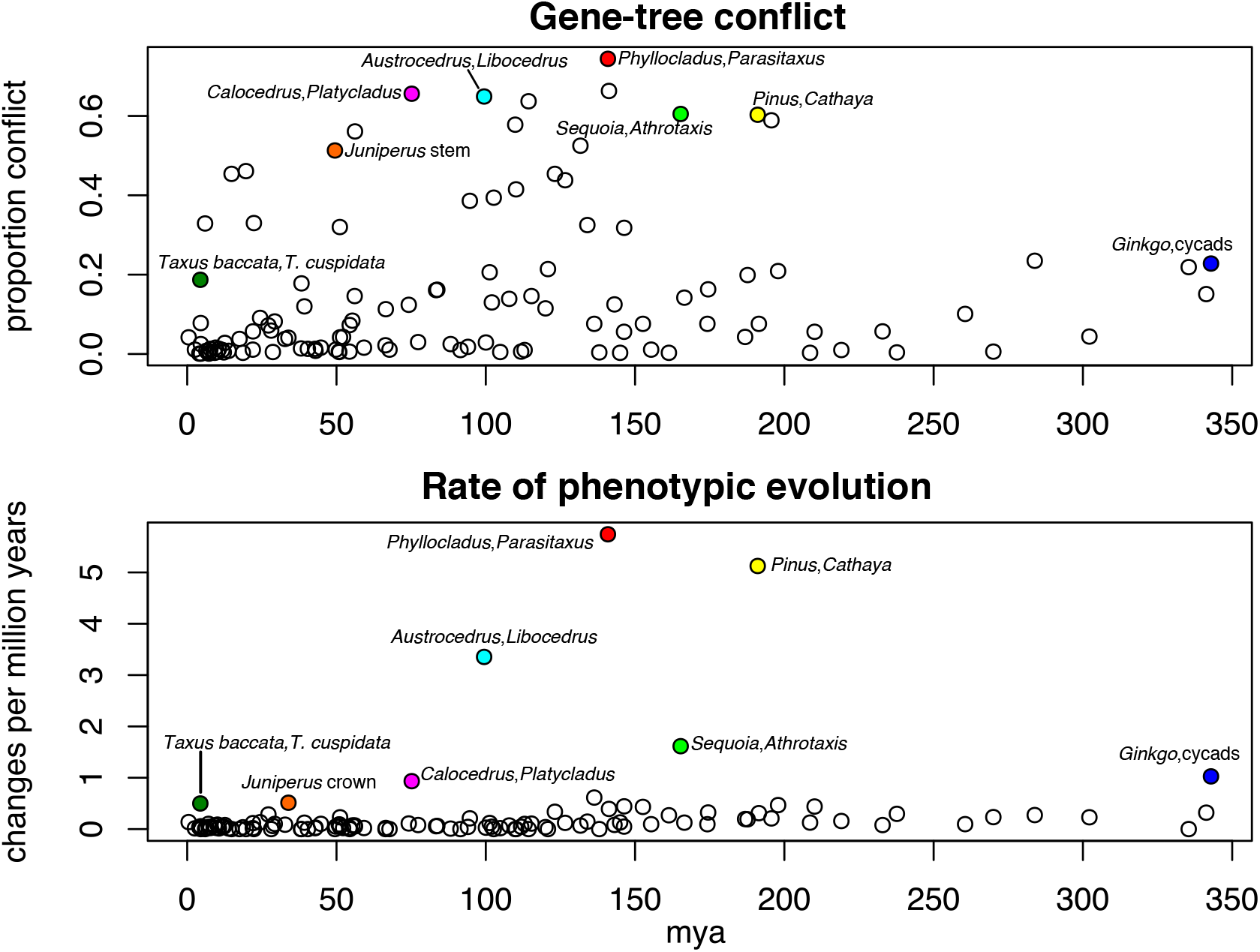
Gene-tree conflict vs. phenotypic rates. Plots juxtaposing rates of phenotypic evolution and levels of gene-tree conflict at corresponding branches across gymnosperm phylogeny (*n* = 119). Note that the conflict node highlighted as corresponding with the *Juniperus* rate node is its immediate parent, comprising *Cupressus sempervirens*, *Juniperus*. The x-axis represents time in millions of years.

We find a positive relationship between levels of phenotypic innovation and levels of gene duplication (linear regression: r^2^ = 0.2038, *p* = 0.000; permutation test: *p* = 0.038; Supplementary Figs. 4 and 5), suggesting that gene duplications may have played a vital role in the generation of phenotypic novelty in gymnosperms evident at a macroevolutionary scale (Fig. 2). Major examples include Araucariaceae, Cupressaceae, *Pinus*, Pinaceae, Podocarpaceae, and Zamiaceae, which all show corresponding, elevated levels of gene duplication and phenotypic innovation. There are also evident temporal patterns, with, for example, a noticeable increase in duplications and phenotypic innovation near the present (Fig. 2). While causation between gene duplication and phenotypic innovation cannot be demonstrated on the basis of these observed patterns, this connection is plausible in cases of complex innovations and clearly deserves future study given that gene duplications have long been considered to play a pivotal role in the origin of evolutionary novelties^34^. It has been suggested that small-scale and large-scale (e.g., WGD) duplication events might tend to have different immediate consequences and evolutionary trajectories^35^. Some of the putative WGD events examined here seem most salient in terms of morphological innovation, but the general correspondence (Supplementary Figs. 4 and 5) suggests that perhaps both large- and small-scale duplication events have played an important role in the generation of novel traits in gymnosperms. It is possible that some of the phylogenetically deeper spikes in phenotypic innovation are exaggerated due to prevalent extinction of stem lineages with intermediate morphologies. However, the sudden appearance of major lineages with distinct morphologies is a common phenomenon across the Tree of Life^13^, and neofunctionalization of duplicate genes (and/or other results of gene or genome duplication) may serve as an important driver of instances of radical and rapid phenotypic change. A better integration of fossil and phylogenomic data might offer important insight into this question.

Although we observed a significant relationship between gene duplication and phenotypic novelty, there are notable discrepancies, the most salient being Gnetales (Fig. 2). However, this is perhaps not surprising given that Gnetales have experienced significant life history changes and stands out among gymnosperms in having relatively small genomes (despite at least recent WGD events^22,25^) and remarkably fast (“angiosperm-like”) rates of molecular evolution^37,38^. *Coniferae* (including Gnetales) also represents a major discrepancy, and the exclusion of Gnetales from the phenotypic reconstructions results in an increase in the number of inferred apomorphies for conifers (Supplementary Fig. 6). However, another explanation for the low levels of innovation inferred for the *Coniferae* node might be the prevalence of traits shared between conifers and *Ginkgo*. Although our species tree places *Ginkgo* sister to cycads (Fig. 1), morphological phylogenetic analyses of seed plants^18^ have typically placed *Ginkgo* with extinct and extant conifers (rather than cycads), and among our gene trees (Fig. 1) the dominant alternative topology for *Ginkgo* is sister to *Coniferae* (with 81 gene trees showing this alternative topology with strong support). This finding suggests that the proposed gymnosperm-wide WGD event might have involved reticulate evolution in the early history of *Acrogymnospermae*, providing a partial explanation for this discrepancy. It is also noteworthy that several well-documented gymnosperm polyploids (e.g., *Sequoia sempervirens, Fitzroya cupressoides*^25^) do not show major phenotypic changes. However, these species both show polysomic inheritance^33^ with gene copies still functioning as alleles, thus constraining the ability of duplicate genes to generate novel phenotypes via neofunctionalization or other processes.

In contrast to cumulative levels of phenotypic innovation, rates of phenotypic evolution show a strong relationship with levels of gene-tree conflict (Fig. 3; linear regression: r^2^ = 0.2308, *p* = 0.000; Supplementary Fig. 7), with extreme values of conflict and rate in particular tending to co-occur (permutation test: *p* = 0.001; Supplementary Fig. 8). This finding is in line with a recent study^13^ documenting a similar correspondence in multiple clades across the Tree of Life. Figure 3 clearly shows that the branches with the highest rates of morphological evolution also have extreme levels of gene-tree conflict. A noteworthy example is the Podocarpaceae subclade including *Phyllocladus* and *Parasitaxus* (Fig. 3), which is characterized by extensive conflict as well as remarkable shifts in phenotype (e.g., the acquisition of phylloclades) and life history (e.g., parasitism) in several of the constituent lineages.

If gene-tree conflict reflects a signature of historical population-level processes such as ILS^39^, the observed relationship between conflict and phenotypic rates suggests that, at macroevolutionary scales, many instances of rapid morphological evolution in gymnosperms are a product of episodic and dramatic population perturbations associated with rapid speciation events. Given the life history characteristics of (most) gymnosperms — including large genome sizes, long time to reproduction, and, in some species, exceptional longevity^40,41^—instances of population reduction or isolation should theoretically be essential for instances of rapid phenotypic evolution. Thus, areas of gymnosperm phylogeny with elevated conflict and morphological rates perhaps reflect the rapid fragmentation of broad, variable ancestral populations. As discussed below, this type of process might explain the prevalence of young conifer and cycad species with highly restricted distributions today. We also note that the processes outlined above are not mutually exclusive putative drivers of phenotypic evolution, given that gene duplications might serve as a source of novelties sorted by population-level processes and that WGD can theoretically result in the rapid establishment of small, reproductively isolated populations^9,42^, which could result in the rapid fixation of polymorphisms via genetic drift.

### Temporal patterns of climatic occupancy evolution and species diversification

Reconstructions of climatic variables on the dated supermatrix phylogeny—focusing on mean annual temperature (Bio 1), mean annual precipitation (Bio 12), and principal components one and two of the 19 WorldClim bioclimatic layers—revealed that rate shifts and jumps in climatic occupancy are concentrated in the last ~70 Myr (and particularly the last ~25 Myr; Supplementary Figs. 9–14). Additionally, many of the branches (10/17) with a diversification shift also included a shift in the rate of climatic evolution (Fig. 4), with four additional diversification shifts separated from a climatic shift by only one or two nodes (i.e., 14/17 diversification shifts are directly or strongly associated with climatic shifts; Fig. 4). Except for two of the concerted shifts (in the branches subtending the cycad genera *Ceratozamia* and *Zamia*), all others involved shifts associated with cooler and/or drier climatic conditions (Fig. 4 and Supplementary Figs. 9–14). Undoubtedly, there have been many deeper bursts of diversification and shifts in climatic occupancy in gymnosperms that are not evident in our phylogeny due to extinction^19^; this renders the inference of diversification dynamics across the depth of gymnosperm history a major, if not insurmountable, challenge without extensive data from the fossil record^43^. Nevertheless, considering an extensive body of research concerning gymnosperm vs. angiosperm functional morphology and competitive dynamics today^44^ and patterns in the fossil record^45^, we might predict many gymnosperms (and especially conifers) to thrive with the expansion of cooler and more arid conditions after late Eocene climatic deterioration and particularly after the mid-Miocene^46^.

**Figure 4.**
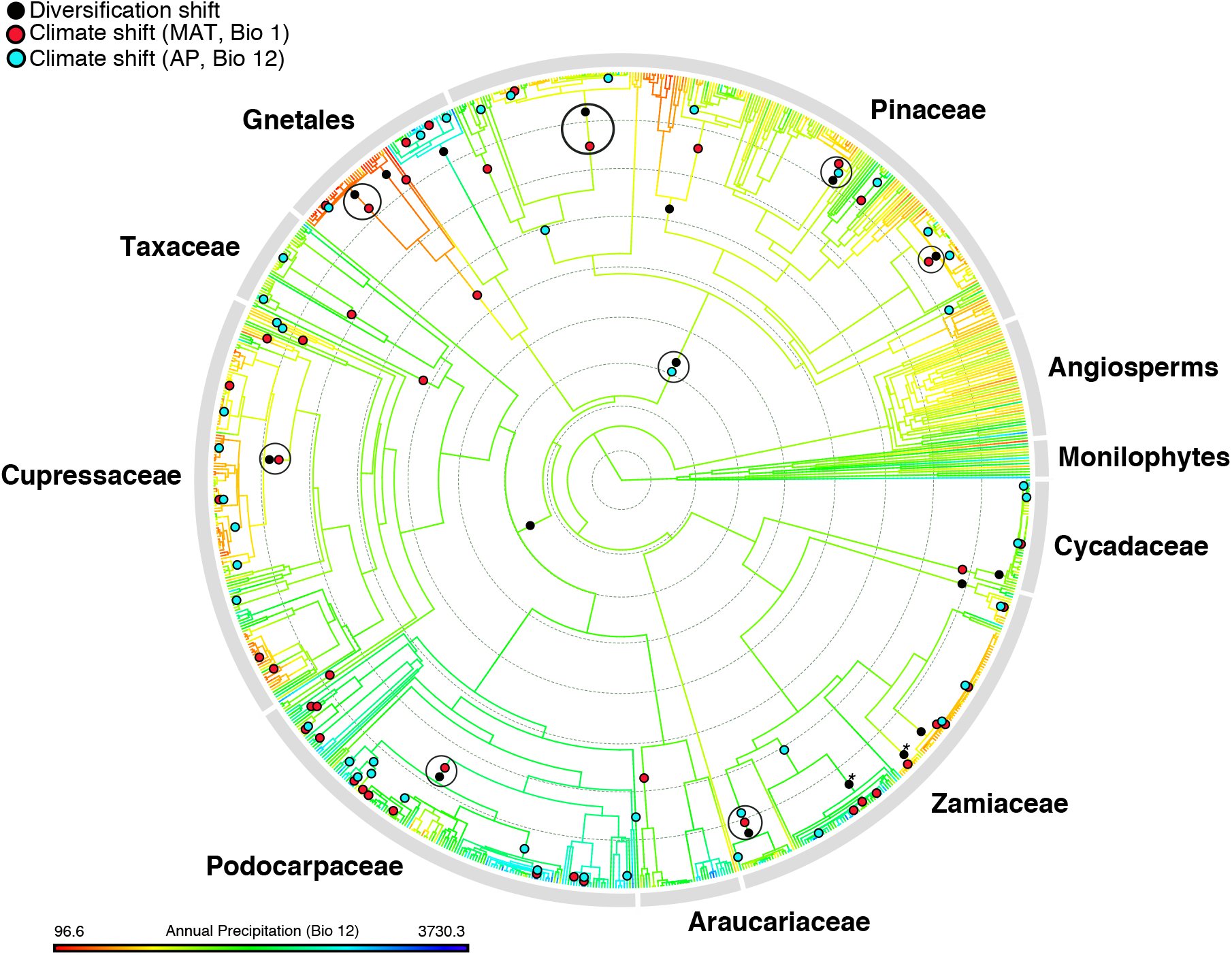
Climatic and diversification shifts across gymnosperms. Ancestral reconstruction of annual precipitation (Bio 12) on the supermatrix phylogeny, showing diversification rate shifts (black circles) and shifts in rates of climatic evolution (Bio 1, mean annual temperature = red circle; Bio 12, annual precipitation = cyan circle) inferred using BAMM. The diversification shifts marked with asterisks (corresponding to *Macrozamia* and *Zamia*) show concerted rate shifts with PC2 of the climatic variables, bringing the number of directly concerted diversification and climate shifts to ten. See Supplementary Fig. 9 for species tip labels.

The global ecological ascendance of angiosperms, from the early Cretaceous to early Cenozoic, was a complex and relatively protracted process, with gymnosperms showing heterogeneous patterns of decline, in terms of abundance and diversity, across different lineages, ecological guilds, geographic areas, and points in time^45,47^. It was not until the early Cenozoic, perhaps in part facilitated by major ecological upheaval at the K-Pg boundary, that angiosperms came to dominate (or co-dominate) nearly all geographic areas and vegetation types^47^. Numerous attributes of angiosperms may underpin their macroevolutionary success, including their reproductive innovations and ecological associations^48^, proclivity for polyploidy^32,49,50^, and faster rates of molecular evolution^37^. However, their functional morphology and correspondingly elevated growth rates have been highlighted as particularly important in the context of competition with gymnosperms^44^. The early Cenozoic was a time of warm, equable climate— conditions well suited to rapid growth strategies of many angiosperms, facilitated by broad leaves and a more efficient vascular system including vessels. Gymnosperms, with narrow tracheids and (generally) smaller, more poorly vascularized evergreen leaves, may have struggled to compete with angiosperms across much of the landscape of the early Cenozoic given the growth-rate limitations imposed by their less efficient vascular systems^44^. Exceptions that prove the rule include *Gnetum* and some members of Podocarpaceae that independently evolved angiosperm-like vasculature and/or broader leaves, allowing them to compete in wet tropical conditions where gymnosperms (especially conifers) are otherwise largely excluded^51^.

However, the same basic traits (and consequent slow-growth strategies) that exclude most gymnosperms from habitats that are more favorable to angiosperm plant growth allow many to persist in more extreme, stressful habitats^52^. The expansion of cool and arid conditions after the Eocene and especially after the mid-Miocene therefore may have created many new opportunities for gymnosperm expansion and diversification. This hypothesis is consistent with our results showing many gymnosperm lineages shifting into cooler and/or more arid conditions after the late Eocene (Supplementary Figs. 9–14). Diversification shifts associated with cool/arid transitions that occurred prior to the late Eocene (e.g., *Pinus* subsection *Ponderosae*, ~56 Ma; the *Juniperus-Cupressus* clade, ~70 Ma; the *Podocarpus-Retrophyllum* clade, ~68 Ma) might represent competitive displacement by angiosperms and/or expansion into upland or montane habitats^53^. Future biogeographic analyses integrating fossil evidence might shed light on the geographic context of these coordinated climatic-diversification shifts. In general, our results challenge suggestions that recent climatic cooling precipitated macroevolutionary declines in conifers^54^ (Supplementary Information). The Northern and Southern Hemispheres, however, clearly do show distinct patterns with respect to the extinction and turnover of conifer diversity during the Cenozoic^21^, and numerous extant gymnosperm species (especially cycads) are at risk of extinction due to pressures including habitat destruction^55,56^.

While climatic shifts have played a significant role in the Cenozoic history of gymnosperms, many major climatic shifts are not associated with changes in diversification (Fig. 4 and Supplementary Figs. 9–14), suggesting that climatic evolution might be an important but insufficient vehicle for diversification. Ecological factors (e.g., environmental heterogeneity related to elevation or soil type) and/or traits (e.g., dispersal strategies) that more directly control population size and spatial structuring are likely necessary concomitant factors for driving diversification patterns in gymnosperms, and more broadly in green plants. This finding is consistent with the observation that many recent gymnosperm radiations (e.g., multiples lineages of cycads, Pinaceae, and Cupressaceae) include numerous species with insular/restricted distributions^57^. The small distributions of many gymnosperm species have typically been interpreted as relictual^40^, but clearly many of these species stem from recent, Neogene radiations given their remarkably short molecular branch lengths^20,21,58^. Aside from these numerous recent radiations, however, extensive data from the fossil record show that many extant gymnosperms — e.g., *Ginkgo biloba*, members of Taxaceae (including *Cephalotaxus*), and the “taxodioid” Cupressaceae—are clearly relics of historically more diverse and broadly distributed groups.

### Macroevolutionary synthesis

Our analyses provide an unprecedented and multifaceted view of genome evolution in gymnosperms, with broad implications for how different attributes of genome evolution might both drive and signal major phases of phenotypic evolution in plants. Using multiple lines of evidence, we show that polyploidy and dynamic shifts in chromosome number and genome size are common occurrences across the phylogenetic breadth of gymnosperms. We also provide compelling evidence for a WGD event in the direct ancestor of gymnosperms, a possibility that has been suggested previously but with limited and conflicting evidence^59^. Furthermore, we provide one of the first direct documentations of a general relationship between levels of gene duplication and phenotypic innovation at a macroevolutionary scale. Gene duplications have long been emphasized as a fundamental source of novelty, but their evolutionary outcomes have typically been studied at a microevolutionary scale or with respect to particular traits or nodes of interest^9,10,11^ (but see Walden et al.^8^). The broader patterns documented here provide a framework of promising areas of gymnosperm phylogeny in which to investigate connections between duplications or gene-family expansions and phenotypic novelties.

Contrary to the association of duplications and innovation, we do not observe a consistent relationship between polyploidy and diversification—consistent with other recent studies also showing a sporadic association^5,6,7^. This implies that, across the euphyllophyte radiation, WGD is more fundamentally tied to phenotypic innovation, and that subsequent diversification is contingent on the nature of the traits acquired and the presence of ecological opportunity^9^. In gymnosperms, recent phases of diversification appear closely tied with shifts in climatic occupancy, suggesting that the expansion of cool and arid climates in the later Cenozoic created new ecological opportunities suited to the basic functional morphology of this clade. Finally, we show that elevated rates of phenotypic evolution appear most closely tied to phylogenetic regions with extensive gene-tree conflict—a signature of dynamic population processes during rapid speciation events. Collectively these threads create a nuanced picture of how different processes—including gene and genome duplication, population dynamics such as ILS and introgression, and ecological opportunity in the face of climate change—may interact to drive the emergence of novel morphologies and major radiations in plants.

## Acknowledgements

We thank the Germplasm Bank of Wild Species at the Kunming Institute of Botany (KIB) for facilitating this study. We thank the curators and staff of the Kunming Botanical Garden of the Kunming Institute of Botany, the University of California Botanical Garden at Berkeley, the Arnold Arboretum of Harvard University, the Missouri Botanical Garden, the Royal Botanic Garden Edinburgh, and the Royal Botanical Gardens Kew for providing fresh and silica-dried leaves and DNA samples. We also thank Andrew Leslie and an anonymous reviewer for their thoughtful comments and suggestions, which greatly improved the manuscript. This work was funded by the Strategic Priority Research Program of the Chinese Academy of Sciences (CAS) (grant No. XDB31000000 to D.-Z.L. and T.-S.Y.), CAS’ large-scale scientific facilities (grant No. 2017-LSF-GBOWS-02 to D.-Z.L.), the National Natural Science Foundation of China [key international (regional) cooperative research project No. 31720103903 to T.-S.Y.], the Yunling International High-end Experts Program of Yunnan Province (grant No. YNQR-GDWG-2017-002 to P.S.S. and YNQR-GDWG-2018-012 to D.E.S.), and the Natural Science Foundation of Shandong province (ZR2020QC022 to X.-J.Q.). G.W.S. acknowledges support from the Chinese Academy of Sciences (CAS) President’s International Fellowship Initiative (No. 2020PB0009) and the China Postdoctoral Science Foundation (CPSF) International Postdoctoral Exchange Program.

## Author contributions

G.W.S., D.-Z.L., S.A.S., and T.-S.Y. conceived the study; X.-J.Q., Y.-Y.Y., Y.H., J.-B.Y., Z.-Y.Y., and H.M. and collected and prepared samples for transcriptome and plastome sequencing; G.W.S. generated the trait dataset, and compiled publically available data for the supermatrix and comparative analyses; G.W.S. conducted analyses with help from C.P.-F., S.A.S., and X.-J.Q; G.W.S, C.P.-F., P.S.S., D.E.S., S.A.S., and T.-S.Y. interpreted the results; G.W.S. wrote the manuscript, with contributions from C.P.-F., P.S.S., D.E.S., S.A.S., and T.-S.Y. All authors approved the manuscript.

---

**Extended Data Figure 1.**
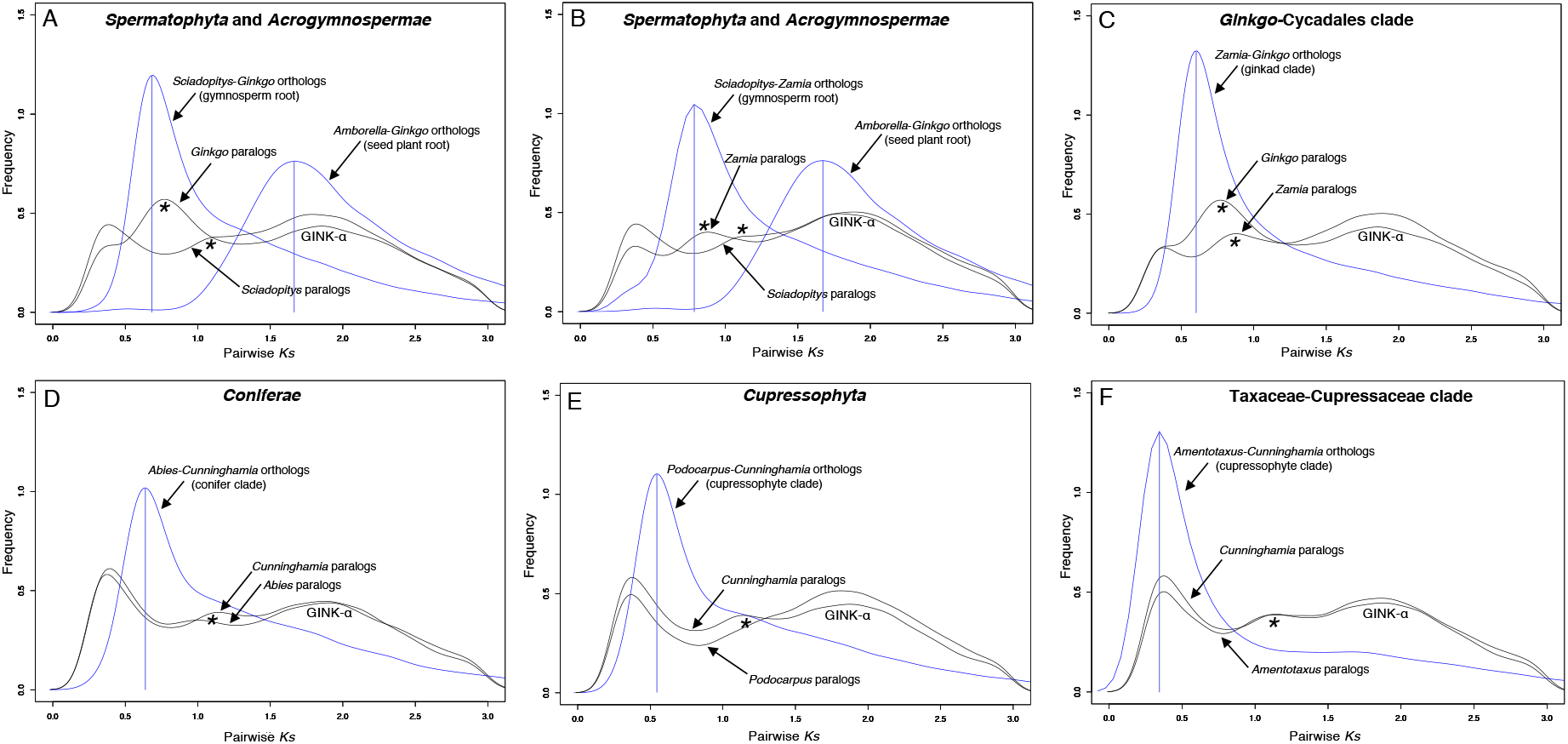
Plots of synonymous substitutions per site (*Ks*) for within-taxon paralog pairs (black lines) and between-taxon ortholog pairs (blue lines). The relative positions of ortholog vs. paralog *Ks* spikes help clarify the phylogenetic positions of possible WGD events. The seed plant WGD event (GINK-α) is labeled. *Ks* peaks corresponding to an inferred WGD for gymnosperms are highlighted with an asterisk. The taxa compared capture the root nodes of (A, B) seed plants and gymnosperms, (C) the ‘ginkad’ clade, (D) conifers, (E) the cupressophyte clade, and (F) the Taxaceae-Cupressaceae clade. Ortholog and paralog *Ks* plots were generated using the pipelines of Walker et al.^83^ and Yang et al.^28^, respectively.

**Extended Data Figure 2.**
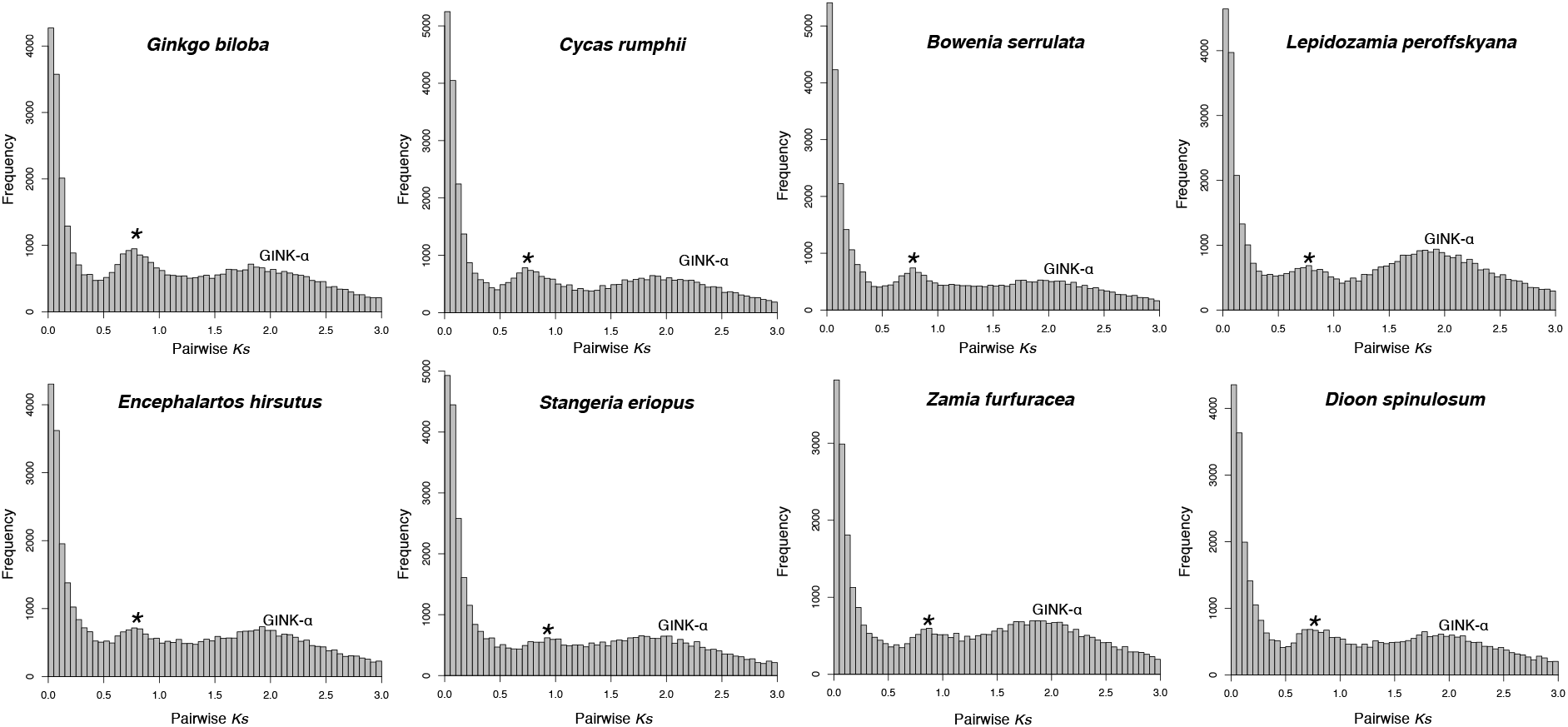
Plots of synonymous substitutions per site (*Ks*) for within-taxon paralog pairs of representatives of the ‘ginkad’ clade. The seed plant WGD event (GINK-α) is labeled. *Ks* peaks corresponding to an inferred WGD for gymnosperms are highlighted with an asterisk.

**Extended Data Figure 3.**
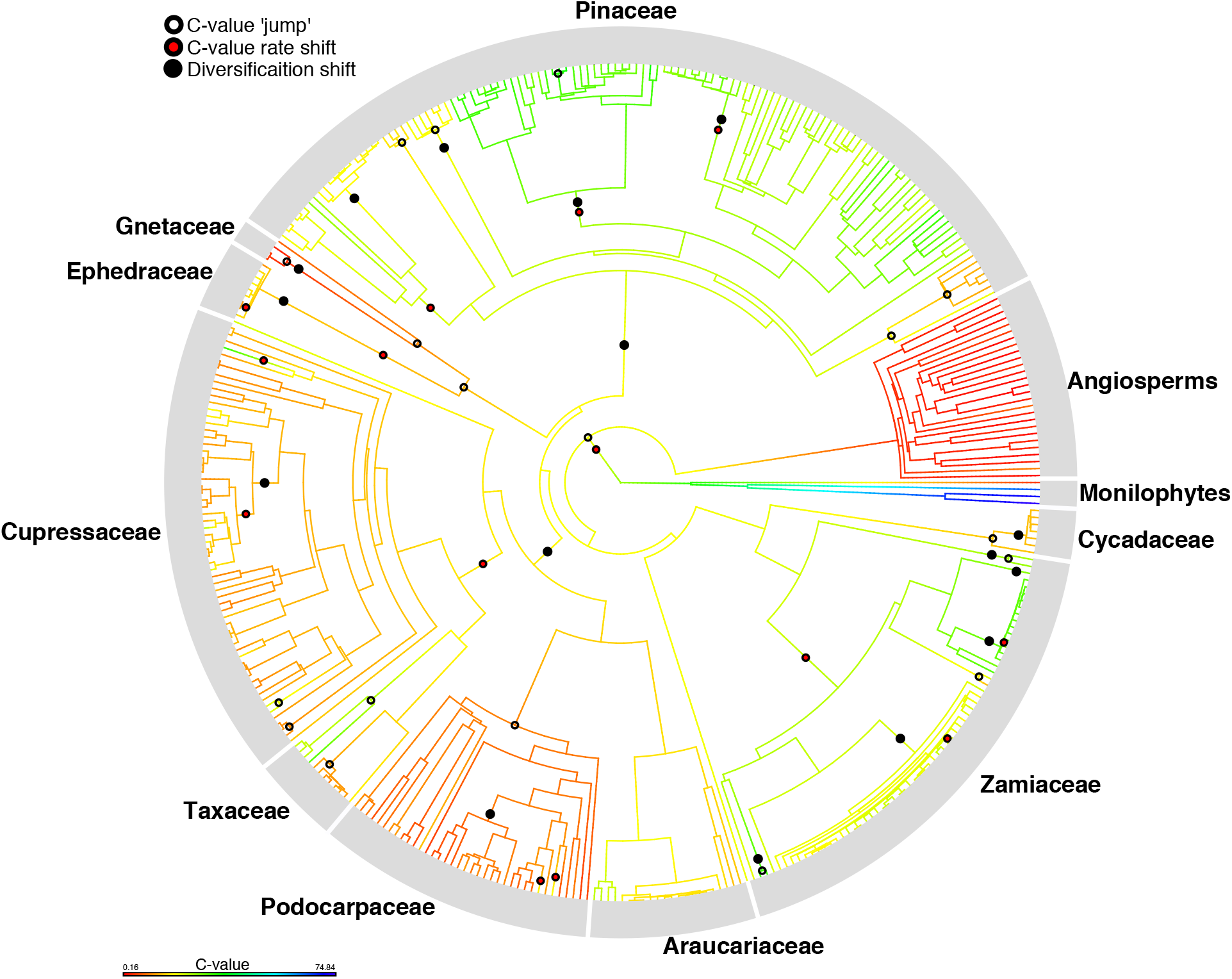
Genome size evolution in gymnosperms. Ancestral reconstruction of genome size (C-value) on the pruned supermatrix phylogeny, showing BAMM rate shifts (red circles) and jumps (i.e., extreme differences in ancestor-descendent values; white circles) in genome size evolution, as well as BAMM diversification shifts (larger black circles). Supplementary Fig. 15 shows this figure with species tip labels.

## Methods

### Sampling overview

We generated two major phylogenetic datasets. The first is a transcriptomic and genomic dataset comprising 144 species (including 121 ingroup species, representing 83 of the 85 gymnosperm genera recognized by the Kew checklist: https://wcsp.science.kew.org/home.do); this was used to resolve broad-scale phylogenetic relationships, characterize patterns of phylogenomic conflict, and infer hotspots of gene duplication. Representatives of angiosperms, monilophytes, and lycophytes were used as outgroups. In all, 77 transcriptomes were newly generated for this study; the remaining transcriptomes were obtained from the One Thousand Plants (1KP) Consortium (https://sites.google.com/a/ualberta.ca/onekp) and several genomes were obtained from the Phytozome database v9.1^60^. See Supplementary Table 1 for transcriptomic and genomic sampling details. The two missing gymnosperm genera, *Pectinopitys* and *Pherosphaera*, represent genera nested within Podocarpaceae and thus their absence should not impact the resolution of major relationships. The newly generated transcriptomes were deposited in the Sequence Read Archive (BioProject PRJNA726756). We also generated a phenotypic dataset of 148 traits for the seed plant species in the transcriptomic dataset to permit examination of the correspondence of different aspects of genomic and phenotypic evolution; the trait dataset is described below.

The other phylogenetic dataset is a plastid supermatrix, based on publicly available plastomes and plastid genes in NCBI (data deposit), as well as 32 plastid genomes newly generated through genome skimming (Supplementary Table 2). In total, this dataset includes 78 plastid genes across 1055 accessions, including 995 ingroup accessions. The ingroup names were checked against the Kew gymnosperm checklist (https://wcsp.science.kew.org/home.do) to resolve issues with synonymy. In total, 890 gymnosperm species are represented, comprising ca. 82% of the 1090 recognized gymnosperm species^56^; our sampling also includes numerous infraspecific taxa (i.e., varieties and subspecies; https://wcsp.science.kew.org/home.do).

The supermatrix dataset was used to generate a comprehensive dated phylogeny for gymnosperms for reconstructions of diversification shifts, climatic evolution, genome size evolution, and chromosome evolution. We used exclusively plastid data for two reasons: (1) plastid genes are more widely sequenced across gymnosperms, allowing for greater species representation; and (2) conflict within phylogenetic datasets can greatly (and negatively) impact branch length estimation and downstream analyses such as divergence-time estimation^61^. While studies show that the plastome is not without conflict^62^, it clearly shows more minimal levels of conflict compared to nuclear genes. Plastid relationships of gymnosperms, as shown by previous studies^56,63^ and our results here, largely agree with those of the nuclear genome, and thus a plastid supermatrix tree should accurately capture major macroevolutionary patterns.

### Transcriptome and plastome sequencing and assembly

To generate the new transcriptomes, total RNAs were isolated from young buds or leaves using the Spectrum™ Plant Total RNA Kit (SIGMA-ALDRICH). Approximately 5 ug of total RNAs were used to construct cDNA libraries (NEBNext Ultra Directional RNA Library Prep Kit for Illumina, Illumina, San Diego, CA, USA). Transcriptome sequencing was performed using the Illumina Hi-Seq 2000 platform. Paired-end reads of 150 bp were *de novo* assembled with Trinity v2.4.0 (Grabherr et al., 2011) using default parameters. Transcriptomes from the 1KP project were previously assembled (Wickett 2014; Leebens-Mack et al. 2019) using SOAPdenovo v1 (Li et al. 2010), and amino acid sequences for three gymnosperm species derived from genome annotations were downloaded from the Phytozome database v9.1^60^. All transcripts were translated using TransDecoder v5.3.0^64^, and then cd-hit-est^65^ (-c 0.99 –n 5) was used to reduce sequence redundancy in all CDS.

Plastid genomes were sequenced and assembled as follows. Total genomic DNA was extracted from fresh leaves using the CTAB method^66^. Approximately one microgram of total genomic DNA from each sample was used for ~400-bp library construction with the NEBNext DNA Library Prep Kit (New England Biolabs, Ipswich, MA, USA) and Covaris-based fragmentation. The Illumina MiSeq instrument (Illumina, San Diego, CA, USA) was used to carry out 150-bp, paired-end sequencing. We assembled the plastomes using both the GetOrganelle pipline^67^ and SPAdes v3.13.0^68^, the latter with the “careful”-option and kmers of 61, 81, 101, and 121. In order to validate the plastome assembly, we mapped all paired-end reads to the consensus sequence of the assembled plastomes with the local-sensitive option of Bowtie v2.3.2^69^. Plastid Genome Annotator^70^ was used to perform plastome annotation, coupled with manual correction in Geneious v8.0.2 (https://www.geneious.com/).

### Phylogenomic workflow

#### Homology and orthology inference

We followed two major procedures for homology and orthology inference in the transcriptomic dataset. First, we employed the hierarchical homology inference approach of Walker et al.^71^. Given the size of our transcriptomic dataset (including 144 transcriptomes), performing an all-by-all BLAST search across all samples for initial homology assessment would have been computationally prohibitive. The Walker et al.^71^ approach overcomes this problem by conducting all-by-all BLAST searches on subsets of the data (*tip clustering*), which are then combined using a post-order (tip to root) tree traversal method (*node clustering)*. We divided our samples into 11 subgroups for tip clustering: **(1)** monilophytes and lycophytes (7 samples), **(2)** angiosperms (16 samples), **(3)** cycads and *Ginkgo* (14 samples), **(4)** Gnetales (5 samples), **(5)** Pinaceae (22 samples), **(6)** Podocarpaceae (23 samples), **(7)** Araucariaceae (6 samples), **(8)** *Sciadopitys* and Taxaceae (12 samples), **(9)** ‘basal’ Cupressaceae (11 samples), **(10)** cupressoid Cupressaceae (17 samples), and **(11)** callitroid Cupressaceae (11 samples); see Supplementary Table 1 for lists of species included in each grouping. For tip clustering, for each subset, we conducted an all-by-all BLASTP search^72^ with an e-value cutoff of 10. The tip clusters were then sequentially combined in a post-order fashion (node-clustering) using the scripts developed by Walker et al.^71^ (https://github.com/jfwalker/Clustering).

For homolog tree and orthology inference from the combined clusters, we followed the Yang and Smith^73^ pipeline (scripts available at https://bitbucket.org/yangya/phylogenomic_dataset_construction/src/master/), which we briefly outline here. Fasta files were written from each cluster. The clusters were then aligned using MAFFT v7^74^ and cleaned using the phyx^75^ program ‘pxclsq’ (retaining alignment columns with at least 10% occupancy), after which homolog trees were built using FastTree v2 with the WAG model^76^. The trees were then pruned of extreme long branches, monophyletic tips were masked (i.e., only one sequence was retained of species/sample-specific alleles/paralogs/duplications), and then long internal branches (representing deep paralogs) were pruned. Fasta files were then written again from the resulting subtrees, and two more iterations of alignment, homolog tree construction, and homolog tree pruning were conducted. A final stage of homolog tree construction was then conducted using RAxML v8.2.12^77^ to generate more robust homolog trees. The resulting 1,992 homolog trees were then used for orthology inference and gene-duplication mapping. For orthology inference, we used the ‘RT’ method of Yang and Smith^73^, which extracts ingroup clades and then cuts paralogs from root to tip. For this procedure, the three lycophyte samples were designated as outgroups (and thus these were pruned and excluded from subsequent analyses), and the minimum ingroup sampling was set at 100. This approach resulted in a set of 790 orthologs. Nucleotide fasta files were then written from each extracted ortholog, re-aligned using MAFFT, and then cleaned by removing alignment columns with under 25% occupancy using ‘pxclsq’ in phyx.

#### Coalescent species-tree inference

Gene trees (from the 790 orthologs inferred above) were constructed using RAxML, using the GTRGAMMA model and 100 bootstrap replicates. The gene trees were then used for species-tree analysis in ASTRAL-III^78^. Prior to the ASTRAL analysis, low supported branches (<10 BS) in the gene trees were collapsed using Newick utilities^79^, as this has been shown to increase accuracy^78^. ASTRAL was then run under default settings, producing a ‘species tree’ tree with local posterior probability (LPP) support values for the bipartitions and branch lengths in coalescent units (Supplementary Fig. 16). The resulting phylogeny was used as a framework for various subsequent analyses, including conflict analysis, analysis of gene duplications, inference of whole-genome duplications, morphological trait reconstructions, and comparisons of phenotypic innovation and rates of phenotypic evolution with levels of genomic conflict and gene duplications.

#### Concatenation species-tree inference

The individual gene/ortholog alignments were concatenated using the phyx program ‘pxcat’. We then inferred phylogenies from the concatenated nucleotide alignment (comprising 1,444,037 bp) using IQ-TREE^80^ with the GTRGAMMA model and ultrafast bootstrapping (1000 replicates). Two analyses were conducted: one unpartitioned, and the other with separate model parameters estimated for each gene region^81^ (Supplementary Fig. 17).

#### Conflict analysis

To examine patterns of gene-tree conflict across gymnosperm phylogeny, we used the program PhyParts^82^, which maps gene trees onto an input species tree and summarizes the number of gene trees in concordance or conflict with each bipartition. When gene trees have inadequate species sampling or bootstrap support (if a support threshold is specified) to speak to a particular bipartition, those are considered uninformative. We use the python script ‘phypartspiecharts.py’ (https://github.com/mossmatters/phyloscripts/tree/master/phypartspiecharts) to visualize the PhyParts results, which are shown in Fig. 1. The levels of conflict calculated for each bipartition were also plotted against corresponding levels of morphological innovation (i.e., the number of new states appearing at each node; see below). We also used the program ‘minority_report.py’ (https://github.com/mossmatters/phyloscripts/tree/master/minorityreport) to examine the number of gene trees supporting alternative placements for particular lineages (namely, *Ginkgo*).

### Inference of whole-genome duplication

We used a multi-faceted approach to examine genomic changes and infer putative WGDs across gymnosperm phylogeny, employing gene-duplication mapping, plots of synonymous substitutions per site (*Ks*), and reconstructions of chromosome number and genome size (C-value). Gene duplications were mapped onto the species tree topology using the scripts (https://bitbucket.org/blackrim/clustering/src/master/) and method of Y ang et al.^28^ This involved extracting rooted orthogroups from the homolog trees (with lycophytes designated as the outgroup), which were then mapped onto the species tree topology. We used two methods of mapping: one maps duplications to the most recent common ancestor (MRCA) in the species tree, with a bootstrap filter of 50% (Supplementary Fig. 18); the other imposes a topology filter such that duplications are only mapped when the sister clade of the gene duplication node contains at least a subset of the taxa present in that clade in the species tree^28^ (Supplementary Fig. 19).

*Ks* plots of within-taxon paralog pairs were generated for the newly sequenced transcriptomes. First, CD-HIT (-c 0.99 –n 5) was used to reduce highly similar sequences, after which an all-by-all BLASTP search was carried out within each taxon (e-value cutoff = 10). The results were then filtered such that sequences with ten or more hits were removed (to avoid overrepresentation of large gene families), as were hits with pident/nident less than 20/50%. *Ks* values were then calculated from the CDS of the remaining paralog pairs, using the pipeline https://github.com/tanghaibao/bio-pipeline/tree/master/synonymous_calculation, and plotted using scripts from Yang et al.^28^ We used between-taxon ortholog pairs to generate ortholog *Ks* plots following the method of Walker et al.^83^; relative comparisons of ortholog and paralog *Ks* peaks are useful for determining whether observed peaks are shared (occurred prior to ortholog/lineage divergence) or more species- or clade-specific (occurred after ortholog/lineage divergence). The ortholog and paralog *Ks* plots (Extended Data Fig. 1 and 2; data deposit) were visually inspected to identify clade-specific shared peaks, which were compared against the results from gene-duplication mapping, as well as the reconstructions of chromosome number and genome size (outlined below). We also generated within-species paralog *Ks* plots using the pipeline of Scott et al.^33^ (https://github.com/nstenz/plot-ks) and these are also provided in the data deposit.

### Supermatrix tree construction and dating

To assemble a comprehensive species-level phylogeny of gymnosperms, we mined plastid gene and genome sequences available on NCBI and combined these with genes extracted from the 32 newly sequenced and assembled plastomes. While the plastomes from NCBI were directly downloaded, individual plastid genes were mined from NCBI using the program PyPhlawd^84^. Large-scale plastid gene clusters obtained using PyPhlawd were then combined with the corresponding individual gene sequences derived from the plastomes.

The plastid genes were aligned individually using MAFFT v7^74^, after which alignment columns with under 25% occupancy were removed using the phyx function ‘pxclsq’. The alignments were then concatenated using the phyx function ‘pxcat’. The final concatenated matrix included 20.35% matrix occupancy (i.e., 79.65% missing data). Prior to phylogenetic analysis, PartitionFinder v2^85^ was used to determine the optimal model partitioning strategy for the concatenated alignment. Phylogenetic analyses were conducted using RAxML-ng^86^ with a topological constraint enforcing major relationships to match the transcriptomic species tree topology: ((*Ginkgo*,cycads),((Gnetales,Pinaceae),((Araucariaceae,Podocarpaceae), (Sciadopityaceae, (Taxaceae,Cupressaceae))))). When unconstrained, the plastome tree only differed from the constraint in the placement of Gnetales. Using RAxML-ng, we conducted 25 independent searches for the maximum likelihood tree (“raxml-ng --search1”), from which the tree with the best likelihood score was selected for downstream analyses of diversification, climate evolution, and genome evolution (chromosome number and cval), although all 25 trees were dated in treePL^87^ to obtain confidence intervals on node ages (see below). We also inferred 100 bootstrap trees to evaluate topological support for the supermatrix phylogeny.

#### Divergence dating

Given the scale of the supermatrix tree (1055 terminals), typical methods of Bayesian divergence dating were unfeasible. Therefore, we used treePL^87^ to generate a dated phylogeny. We used extensive fossil evidence to inform our calibration scheme (see Supplementary Information for detailed discussion of the fossils used, their justification, and references). We conducted independent analyses on the 25 inferred ML trees using the same set of calibrations as a means of obtaining confidence intervals for inferred node ages (Supplementary Fig. 20). Cross-validation was performed for each analysis to determine the optimal smoothing parameter. Node ages from the dated supermatrix tree were then applied as fixed ages (also using treePL) to the corresponding nodes in the transcriptomic species tree. This allowed us to place subsequent analyses involving the transcriptomic phylogeny (i.e., examinations of conflict and rates of morphological evolution) in a temporal context.

Given uncertainty in the placements of two calibrations (for the *Acmopyle* stem and the *Keteleeria* stem; see Supplementary Methods), we also conducted a dating analysis on the best ML tree excluding these calibrations to assess their impact on the ages inferred for major relevant clades (i.e., Pinaceae, Podocarpaceae). The resulting ages for these clades fell within the confidence intervals defined by the analyses of the 25 ML trees noted above (Supplementary Fig. 20), and therefore we proceeded with our initial analyses including these calibrations.

### Chromosome number and genome size reconstructions

Chromosome counts were obtained from the Chromosome Count Database^49^ (http://ccdb.tau.ac.il; accessed 9 December 2019), and the parsed (haploid) counts were used for subsequent analysis. In cases where several different counts were recorded for a given species, the most frequently occurring count was used. Measurements of genome size (i.e., C-values, the amount of DNA in the haploid genome) were obtained from the Plant DNA C-values Database^50^ (https://cvalues.science.kew.org; accessed 9 December 2019). The dated supermatrix phylogeny was pruned to match the sampling in each dataset (chromosome counts and C-values), and these two trees were used in the analyses outlined below. The chromosome count tree/dataset included 478 species, and the cval tree/dataset include 373 species. The chromosome count and C-value datasets examined are available in the data deposit.

Reconstructions of ancestral chromosome number were implemented in ChromEvol v2.0^88^. We examined the data under eight different available models, each including different combinations of the rate parameters *gain* (ascending dysploidy), *loss* (descending dysploidy), *duplication* (polyploidization), and *demi-duplication* (multiplication of the chromosome number by a factor of 1.5), with the rates for each treated either as constant or linear. The model with the lowest Akaike information criterion (AIC) score (i.e., ‘LINEAR RATE DEMI EST’) was used in a more thorough analysis involving default settings and 100,000 simulations. We then visualized changes in chromosome number across the tree using FigTree (Supplementary Fig. 2) and compared changes against reconstructions of genome size and gene duplication.

Ancestral reconstructions of genome size (using C-values) were implemented in R^89^ (R Core Team 2020) using the ‘fastAnc’ function in phytools^90^. The C-value data were not transformed prior to analysis as they conform to a more-or-less normal distribution. To identify the most extreme changes in genome size (which we call C-value ‘jumps’), we then calculated the difference between each node’s reconstructed value and that of its immediate parent node, and selected branches in the 95th percentile of the distribution of parent-child C-value differences. These are shown in Extended Data Fig. 3. We also examined rates of genome size evolution using BAMM^91,92^, using the same tree as above. The priors for the BAMM analysis were determined using the function ‘setBAMMpriors’ in the BAMMtools package^93^. The analysis was run for one billion generations using the phenotypic evolution model, with convergence assessed using BAMMtools. Rate shifts from the maximum a posteriori probability (MAP) shift configuration are shown in Extended Data Fig. 3.

### Species diversification analysis

We used BAMM to detect diversification rate shifts across gymnosperm phylogeny, using the dated supermatrix tree as input. The analysis employed the speciation-extinction model, with the priors determined using the function ‘setBAMMpriors’ in BAMMtools. We also specified sampling fractions specified for each family of *Acrogymnospermae*. This analysis was initially run for 1 billion generations, but then was re-run for 2 billion generations in order to achieve convergence. Rate shifts from the MAP shift configuration are shown in Fig. 1, 4, and Supplementary Fig. 3.

### Reconstructions of climatic occupancy evolution

Geographic occurrences of gymnosperms were pulled from GBIF (https://www.gbif.org; accessed 15 May 2020), and then compared against occurrences from iDigBio (https://www.idigbio.org/portal/search) to check for available data of the species missing from GBIF. Occurrences were then cleaned using the R package CoordinateCleaner^94^, removing duplicate records as well as those with problematic coordinates (e.g., non-terrestrial, identical lat/lon, located within major cities or biodiversity institutions). Synonymy was reconciled using data from the Kew checklist (https://wcsp.science.kew.org/home.do). The occurrences were then used to extract climatic values from the 19 WorldClim bioclimatic variables, and means were calculated for each species for each climatic variable. We dropped tips from the dated supermatrix tree that were absent from the climatic dataset sampling, resulting in a matched sampling of 908 species for subsequent climatic analyses. We then conducted a principal component analysis in R, and principal components (PC) 1 and 2 were used in subsequent analyses of climatic evolution. We also conducted analyses of mean annual temperature (Bio1) and annual precipitation (Bio12), using the mean values for each species.

As with the genome size analyses, for the climate analysis we reconstructed major jumps and rate shifts in climatic evolution, involving separate analyses for PC1, PC2, Bio1, and Bio12. Jumps were determined by reconstructing ancestral climatic values (using the ‘fastAnc’ function in phytools), calculating the difference between each node and its immediate parent, and identifying those branches in the 95th percentile. Extreme climatic jumps for PC1, PC2, Bio1, and Bio12 are shown in Supplementary Figs. 13 and 14. BAMM analysis of rates of climatic occupancy evolution were also conducted on PC1, PC2, Bio1, and Bio12, with each run for 1 billion generations, employing the phenotypic evolution model, with priors determined using BAMMtools. Rate shifts from the MAP shift configuration for Bio1 and Bio 12 are shown in Fig. 4 and Supplementary Figs. 9 and 10; those of PC1 and PC2 are shown in Supplementary Figs. 11 and 12.

Although analysis of historical climatic information would be ideal for understanding the climatic context of previous diversification shifts in greater detail, this absence of reliable estimates spanning the depth of gymnosperm evolutionary history makes this, at the present, not feasible given the scope of our study. Thus, we only used current global climate data for our reconstructions. However, the use of historical climate data to investigate particular Cenozoic radiations of gymnosperms would be a promising avenue for future research.

### Morphological reconstructions and genomic comparisons

A phenotypic dataset comprising 148 traits across the 137 seed plant species included in the transcriptomic dataset was assembled to examine the correspondence of phenotypic and genomic evolution (matrix available in the data deposit). Many of the traits examined were also included in previous morphological datasets. However, the taxonomic sampling in some of these previous datasets was limited; furthermore, many traits from clade-specific datasets needed to be modified and reconciled with similar traits in other datasets in order to establish characters applicable across all sampled taxa. The traits examined (and their corresponding states) are listed in Supplementary Information, and we indicate how these relate to traits in previously published datasets. The primary source datasets were those of Hart^95^, Hilton and Bateman^96^, Mao et al.^97^, Escapa and Catalano^98^, Coiro and Pott^99^, Herrera et al.^100,101^, Gernandt et al.^102^, and Andruchow-Colombo et al.^103^ See Supplementary Information for a complete list of references from which phenotypic data were obtained. We note that while determining homology for character delimitation in seed plants is a major challenge^104^, we do not believe that instances of misidentified homology (i.e., treating independently evolved structures as a single trait or expression of a trait) should necessarily undermine our goals here, given that we are not using this trait set to reconstruct phylogeny. Our goal is to identify state changes along branches, and even in instances of conflated homology, e.g., with two distantly related lineages scored with the same state for a particular trait despite independent origins of the relevant structures, a state change should nevertheless be ‘correctly’ reconstructed and recorded for each branch.

Morphological reconstructions on the species-tree were performed using Maximum Parsimony, following the procedure of Parins-Fukuchi et al.^13^, briefly outlined here. Each trait was reconstructed on the phylogeny, and then the number of state changes along each branch was tallied—this is considered the degree or level of phenotypic ‘innovation’ or ‘novelty’ along each branch. To determine rates of phenotypic evolution for each branch, we divided the number of state changes by the duration of branch (in million years). We then plotted levels of morphological innovation and rates of phenotypic evolution for each branch/node (with the x-axis representing time or node age), and compared these against plots of the same nodes showing their levels of gene duplication and gene-tree conflict (Figs. 2 and 3), allowing for visual inspection of these patterns and their correspondence through time. Because Gnetales was detected as a major outlier, with extreme levels of morphological innovation perhaps driven by life history shifts, we also conducted morphological reconstructions excluding Gnetales to ensure that the inclusion/placement of Gnetales was not significantly biasing reconstructions for other near-by nodes (Supplementary Fig. 6).

On the basis of the visual correspondences observed from the plots noted above, we then explicitly tested the relationships of (1) phenotypic innovation vs. gene duplication and (2) phenotypic rates vs. gene tree conflict by conducting linear regression and permutation analyses of each pair of variables. Linear regression was performed using the ‘lm’ function in R, treating phenotypic innovation and rate as the dependent variables in the respective analyses. For the examination of innovation vs. duplication, we conducted an analysis including all nodes (Supplementary Fig. 21), as well as one excluding two major outlier nodes: Gnetales and conifers (Supplementary Fig. 6). As noted in the main text, we believe exclusion of these outliers is warranted given that (1) Gnetales has experienced major changes in genome size (i.e., reduction) and life history with generally elevated rates of molecular evolution that might erase signatures of a relationship between gene duplication and morphological innovation, and (2) trait reconstruction for conifers is complicated by its inclusion of Gnetales (a highly derived lineage) and the prevalence of features shared among conifers and *Ginkgo*, which may have been involved in a reticulation event early in the evolution of the conifer lineage based on patterns of gene-tree conflict.

The permutation test, developed by Parins-Fukuchi et al.^13^, identifies nodes with significantly elevated values for a particular variable by resampling (with replacement) observed values 1000 times and using the means from each replicate to create an empirical null distribution. We executed this procedure for each of the four variables considered above (duplication levels, conflict levels, innovation levels, and phenotypic rate), with nodes/values falling in the 95th percentile deemed significant. Then, to examine the strength of the association between (1) innovations and duplications and (2) rates and conflict, we determined whether the number of nodes with significantly high values for both of the variables in question (e.g., innovations and duplications) departed from the expectation if the same number of nodes was randomly sampled (Supplementary Figs. 5 and 8). This permutation test was designed specifically to examine whether extreme/significant values (in, for example, conflict and rates) tend to coincide, even if there is not a strict positive monotonic relationship between the two variables across the spectrum of values. Finally, we also used this test to determine if diversification shifts tend to co-occur with significant/extreme levels of phenotypic innovation (Supplementary Fig. 22).

### Data availability

The newly generated raw sequence data is available at the NCBI Sequence Read Archive (https://www.ncbi.nlm.nih.gov/sra) under BioProject PRJNA726756 (transcriptomic samples) and PRJNA726638 (genome skimming samples). The newly assembled plastid genomes are also available at NCBI (https://www.ncbi.nlm.nih.gov); see Supplementary Table 2 for sample accession numbers. Sequence alignments, phylogenies, *Ks* plots, phenotypic trait data, and other data analyzed (chromosome counts, c-values) are available on figshare (10.6084/m9.figshare.14547354).

