## Supplementary Information for "Gene duplications and phylogenomic conflict underlie major pulses of phenotypic evolution in gymnosperms"

### 1. Supplementary Methods

#### Phylogenetic concepts and scope of the present study

This study focuses on the clade comprising living gymnosperms, i.e., Cycadales, Gnetales, conifers (in the traditional sense), and *Ginkgo*. Many important clades that we discuss here do not have traditional (ranked) taxonomic names available, or their composition has varied, making their usage unclear. For these clades, we use names established in the PhyloCode[14], and these are presented throughout in italics. ***Acrogymnospermae*** comprises all living gymnosperms, i.e., Cycadales, Gnetales, traditional conifers, and *Ginkgo*; it is a node-based definition and, therefore, when we refer to the age of *Acrogymnospermae* we are referring to the age of the crown clade formed by its constituent extant lineages. ***Coniferae*** comprises all traditional conifer families (Pinaceae, Araucariaceae, Podocarpaceae, Sciadopityaceae, Cupressaceae, and Taxaceae, with the latter including Cephalotaxaceae) as well as Gnetales. *Coniferae* was defined such that it would either comprise all traditional conifer families to the exclusion of Gnetales, or all conifer families including Gnetales, depending on the phylogenetic position of Gnetales. Because our study shows Gnetales sister to the conifer family Pinaceae (consistent with other recent studies[38,105]), we thus use this name in the broader sense, encompassing both traditional conifers and Gnetales. ***Cupressophyta*** is a subclade of *Coniferae*, comprising the families Araucariaceae, Podocarpaceae, Sciadopityaceae, Cupressaceae, and Taxaceae.

The composition of the *Acrogymnospermae*[106,107] relative to the colloquial term "gymnosperm" merits further elaboration, given that the term "gymnosperm" has different meanings depending on whether fossil diversity is considered. Gymnosperms (or seed plants with "naked seeds") in the broad sense are essentially seed plants to the exclusion of angiosperms (where angiosperms are defined as seed plants with fruits, or enclosed seeds), and phylogenetic analyses including fossil representatives[18] clearly show that gymnosperms in this sense are a paraphyletic group (with respect to angiosperms). However, living gymnosperms (or *Acrogymnospermae*) do indeed form a clade, positioned sister to angiosperms[32]. Throughout this study, when we refer to "gymnosperms", we are referring to this clade of living gymnosperms unless clearly stated otherwise. There remain major outstanding questions in seed plant phylogeny, including the placements of extinct lineages such as Bennettitales and Caytoniales with respect to *Acrogymnospermae* and the angiosperm clade. It is possible, for example, that Bennettitales are related to Gnetales[108], and thus placed within *Acrogymnospermae*. The extinct lineage Cordaitales is also likely placed within the *Acrogymnospermae*[107]. This is not inherently problematic for the concept of *Acrogymnospermae*; it simply means that it might include additional, now-extinct major lineages, which is to be expected for most major clades across the Tree of Life.

**Fossil calibrations used for molecular dating**

The following fossils were used as calibrations in the treePL dating analyses of the supermatrix-based phylogeny. For each calibration, we list the fossil taxon name(s), the corresponding age(s), the extant taxa defining the calibration node (i.e., the most-recent common ancestor, mrca), the relevant citation(s), and justification for its use. Wherever possible, we used fossils according to their placement in previous phylogenetic analyses.

1. **Euphyllophyte crown**: min 388.0, max 423.0 Ma; mrca = *Ginkgo biloba*, *Osmunda angustifolia*. The minimum age for the euphyllophyte crown is based on the fossil taxon *Rellimia*[109], the earliest fossil evidence of progymnosperms, which, by extension, represents a minimum age for the split of seed plants and monilophytes. The oldest fossil evidence of crown vascular plants, *Baragwanathia*[110], was used to set a maximum bound for the euphyllophyte subclade. See also Beaulieu et al.[111].
2. **Leptosporangiate crown**: min 251.0 Ma; mrca = *Osmunda angustifolia*, *Blechnum polypodioides*. This calibration is based on the fossil taxon *Palaeosmunda* sp.[112], which is interpreted as the earliest fossil evidence of leptosporangiate ferns and has been used as such in previous dating analyses[111]. We used this fossil as a minimum constraint (at 251 Ma) on the leptosporangiate crown.
3. **Spermatophyte crown**: max 366.8 Ma, mrca = *Amborella trichopoda*, *Ginkgo biloba*. We used the earliest fossil evidence of the seed plant lineage (*Elkinsia polymorpha*, 363 to 366.8 Ma[113]) to set a maximum bound on the age of crown seed plants. The fossil record and phylogeny of seed plants[18,96,104] clearly show that *Elkinsia* is, if not ancestral, at least early-diverging with respect to seed plants, and thus it is reasonable to assume that crown seed plants should not predate the earliest fossil seed plants.
4. **Angiosperm crown**: min 132.9 Ma, max 139.8 Ma; mrca = *Amborella trichopoda*, *Gentiana verna*. Based on the earliest known fossil evidence of putative crown angiosperms[111], comprising pollen with reticulate-columellate wall structure from the

Valanginian-Hauterivian boundary, approximately 132.9 Ma[115,116]. The calibration minimum age was set to that of the pollen, with the maximum age set to that inferred by Magallon et al.[117]

5. **Eudicot crown**: min 125.0, max 132.9 Ma; mrca = *Papaver somniferum*, *Gentiana verna*. The minimum age for eudicots was set based on the first appearance of tricolpate pollen[118,119], a synapomorphy for this clade[120]; the maximum age is based on the earliest evidence of crown angiosperms. The appearance of angiosperm lineages in the fossil record follows expectations from phylogenetic relationships, and continued paleobotanical work continues to support the notion that the radiation of crown angiosperms was a Cretaceous phenomenon[114]. Therefore, we reasoned that the age of eudicots should not exceed the earliest angiosperm fossil evidence.

6. ***Acrogymnospermae* crown**: min 306.2 Ma, mrca = *Ginkgo biloba*, *Cycas circinalis*, *Pinus bungeana*, *Gnetum parvifolium*. *Cordaixylon iowensis*[121,122] represents the oldest fossil evidence of putative conifers[123], and has been placed among conifers in previous phylogenetic analyses[96]. Despite phylogenetic uncertainties, as well as the complications of Gnetales likely being a member of the 'conifer' clade based on molecular evidence, this fossil nevertheless should serve as a reasonable minimum age for *Acrogymnospermae* (including *Ginkgo* and cycads, as well as conifers) as a whole, which is how we used it here.

7. **Araucariaceae + Podocarpaceae clade**: min 213.0 Ma; mrca = *Araucaria heterophylla*, *Podocarpus macrophyllus*. This calibration is based on the fossil *Araucarities rudicula*[124], which represents perhaps the best early macrofossil evidence of the Araucariaceae stem, and thus can be used as a minimum constraint on the Araucariaceae + Podocarpaceae node. This fossil was similarly used in Beaulieu et al. [111].

8. **Araucariaceae crown**: min 145.0 Ma; mrca = *Araucaria heterophylla*, *Agathis borneensis*. Several fossil taxa of Araucariaceae from the middle to late Jurassic (e.g., *Araucaria brownii*, *Araucaria mirabilis*, *Araucarites bindrabunensis*[125]) have been phylogenetically placed in crown Araucariaceae[98], and therefore we used the late Jurassic-early Cretaceous boundary as a minimum age for the divergence of crown Araucariaceae.

9. ***Agathis* crown**: min 52.2 Ma; mrca = *Agathis australis*, *Agathis silbae*. The fossil *Agathis zamunerae*[126] represents some of the best early unequivocal fossil evidence of this genus and was suggested to likely represent a crown member of the genus. Thus, we used the fossil as a minimum constraint on the crown of *Agathis*.

10. ***Acmopyle* stem**: min 55.2 Ma; mrca = *Acmopyle sahniana*, *Podocarpus macrophyllus*. Eocene fossil evidence of *Acmopyle*[127] is used to calibrate the stem of this lineage, i.e., the bipartition defined by the two taxa above. However, given some uncertainty concerning the placement of this fossil, we also conducted an analysis without it to ensure that its inclusion was not overly influential in the main analysis.

11. ***Dacrycarpus* crown**: min 52.5 Ma; mrca = *Dacrycarpus vieillardii*, *Dacrycarpus cinctus*. The fossil seed cones *Dacrycarpus puertae* and associated organs[128] represent convincing early evidence of crown *Dacrycarpus*, and thus we used the age of this fossil to set a minimum age on the divergence of extant *Dacrycarpus*.

12. **Cupressaceae crown**: min 182.7 Ma; mrca = *Cunninghamia lanceolata*, *Athrotaxis laxifolia*, *Juniperus virginiana*. The fossil taxon *Austrohamia minuta*[129] represents early fossil evidence of crown Cupressaceae, supported by previous phylogenetic analyses[97]. We therefore used this fossil as a minimum constraint on the Cupressaceae crown.

13. ***Athrotaxis* stem**: min 113.0 Ma; mrca = *Athrotaxis laxifolia*, *Metasequoia glyptostroboides*, *Juniperus virginiana*. *Athrotaxis ungeri*[130] represents the oldest fossil evidence of the extant genus *Athrotaxis*, but phylogenetic analyses[97] indicate a stem position, with some uncertainty in its precise placement. We thus used this fossil to constrain the age of the subclade of Cupressaceae excluding *Cunninghamia* and *Taiwania*, positioned outside the clade including the fossil.

14. ***Callitris* complex stem**: min 27.8 Ma; mrca = *Callitris endlicheri*, *Actinostrobus pyramidalis*, *Neocallitropsis pancheri*. Fossil evidence of *Callitris*, represented by *Callitris leaenis*[131], was used as a minimum constraint for the clade including *Callitris* and close relatives, based on the phylogenetic placement of this fossil from Mao et al.[97]

15. ***Fitzroya* stem**: min 27.8 Ma; mrca = *Fitzroya cupressoides*, *Diselma archeri*. The Oligocene fossil taxon *Fitzroya acutifolia*[131] was used to calibrate the stem of *Fitzroya cupressoides*, following the phylogenetic placement inferred by Mao et al.[97] for this fossil.

16. ***Fokienia-Chamaecyparis* crown**: min 61.1 Ma; mrca = *Fokienia hodginsii*, *Chamaecyparis lawsoniana*. The Paleocene fossil *Fokienia ravenscragensis*[132,133] was used to constrain the *Fokienia-Chamaecyparis* clade based on its inferred phylogenetic position from Mao et al.[97]

17. ***Juniperus* crown**: min 33.9 Ma; mrca = *Juniperus drupacea*, *Juniperus virginiana*. The oldest known juniper fossil, *Juniperus pauli*[134], was used as a minimum constraint on crown *Juniperus*, based on its inferred phylogenetic position in Mao et al.[97]

18. **Sequoioideae crown**: min 93.9 Ma; mrca = *Metasequoia glyptostroboides*, *Sequoiadendron giganteum*. Fossil evidence of *Metasequoia* dates back to the late Cretaceous[135], and its inclusion in previous phylogenetic analyses[97] supports its use as a minimum constraint for the small clade including *Metasequoia*, *Sequoia*, and *Sequoiadendron*.

19. **Taxodioideae crown**: min 113.0 Ma; mrca = *Cryptomeria japonica*, *Taxodium distichum*, *Glyptostrobus pensilis*. Fossil evidence of *Glyptostrobus*[136] dates to the late early Cretaceous (ca. 113 Ma), with phylogenetic analyses[97] supporting a placement within the clade including *Cryptomeria*, *Taxodium*, and *Glyptostrobus*. Therefore, we used this fossil evidence as a minimum age constraint for the clade.

20. ***Tetraclinis* stem**: min 32.2 Ma; mrca = *Tetraclinis articulata*, *Platycladus orientalis*, *Microbiota decussata*. Fossil evidence of *Tetraclinis* dates to the Oligocene, based on the taxon *Tetraclinis salicornioides*[137], with phylogenetic analyses[97] supporting its use as a stem representative of this genus. Thus, we used this fossil evidence to constrain the immediate broader clade including *Tetraclinis*.

21. ***Thuja* stem**: min 59.2 Ma; mrca = *Thuja occidentalis*, *Thujopsis dolabrata*. Fossil evidence of *Thuja* dates back to the Paleocene (*Thuja polaris*[138]), which we used as a minimum constraint on the divergence between *Thuja* and *Thujopsis*.

22. **Cycad stem**: min 298.9 Ma; mrca = *Zamia furfuracea*, *Ginkgo biloba*. The fossil record of Cycadales traces back to the late Pennsylvanian[139], with representative taxa including the fossil genus *Spermopteris*[140] and associated reproductive structures. Given that this fossil evidence likely represents early or stem representatives of the cycad lineage, we used the age of *Spermopteris* as a minimum constraint on the cycad + *Ginkgo* divergence.

23. **Cycad crown**: min 270.0 Ma; mrca = *Cycas circinalis*, *Zamia furfuracea*. There are two fossil genera that seem to represent early evidence of the Cycadales crown. These are *Crossozamia*[141,142] and *Primocycas*[143], both of which are ca. 270 Ma, and the former of which was placed phylogenetically by Hermsen et al.[144] We therefore used 270 Ma as a minimum age constraint for the cycad crown.

24. **Zamiaceae crown**: min 218 Ma; mrca = *Macrozamia elegans*, *Dioon edule*, *Bowenia spectabilis*, *Stangeria eriopus*, *Zamia furfuracea*. The fossil taxon *Lyssoxylon grigsbyi*[145] was used to calibrate the crown of Zamiaceae. *Lyssoxylon*, along with several other Triassic cycads, was shown to be nested within Zamiaceae based on the phylogenetic analyses of Hermsen et al.[144], thus justifying its use as a minimum age calibration for the crown of Zamiaceae.

25. **Gnetales stem**: min 265.1 Ma; mrca = *Gnetum gnemon*, *Pinus bungeana*. Polyplicate pollen (similar to that of *Ephedra* and *Welwitschia*) recovered from Permian sediments (268.8 to 265 Ma) likely represents the oldest fossil evidence of the Gnetales lineage[146], and therefore we used this evidence to constrain the Gnetales stem, i.e., the Gnetales + Pinaceae divergence.

26. **Gnetaceae + Welwitschiaceae crown**: min 114.0 Ma; mrca = *Gnetum gnemon*, *Welwitschia mirabilis*. A fossilized cotyledon, *Cratonia cotyledon*[147], which shows striking resemblance to *Welwitschia*, represents the oldest fossil evidence of the lineage leading to the extant monotypic genus *Welwitschia* and therefore can be used as a minimum age constraint for the *Gnetum* + *Welwitschia* clade.

27. ***Ephedra* crown**: min 125 Ma; mrca = *Ephedra breana*, *Ephedra frustillata*, *Ephedra ochreata*. The fossil taxon *Ephedra drewriensis*[148] represents seeds with in situ pollen showing strong morphological similarities to the living species comprising crown *Ephedra*, and therefore we used this fossil as a minimum constraint on the crown.

28. **Pinaceae crown**: min 151.1 Ma; mrca = *Pinus bungeana*, *Cedrus deodara*. The taxon *Eathiestrobus mackenziei* represents the oldest fossil evidence of Pinaceae[149] and was shown to represent a member of the crown clade in phylogenetic analyses by Gernandt et al.[102]

29. ***Keteleeria* stem**: min 120 Ma; mrca = *Keteleeria fortunei*, *Abies koreana*. The fossil *Pitystrobus corneti* (age from Klymiuk and Stockey[150]) was placed in recent phylogenetic

analyses[102] as sister to *Keteleeria* (i.e., on the *Keteleeria* stem), and therefore we used this fossil to calibrate the *Keteleeria* + *Abies* clade. However, given some uncertainty concerning the placement of this fossil, we also conducted an analysis without it to ensure that its inclusion was not overly influential in the main analysis.

30. ***Larix* crown**: min 41.2 Ma; mrca = *Larix gmelinii*, *Larix sibirica*. The fossil taxon *Larix altoborealis* was described by LePage and Basinger[151] as virtually identical to modern *Larix*, and therefore we used it as a minimum constraint for the crown clade.
31. ***Picea* stem**: min 132.9 Ma; mrca = *Pinus bungeana*, *Picea abies*. The taxon *Picea burtonii* represents the oldest fossil evidence of Picea[150], phylogenetically placed along the Picea stem[102,150], and therefore we use it as a minimum constraint for the *Picea* + *Pinus* divergence.
32. ***Pinus* subgenus *Pinus* crown**: min 129.0 Ma; mrca = *Pinus ponderosa*, *Pinus contorta*. This calibration is based on *Pinus yorkshirensis*, the oldest phylogenetically placed fossil representative of *Pinus* subgenus *Pinus*[102,150].
33. ***Pinus* subgenus *Strobus* crown**: min 125.0 Ma; mrca = *Pinus bungeana*, *Pinus nelsonii*. This calibration is based on *Pinus belgica,* the oldest phylogenetically placed fossil representative of *Pinus* subgenus *Strobus*[102,150].
34. ***Tsuga* stem**: min 41.2 Ma; mrca = *Tsuga canadensis*, *Nothotsuga longibracteata*. Cretaceous fossil pollen has been attributed *Tsuga* and used as a lower bound in previous dating analyses[21], but given the taxonomic ambiguity of the pollen evidence, we instead use the Eocene fossil taxon *Tsuga swedaea*[152], considered a stem representative, as a minimum constraint for the *Tsuga* + *Nothotsuga* divergence.
35. **Taxaceae stem**: min 197.0 Ma; mrca = *Taxus baccata*, *Torreya taxifolia*, *Cephalotaxus lanceolata*. This constraint is based on the fossil *Palaeotaxus redivia*[153], considered a stem member of Taxaceae in the traditional sense (i.e., *sensu stricto*, excluding *Cephalotaxus*). *Cephalotaxus* is generally included within Taxaceae today, but our results show that it is positioned sister to the remainder of the family (Taxaceae s.s.). We therefore used this fossil as a minimum constraint for Taxaceae s.l., i.e., for the divergence of *Cephalotaxus* and the rest of the family.

**List of traits scored for the phenotypic reconstructions**

Many of the traits examined here (or variants thereof) have been included in previous trait datasets, either for seed plants as a whole or for specific subclades of gymnosperms. In such cases, we cite the relevant dataset(s) and corresponding character number(s): Hilton and Bateman[96] (=HB2006); Friis et al.[108] (=FEA2007); Mao et al.[97] (=MEA2012); Escapa and Catalano[98] (=EC2013); Gernandt et al.[102] (=GEA2018); Coiro and Pott[99] (=CP2017); Hart[95] (=H1987); Herrera et al.[100] (=HEA2017); Herrera et al.[101] (=HEA2020); Schulz et al.[154] (=SEA2014); Herting et al.[155] (=JHEA2020); Andruchow-Colombo et al.[103] (=AEA-2019). Additional trait data were obtained from the primary literature[156–187].

1. **Life history**: homosporous (0), heterosporous (1). HB2006-1
2. **Secondary growth (wood production)**: present (0), absent (1). FEA2007-2, EC2013-48.
3. **Growth form**: tree (0), shrub (1), liana/woody climber (2), herb (3). MEA2012-1, HB2006-2, EC2013-48.
4. **Primary stem location**: Above ground (0); subterranean (1); aquatic/submerged (2).
5. **Sex distribution**: bisexual structures, functionally (0); bisexual structures, some vestigial (1); unisexual structures, monoecious (2); unisexual structures, dioecious (3). MEA2012-2, GEA2018-8, CP2017-57.
6. **Leaf, branch persistence**: evergreen (0); deciduous (1) branches; deciduous leaves (2). MEA2012-3, GEA2018-46.
7. **Short shoots**: absent (0); present (1). GEA2018-4, HB2006-5.
8. **Secondary growth**: absent (0); present (1).
9. **Vessels**: absent (0); present, "gnetalean" (1); present, "angiospermic" (2). HB2006-25, Thompson[156].
10. **Torus**: absent (0); present (1). HB2006-27, GEA2018-17.
11. **Crassulae**: absent (0); present (1). H1987-16.
12. **Resin ducts in secondary wood:** absent (0); present (1). H1987-17.
13. **Resin ducts in rays**: absent (0); present (1). H1987-19.
14. **Primary xylem**: mesarch (0); exarch (1); endarch (2). CP2017-9, HB2006-21.
15. **Wood parenchyma**: minimal (pycnoxylic) (0); abundant (manoxylic) (1).
16. **Girdling leaf traces**: absent (0); present (1). HB2006-14.

28. **Leaf lamina complexity**: Broad, flattened lamina (0); scale (1); awl (2); needle (3); broad, compound (4); broad; deeply lobed (5); bilaterally flattened (6); reduced (7). SEA2014.

29. **Leaf needle shape**: rounded (0); three-angled (1); four-angled (2).

55. **Number of microsporophylls/sporangiophores/stamens per strobilis**: under 10, definite number (0); 25 or under (1); 26 to 100 (2); more than 100 (3).

73. **Pollen aperture number**: absent/zero (0); one (1); more than three (2); three (3).

84. **Carpel arrangement**: spiral (0); whorled (1).

85. **Carpel closure**: by secretion, without postgenital fusion (0); with a continuous secretory canal but partial postgenital fusion at the periphery (1); with an incomplete secretory canal and complete postgenital fusion at the periphery (2); complete postgenital fusion (3). Seed-HB2006-39.

86. **Position of ultimate ovule-bearing structure** (e.g., cone or flower): lateral or axillary (0); terminal (1). HEA2020-29, GEA2018-102; H1987-98.

87. **Seed cones solitary or aggregated:** solitary (0); clustered/aggregated (1); borne on spike-like structure (2). MEA2012-25.

88. **Seed cone orientation at pollination time**: erect (0); horizontal/plagiotropic (1); downward/pendulous (2); irregular (3). GEA2018-105.

89. **Seed cone orientation at seed dispersal**: erect (0); horizontal/plagiotropic (1) downward/pendulous (2); irregular (3).

90. **Cone/seed dispersal at maturity**: cones open, seeds released (0); cones disintegrating, releasing seeds and scales (1); cones open, seeds released with part of supporting structure (2); cones closed, berry-like (3); cones reduced, drupe-like, with or without receptacle (4). EC2013-2, MEA2012-39, GEA2018-108, AEA2019-54, AEA2019-55, EC2013-19; GEA2018-108, AEA2019-53; HEA2017-15, GEA2018-107, EC2013-7, AEA2019-55; JHEA 2020.

91. **Seed wings**: absent (0); present (1).

92. **Seed wing derivation**: derived from seed integument (0); derived from scale tissue (1); scales functioning as wings (seeds and scales entirely fused) (2); derived from bract (3). AEA2019-56, EC2013-20, MEA2012-48, GEA2018-181, GEA2018-180, H1987-119, GEA2018-123; AEA2019-45, EC2013-17.

93. **Seed wing number**: one encircling (0); one (1); two (2); three (3). AEA2019-57, EC2013-21, MEA2012-49.

94. **Seed wing symmetry** (for single-wing seeds): symmetric (0); asymmetric (1). AEA2019-58, EC2013-22, MEA2012-51.

95. **Seed wing symmetry** (for two winged seeds): symmetric (0); asymmetric (1).

96. **Seed wing position**: lateral/marginal (0); encircling (1); terminal (2). HEA2017-22, MEA2012-50.

97. **Seed cone constitution**: all components sclerified (0); not all components sclerified (1). JHEA2020.

98. **Cone persistence:** cone caudućous (0); cone disintegrating (1); cone persistent (2); cone persistent until fire (serotinous) (3). MEA2012-24, GEA2018-106.

99. **Seed maturation/reproductive cycle**: one (0); two (1); three (2). MEA2012-46.

100. **Main cone scales**: bract scales (0); seed scales (1); bract-seed scale complexes (2); reduced and modified/fleshy (3).

101. **Fertile and sterile scales**: all fertile (0); fertile in middle (sterile scales proximally and distally) (1); sterile scales proximally (fertile distally) (2); sterile scales distally (fertile proximally) (3); sterile scales proximally (fertile distally, plus ovules on terminal on axis) (4); usually only one (or two) seeds on terminal fertile scale (5).

102. **Bract scale morph**: reduced (0); fused with scale (1); flattened, broad (2); peltate (3); becoming fleshy (4). GEA2018-115, HEA2020-33.

103. **Bract scale apex morphology**: apical protruding spine/point (0); central point (displaced from apex) (1); three pointed/lobed (2); numerous teeth and point (3); rounded (spine or point lacking) (4); distinctly truncate (no spine) (5); fork with prominent middle spine/vein (6).

104. **Bract-seed scale relative length when separate** (not fused): bracts shorter (0); roughly equivalent with seed scales (1); emergent (2). AEA2019-43, EC2013-11.

105. **Seed scale apex with distinct spiny projection**: absent (0); simple point (1); apophysis and umbo (2). GEA2018-122, GEA2018-128, GEA2018-126, GEA2018-127.

106. **Bract/scale fusion**: entirely fused (0); free (1); fused (seed scale highly reduced) (2); fused (seed scale absent or not visible) (3). EC2013-9, AEA2019-41, AEA2019-42, EC2013-10, MEA2012-40, GEA2018-111, HEA2020-32, HB2006-55, MEA2012-44, H1987-101.

107. **Cone scale attachment**: spiral (0); decussate (1); whorled (2). AEA2019-36, EC2013-5, MEA2012-26, HB2006-40, 54.

108. **Cone scale arrangement**: imbricate (0); valvate (1).

109. **Number of scales per cone**: few (0); four to ten (1); eleven to 50 (2); more than 50 (3).

110. **Ovule number per ovuliferous scale/megasporophyll**: more than three (0); one (1); two (2); three (3). HEA2017-19, GEA2018-172, HEA2020-41, AEA2019-44, EC2013-16, MEA2012-47, CP2017-79.

111. **Ovule position on bract/scale or supporting structure**: terminal (0); perisporangiate (peltate) (1); marginal (2); in axil of bract (3); pseudoterminal (4); on adaxial scale surface (5); abaxially on basal part of cone scale (6). CP2017-78, GEA2018-169, MEA2012-33, EC2013-23, HEA2017-21, HEA2020-39, 40, HB2006-36, AEA2019-37.

112. **Ovule on axillar pre-developed structure**: absent (0); present (1). JHEA2020.

113. **Ovule orientation**: erect (0); inverted (1). MEA2012-32, GEA2018-171, HEA2020-36, EC2013-18, AEA2019-52, HEA2017-20, HB2006-37, H1987-114, JHEA2020.

114. **Ovule apex/micropyle**: simple apex (0); straight tubular micropyle (1); bifid (2). HEA2020-48, HB2006-63, CP2017-84, GEA2018-162.

115. **Micropyle position at pollination**: pointing to cone axis/base at pollination (0); pointing to cone periphery/apex (1). JHEA2020.

116. **Change in micropyle orientation**: orientation changes before pollination (0); after pollination (1); does not change (2). JHEA2020.

127. **Fleshy receptacle**: absent (0), present (1).

#### 2. Supplementary Tables

**Supplementary Table S1**. Transcriptomes and genomes sequenced and sampled. Newly generated transcriptomic data are available <here>. See the 1KP website (http://www.onekp.com/samples/list.php) for further information on the 1KP sampled included in the study. Herbarium codes (following Index Herbariorum: http://sweetgum.nybg.org/science/ih/) are presented in parentheses where relevant. Botanical garden acronyms are as follows: UCBG: University of California Botanical Garden at Berkeley; KBG: Kunming Botanical Garden, Chinese Academy of Sciences; MBG: Missouri Botanical Garden; AAHU: The Arnold Arboretum of Harvard University; RBGE: Royal Botanic Garden Edinburgh; Kew: Royal Botanic Garden Kew.

| **Taxon** | **Data deposition or source** | **Voucher info** | **Cluster grouping** |
|---|---|---|---|
| *Equisetum_hyemale* | 1KP: JVSZ | Rothfels 4137 (DUKE) | Ferns-Lycos |
| *Selaginella_apoda* | 1KP: LGDQ | Rothfels 4118 (DUKE) | Ferns-Lycos |
| *Isoetes_tegetiformans* | 1KP: PKOX | JLM2013-47 (GA) | Ferns-Lycos |
| *Psilotum_nudum* | 1KP: QVMR | See 1KP website | Ferns-Lycos |
| *Marattia_attenuata* | 1KP: UGNK | NYBG 1295/78-A | Ferns-Lycos |
| *Ophioglossum_petiolatum* | 1KP: WTJG | 132699 (ALTA) | Ferns-Lycos |
| *Huperzia_selago* | 1KP: YHZW | A. Larsson 80 (UPS) or AL80 (UPS) | Ferns-Lycos |
| *Nuphar_advena* | 1KP: WTKZ | Soltis and Miles 2783 | Angiosperms |
| *Arabidopsis_thaliana* | Phytozome: 447_Araport11 | - | Angiosperms |
| *Daucus_carota* | Phytozome: 388_v2.0 | - | Angiosperms |
| *Austrobaileya_scandens* | 1KP: FZJL | JLM2013-22 (GA) | Angiosperms |
| *Malus_domestica* | Phytozome: 491_v1.1 | - | Angiosperms |
| *Acorus_americanus* | 1KP: MTII | M.K. Deyholos 2016-121 | Angiosperms |
| *Kadsura_heteroclita* | 1KP: NWMY | T. Chen et. al. 081210-04 | Angiosperms |
| *Sarcandra_glabra* | 1KP: OSHQ | T. Chen et. al. 081210-04 | Angiosperms |
| *Nymphaea_sp* | 1KP: PZRT | T. Chen et al. 2010090805 | Angiosperms |
| *Papaver_somniferum* | 1KP: RQNK | T. Kutchan 6173586 (MO) | Angiosperms |
| *Solanum_tuberosum* | Phytozome: 448_v4.03 | - | Angiosperms |
| *Amborella_trichopoda* | 1KP: URDJ | Soltis and Miles 2778 | Angiosperms |
| *Illicium_floridanum* | 1KP: VZCI | Soltis and Miles 2960 | Angiosperms |
| *Magnolia_grandiflora* | 1KP: WBOD | Soltis and Miles 2764 | Angiosperms |
| *Persea_borbonia* | 1KP: WIGA | Soltis and Miles 2980 | Angiosperms |
| *Zea_mays* | Phytozome:PH207_443_v1.1 | - | Angiosperms |
| *Bowenia_serrulata* | SRA: SRR14381590 (new) | 2008.0429 in UCBG | Cycads-Ginkgo |
| *Ceratozamia_hildae* | SRA: SRR14381651 (new) | 2005.0703 in UCBG | Cycads-Ginkgo |
| *Cycas_rumphii* | SRA: SRR14381646 (new) | 1984-0207-1 in MBG | Cycads-Ginkgo |
| *Dioon_spinulosum* | SRA: SRR14381642 (new) | 2004.0995 in UCBG | Cycads-Ginkgo |
| *Encephalartos_hirsutus* | SRA: SRR14381640 (new) | 2005.0167 in UCBG | Cycads-Ginkgo |
| *Ginkgo_biloba* | SRA: SRR14381635 (new) | Yi19048 in KBG | Cycads-Ginkgo |
| *Encephalartos_barteri* | 1KP: GNQG | See 1KP website | Cycads-Ginkgo |
| *Lepidozamia_peroffskyana* | SRA: SRR14381625 (new) | 2004.0927 in UCBG | Cycads-Ginkgo |
| *Microcycas_calocoma* | SRA: SRR14381618 (new) | 2005.0695 in UCBG | Cycads-Ginkgo |
| *Macrozamia_miguelii* | SRA: SRR14381622 (new) | 2004.0950 in UCBG | Cycads-Ginkgo |
| *Stangeria_eriopus* | SRA: SRR14381598 (new) | 2005.1126 in UCBG | Cycads-Ginkgo |
| *Dioon_edule* | 1KP: WLIC | 1576/87-G; NYBG | Cycads-Ginkgo |
| *Cycas_micholitzii* | 1KP: XZUY | See 1KP website | Cycads-Ginkgo |
| *Zamia_furfuracea* | SRA: SRR14381583 (new) | 95.0069 in UCBG | Cycads-Ginkgo |

| | | | |
|---|---|---|---|
| *Gnetum_montanum* | 1KP: GTHK | See 1KP website | Gnetales |
| *Ephedra_equisetina* | SRA: SRR14381639 (new) | 2015-0256-2 in MBG | Gnetales |
| *Gnetum_parvifolium* | SRA: SRR14381632 (new) | 1993-0791-2 in MBG | Gnetales |
| *Ephedra_sinica* | 1KP: VDAO | 132703 (ALTA) | Gnetales |
| *Welwitschia_mirabilis* | SRA: SRR14381587 (new) | 2012.0748 in UCBG | Gnetales |
| *Tsuga_forrestii* | SRA: SRR14381588 (new) | Yi19049 in KBG | Pinaceae |
| *Larix_speciosa* | 1KP: WVWN | R. Mindell & B.C. Zhuang 36696 (UBC) | Pinaceae |
| *Abies_koreana* | SRA: SRR14381657 (new) | 93.065 in UCBG | Pinaceae |
| *Nothotsuga_longibracteata* | 1KP: AREG | JLM2013-51 (GA) | Pinaceae |
| *Picea_engelmannii* | 1KP: AWQB | 80226B | Pinaceae |
| *Cathaya_argyrophylla* | SRA: SRR14381654 (new) | 2000.0274 in UCBG | Pinaceae |
| *Cedrus_deodara* | SRA: SRR14381653 (new) | 45.0273 in UCBG | Pinaceae |
| *Pinus_radiata* | 1KP: DZQM | P. Thomas 20090072 (E) | Pinaceae |
| *Tsuga_heterophylla* | 1KP: GAMH | 6182009A | Pinaceae |
| *Cedrus_libani* | 1KP: GGEA | B.C. Zhuang & C. Gallant bg102/7617 (UBC) | Pinaceae |
| *Pinus_parviflora* | 1KP: IIOL | P. Thomas 19721337 (E) | Pinaceae |
| *Pseudotsuga_wilsoniana* | 1KP: IOVS | R. Mindell & B.C. Zhuang 31183 (UBC) | Pinaceae |
| *Pinus_ponderosa* | 1KP: JBND | Stewart and Burris | Pinaceae |
| *Keteleeria_evelyniana* | 1KP: JUWL | JLM2013-48 (GA) | Pinaceae |
| *Keteleeria_calcarea* | SRA: SRR14381628 (new) | Yi19050 in KBG | Pinaceae |
| *Larix_decidua* | SRA: SRR14381626 (new) | 804-33*A in AAHU | Pinaceae |
| *Pinus_jeffreyi* | 1KP: MFTM | Stewart and Burris | Pinaceae |
| *Picea_abies* | SRA: SRR14381614 (new) | 1053-24*A in AAHU | Pinaceae |
| *Pseudolarix_amabilis* | SRA: SRR14381607 (new) | 2001.0806 in UCBG | Pinaceae |
| *Pinus_bungeana* | SRA: SRR14381611 (new) | Yi19051 in KBG | Pinaceae |
| *Pseudotsuga_sinensis* | SRA: SRR14381605 (new) | Yi19052 in KBG | Pinaceae |
| *Abies_lasiocarpa* | 1KP: VSRH | 2036/2007A | Pinaceae |
| *Nageia_nagi* | SRA: SRR14381617 (new) | Yi19053 in KBG | Podocarpaceae |
| *Afrocarpus_falcatus* | SRA: SRR14381656 (new) | 1990-3396-1 in MBG | Podocarpaceae |
| *Lepidothamnus_sp* | 1KP: BBDD | B.C. Zhuang & C. Gallant bg049/25451 (UBC) | Podocarpaceae |
| *Dacrydium_cupressinum* | SRA: SRR14381643 (new) | 56.0445 in UCBG | Podocarpaceae |
| *Dacrycarpus_dacrydioides* | SRA: SRR14381644 (new) | 61.1084 in UCBG | Podocarpaceae |
| *Dacrycarpus_compactus* | 1KP: FMWZ | P. Thomas 19643273 (E) | Podocarpaceae |
| *Falcatifolium_taxoides* | SRA: SRR14381638 (new) | 19842581 in RBGE | Podocarpaceae |
| *Halocarpus_bidwillii* | SRA: SRR14381631 (new) | 19832579 in RBGE | Podocarpaceae |
| *Acmopyle_pancheri* | 1KP: HILW | JLM2013-15 (GA) | Podocarpaceae |
| *Dacrydium_balansae* | 1KP: IZGN | JLM2013-34 (GA) | Podocarpaceae |
| *Phyllocladus_hypophyllus* | 1KP: JRNA | P. Thomas 19781711 (E) | Podocarpaceae |
| *Parasitaxus_usta* | 1KP: JZVE | See 1KP website | Podocarpaceae |
| *Lagarostrobos_franklinii* | SRA: SRR14381627 (new) | 19588943 in RBGE | Podocarpaceae |
| *Manoao_colensoi* | SRA: SRR14381621 (new) | 19842513 in RBGE | Podocarpaceae |
| *Microcachrys_tetragona* | 1KP: MHGD | B.C. Zhuang & C. Gallant bg048/38870 (UBC) | Podocarpaceae |
| *Prumnopitys_andina* | SRA: SRR14381608 (new) | 94.0787 in UCBG | Podocarpaceae |
| *Podocarpus_macrophyllus* | SRA: SRR14381609 (new) | Yi19054 in KBG | Podocarpaceae |
| *Phyllocladus_trichomanoides* | SRA: SRR14381615 (new) | 92.0331 in UCBG | Podocarpaceae |
| *Retrophyllum_rospigliosii* | SRA: SRR14381604 (new) | 2008.0433 in UCBG | Podocarpaceae |

| | | | |
|---|---|---|---|
| *Sundacarpus_amarus* | SRA: SRR14381597 (new) | 20030752A in RBGE | Podocarpaceae |
| *Saxegothaea_conspicua* | SRA: SRR14381603 (new) | 61.1088 in UCBG | Podocarpaceae |
| *Retrophyllum_minus* | 1KP: VGSX | JLM2013-62 (GA) | Podocarpaceae |
| *Podocarpus_rubens* | 1KP: XLGK | P. Thomas 20000597*A4 (E) | Podocarpaceae |
| *Agathis_australis* | SRA: SRR14381645 (new) | 73.0561 in UCBG | Araucariaceae |
| *Agathis_macrophylla* | 1KP: ACWS | 88/53-A; (MBC) | Araucariaceae |
| *Araucaria_montana* | SRA: SRR14381623 (new) | 2011-2071-1 in MBG | Araucariaceae |
| *Agathis_robusta* | 1KP: MIXZ | 87/98-A; (MBC) | Araucariaceae |
| *Wollemia_nobilis* | SRA: SRR14381585 (new) | 2006.0775 in UCBG | Araucariaceae |
| *Araucaria_rulei* | 1KP: XTZO | JLM2013-19 (GA) | Araucariaceae |
| *Taxus_baccata* | 1KP: WWSS | 132705 (ALTA) | Sciadopitys-Taxaceae |
| *Amentotaxus_formosana* | SRA: SRR14381634 (new) | 2001.0542 in UCBG | Sciadopitys-Taxaceae |
| *Austrotaxus_spicata* | 1KP: BTTS | See 1KP website | Sciadopitys-Taxaceae |
| *Cephalotaxus_fortunei* | SRA: SRR14381652 (new) | Yi19055 in KBG | Sciadopitys-Taxaceae |
| *Torreya_taxifolia* | 1KP: EFMS | JLM2013-73 (GA) | Sciadopitys-Taxaceae |
| *Torreya_nucifera* | 1KP: HQOM | B.C. Zhuang & C. Gallant bg106/6921 (UBC) | Sciadopitys-Taxaceae |
| *Amentotaxus_argotaenia* | 1KP: IAJW | P. Thomas 19763949 (E) | Sciadopitys-Taxaceae |
| *Pseudotaxus_chienii* | SRA: SRR14381606 (new) | 34-2000*D in AAHU | Sciadopitys-Taxaceae |
| *Sciadopitys_verticillata* | SRA: SRR14381602 (new) | 88.0163 in UCBG | Sciadopitys-Taxaceae |
| *Torreya_californica* | SRA: SRR14381589 (new) | 71.0599 in UCBG | Sciadopitys-Taxaceae |
| *Taxus_yunnanensis* | SRA: SRR14381594 (new) | Yi19056 in KBG | Sciadopitys-Taxaceae |
| *Taxus_cuspidata* | 1KP: ZYAX | 132706 (ALTA) | Sciadopitys-Taxaceae |
| *Callitropsis_nootkatensis* | SRA: SRR14381581 (new) | 19687302A in RBGE | Cupressaceae-1 |
| *Calocedrus_decurrens* | 1KP: FRPM | B.C. Zhuang & C. Gallant bg422/40057 (UBC) | Cupressaceae-1 |
| *Calocedrus_macrolepis* | SRA: SRR14381655 (new) | Yi19057 in KBG | Cupressaceae-1 |
| *Chamaecyparis_lawsoniana* | SRA: SRR14381650 (new) | Yi19058 in KBG | Cupressaceae-1 |
| *Cupressus_sempervirens* | SRA: SRR14381647 (new) | 2004-1437-1 in MBG | Cupressaceae-1 |
| *Fokienia_hodginsii* | SRA: SRR14381636 (new) | Yi19059 in KBG | Cupressaceae-1 |
| *Hesperocyparis_glabra* | SRA: SRR14381630 (new) | Yi19060 in KBG | Cupressaceae-1 |
| *Juniperus_formosana* | SRA: SRR14381629 (new) | Yi19061 in KBG | Cupressaceae-1 |
| *Juniperus_scopulorum* | 1KP: XMGP | E. La Fountaine 2013-002 (UBC) | Cupressaceae-1 |
| *Microbiota_decussata* | SRA: SRR14381619 (new) | 180-2002*F in in AAHU | Cupressaceae-1 |
| *Platycladus_orientalis* | SRA: SRR14381610 (new) | Yi19062 in KBG | Cupressaceae-1 |
| *Tetraclinis_articulata* | SRA: SRR14381593 (new) | 88.1295 in UCBG | Cupressaceae-1 |
| *Tetraclinis_sp* | 1KP: CGDN | JLM2013-72 (GA) | Cupressaceae-1 |

| | | | |
|---|---|---|---|
| *Thuja_plicata* | 1KP: VFYZ | B.C. Zhuang & C. Gallant bg426/39575 (UBC) | Cupressaceae-1 |
| *Thuja_standishii* | SRA: SRR14381592 (new) | Yi19063 in KBG | Cupressaceae-1 |
| *Thujopsis_dolabrata* | SRA: SRR14381591 (new) | Yi19064 in KBG | Cupressaceae-1 |
| *Xanthocyparis_vietnamensis* | SRA: SRR14381584 (new) | Yi19065 in KBG | Cupressaceae-1 |
| *Austrocedrus_chilensis* | SRA: SRR14381601 (new) | 2010.0382 in UCBG | Cupressaceae-2 |
| *Callitris_endlicheri* | SRA: SRR14381582 (new) | 87.0118 in UCBG | Cupressaceae-2 |
| *Callitris_gracilis* | 1KP: IFLI | DJ533A | Cupressaceae-2 |
| *Diselma_archeri* | SRA: SRR14381641 (new) | 65.017 in UCBG | Cupressaceae-2 |
| *Fitzroya_cupressoides* | SRA: SRR14381637 (new) | 2007.0165 in UCBG | Cupressaceae-2 |
| *Libocedrus_plumosa* | SRA: SRR14381624 (new) | 90.0647 in UCBG | Cupressaceae-2 |
| *Neocallitropsis_pancheri* | 1KP: JDQB | See 1KP website | Cupressaceae-2 |
| *Papuacedrus_papuana* | SRA: SRR14381616 (new) | 19682163C1 in RBGE | Cupressaceae-2 |
| *Pilgerodendron_uviferum* | SRA: SRR14381613 (new) | 2001.0044 in UCBG | Cupressaceae-2 |
| *Widdringtonia_cedarbergensis* | 1KP: AUDE | L. DeGironimo 637/2004-B (NY) | Cupressaceae-2 |
| *Widdringtonia_nodiflora* | SRA: SRR14381586 (new) | 2015.0313 in UCBG | Cupressaceae-2 |
| *Athrotaxis_cupressoides* | 1KP: XIRK | P. Thomas 20052511 (E) | Cupressaceae-3 |
| *Athrotaxis_laxifolia* | SRA: SRR14381612 (new) | 2012.0753 in UCBG | Cupressaceae-3 |
| *Cryptomeria_japonica* | SRA: SRR14381649 (new) | Yi19066 in KBG | Cupressaceae-3 |
| *Cunninghamia_lanceolata* | SRA: SRR14381648 (new) | Yi19067 in KBG | Cupressaceae-3 |
| *Glyptostrobus_pensilis* | SRA: SRR14381633 (new) | Yi19068 in KBG | Cupressaceae-3 |
| *Metasequoia_glyptostroboides* | SRA: SRR14381620 (new) | Yi19069 in KBG | Cupressaceae-3 |
| *Sequoia_sempervirens* | SRA: SRR14381600 (new) | Yi19070 in KBG | Cupressaceae-3 |
| *Sequoiadendron_giganteum* | SRA: SRR14381599 (new) | 2002.1062 in UCBG | Cupressaceae-3 |
| *Taiwania_cryptomerioides* | SRA: SRR14381596 (new) | Yi19071 in KBG | Cupressaceae-3 |
| *Taxodium_ascendens* | SRA: SRR14381595 (new) | Yi19072 in KBG | Cupressaceae-3 |
| *Taxodium_distichum* | 1KP: FHST | B.C. Zhuang & C. Gallant bg104/27812 (UBC) | Cupressaceae-3 |

**Supplementary Table S2**. Newly sequenced plastid genomes. The key to botanical garden acronyms is presented in the caption for Supplementary Table 1.

| Taxon | Voucher info | NCBI accession number | SRA accession number (reads) |
|---|---|---|---|
| *Acmopyle pancheri* | M803 in Kew (DNA) | MW470971 | SRR14374001 |
| *Actinostrobus pyramidalis* | 19923616AB in Kew (fresh) | MW470972 | SRR14374000 |
| *Afrocarpus falcatus* | 1990-3396-1 in MBG (fresh) | MW470973 | SRR14373989 |
| *Athrotaxis laxifolia* | 2012.0753 in UCBG (fresh) | MW470974 | SRR14373978 |

| | | | |
|---|---|---|---|
| *Austrocedrus chilensis* | 2010.0382 in UCBG (fresh) | MW470975 | SRR14373975 |
| *Austrotaxus spicata* | M37741 in Kew (DNA) | MW470976 | SRR14373974 |
| *Calocedrus decurrens* | Yi14152-28 in KBG (specimen) | MK241803 | SRR14373973 |
| *Calocedrus formosana* | Yi16045 in KBG (specimen) | KX832620 | SRR14373972 |
| *Calocedrus rupestris* | Yi13630 in KBG (specimen) | MK225565 | SRR14373971 |
| *Dacrydium cupressinum* | 56.0445 in UCBG (fresh) | MW470977 | SRR14373970 |
| *Diselma archeri* | 65.017 in UCBG (fresh) | MW470978 | SRR14373999 |
| *Falcatifolium taxoides* | S10378 in Kew (specimen) | MW470979 | SRR14373998 |
| *Fitzroya cupressoides* | 2007.0165 in UCBG (fresh) | MW470980 | SRR14373997 |
| *Glyptostrobus pensilis* | Yi19068 in KBG (fresh) | MW470981 | SRR14373996 |
| *Halocarpus bidwillii* | 19832579 in RBGE (fresh) | MW470982 | SRR14373995 |
| *Lagarostrobos franklinii* | 19588943 in RBGE (fresh) | MW470983 | SRR14373994 |
| *Libocedrus plumosa* | 90.0647 in UCBG (fresh) | MW470984 | SRR14373993 |
| *Manoao colensoi* | 19842513 in RBGE (fresh) | MN016935 | SRR14373992 |
| *Microbiota decussata* | 180-2002*F in in AAHU (fresh) | MW470985 | SRR14373991 |
| *Papuacedrus papuana* | 19682163C1 in RBGE (fresh) | MW470986 | SRR14373990 |
| *Parasitaxus usta* | M37534 in Kew (DNA) | MN016936 | SRR14373988 |
| *Pherosphaera fitzgeraldii* | M804 in Kew (DNA) | MW470987 | SRR14373987 |
| *Phyllocladus trichomanoides* | 92.0331 in UCBG (fresh) | MW470988 | SRR14373986 |
| *Pilgerodendron uviferum* | 2001.0044 in UCBG (fresh) | MW470989 | SRR14373985 |
| *Prumnopitys andina* | 94.0787 in UCBG (fresh) | MW470990 | SRR14373984 |
| *Pseudotaxus chienii* | 34-2000*D in AAHU (fresh) | MW470991 | SRR14373983 |
| *Saxegothaea conspicua* | 61.1088 in UCBG (fresh) | MW470992 | SRR14373982 |
| *Sequoiadendron giganteum* | 2002.1062 in UCBG (fresh) | MW470993 | SRR14373981 |
| *Sundacarpus amarus* | S10385 in Kew (specimen) | MW470994 | SRR14373980 |
| *Taxodium distichum* | Yi14492 in KBG (fresh) | MW470995 | SRR14373979 |
| *Tetraclinis articulata* | 88.1295 in UCBG (fresh) | MW470996 | SRR14373977 |
| *Widdringtonia nodiflora* | 2015.0313 in UCBG (fresh) | MW470997 | SRR14373976 |

504

### 3. Supplementary Results and Discussion

**Phylogenetic relationships and phylogenomic conflict**

Inferring relationships among major gymnosperm lineages, and particularly the placements of *Ginkgo* and Gnetales, has proven to be one of the biggest challenges in plant phylogenetics[188]. Our results placed, with maximal support, *Ginkgo* sister to cycads, and Gnetales within conifers sister to Pinaceae, regardless of analysis type (Fig. 1 and Supplementary Figs. 16 and 17). Recently, Gnetales were placed sister to all conifers in a large phylogenomic analysis of green plant transcriptomes[32]. However, our analyses, based on an expanded sampling of gymnosperm taxa, show strong concordance among gene trees with our inferred placement of Gnetales sister to Pinaceae (Fig. 1), consistent with several other recent nuclear phylogenomic studies[38,105,189]. Ran et al.[38] and Smith et al.[105] both used various approaches to dissect the phylogenetic signal in their datasets and found strong support for the major relationships also inferred here. The study by Leebens-Mack et al.[32] was a broader investigation of green plant phylogenomics and also included a smaller number of genes in the analyses, and thus their inferred topology for gymnosperms might be an artifact of the complexities of phylogenetic analysis across such a large and heterogeneous clade. Focused analyses on gymnosperms[38] (and here) show that the nuclear genome predominantly supports the major relationships inferred here.

However, conflict/concordance analysis revealed that there were nevertheless notable levels of gene-tree conflict underlying the placements of Gnetales and *Ginkgo*, and particularly the latter taxon (Fig. 1). The placement of *Ginkgo* sister to cycads was supported by 310 gene trees (with > 70 percent bootstrap support), while 92 gene trees conflicted with this topology, with 81 of these supporting the alternative topology of *Ginkgo* sister to *Coniferae*. As noted in the text, morphological phylogenetic analyses[14] have often placed Ginkgoales with conifers (extinct and extant), rather than cycads. These results suggest the possibility of reticulation in the early evolution of gymnosperms, but we emphasize that this possibility requires further study—perhaps involving synteny analysis of complete genomes. Inferring ancient reticulation is a major challenge[190,191], and other processes might be responsible for the pattern of alternative gene trees observed here as well as for the shared morphological features.

**Polyploidy and genome evolution**

Inferring ancient WGDs is a major challenge given that diploidization can mask WGD events and changes in genome size and chromosome number can occur in unpredictable ways following polyploidy[192,193], and often occur without WGD[194,195]. To circumvent the limitations of particular datatypes or methods, we employed a multifaceted approach (Methods), finding that major genomic changes—including WGD, changes in chromosome number, and/or abrupt jumps or rate shifts in genome size—underlie the origins of most major clades and families of gymnosperms (see also Leitch et al.[196] and Burleigh et al.[197]). While polyploidy may occur at a lower frequency in gymnosperms compared to ferns and angiosperms[1,27,198], the notion that polyploidy is rare or insignificant in this group seems untenable in light of recent studies documenting prevalent neopolyploidy[24,25] and our results here documenting several possible ancient WGD events (see also Li et al.[22] and Leebens-Mack et al.[32]). Thus, polyploidy appears to be a prevalent phenomenon across all major extant euphyllophyte lineages.

Our results regarding WGD inference differ in several respects from previous studies[22,29,32,59]. This is likely due to a combination of differences in sampling and method of WGD inference. The studies by Jiao et al.[29], Li et al.[22], Roodt et al.[30], and Zwaenepoel and Van de Peer[59] examined a relatively small number of gymnosperm samples. If descendants of a WGD event show heterogeneous levels of signal, sampling one or two representatives could result in events going undetected, depending on the method(s) employed. Additionally, limited sampling can result in ambiguity with respect to the precise placement of detected WGDs. Both the studies by Li et al.[22] and Leebens-Mack et al.[32] also used a different tool—i.e., the multi-taxon polyploidy search (MAPS) tool—for mapping gene duplications and locating putative WGD events. Unlike our method of duplicate gene mapping, which maps gene duplications at all nodes in a tree of interest, MAPS is used to examine specific nodes of interest, and as a result, possible WGD events could be missed unless all nodes are specifically examined. The necessity of breaking up the total sampling into subsets (i.e., step-wise guide trees for mapping) can also make it difficult to reconcile results across the independent analyses, especially when partially overlapping subsets seem to show different results[196]. Finally, our species tree for gene-duplication mapping (which included Gnetales sister to Pinaceae) differed from the species tree used in Li et al.[22] and Leebens-Mack et al.[32], both of which placed Gnetales sister to conifers in the traditional sense. This might also lead to differences in gene-duplication mapping and WGD inference.

Regarding particular inferences, a gymnosperm-specific WGD event was not detected in the studies of Jiao et al.[29], Li et al.[22], or Leebens-Mack et al.[32], but here we found a high percentage of gene duplications unique to the gymnosperm branch, suggestive of a WGD event, consistent with tentative results from Zwaenepoel and Van de Peer[59]. Roodt et al.[30] highlighted *Ks* peaks shared between the cycad *Encephalartos* and *Ginkgo* and noted that these might represent an event unique the cycad-Ginkgo clade, or perhaps a deeper WGD for gymnosperms or seed plants, but not including sampling beyond 'ginkads' to evaluate these alternatives. Our results suggest that the *Ks* peaks observed by Roodt et al.[30] represent an ancestral gymnosperm WGD event. A conifer-specific WGD was also not detected in previous studies, but we find high levels of gene duplication in support of a possible conifer-specific WGD event. Consistent with Leebens-Mack et al.[32], we found elevated percentages of gene duplication in the branch subtending Cupressophytes, but no *Ks* peak. A Pinaceae-specific WGD event has been disputed in several studies[59,201], but we found that both gene-duplicaition mapping and *Ks* plots show signatures of a Pinaceae-specific WGD event (Fig. 1; data deposit), consistent with Li et al.[22] The discrepancies among these studies underscore the inherent challenges of ancient WGD inference[196], and our study in particular underscores the importance of increased sampling and the use of multiple, complementary methods for WGD inference. Ultimately, however, analyses of complete genomes will provide the most definitive evidence of the extent and locations of ancient WGD in gymnosperms[202].

Gymnosperms (especially conifers) are remarkable for having large genomes relative to most other major green plant lineages[41]. Transposable element (TE) proliferation seems to be the primary driver of extreme (large) genome sizes in gymnosperms [33,201,203], but WGD may have played an underappreciated role in genome size evolution in gymnosperms[22]—perhaps both in size increases (e.g., Pinaceae) and in triggering size decreases (as in Podocarpaceae). With increased sequencing efforts[33,204], we are also gaining a greater appreciation of the heterogeneity in genomic characteristics across gymnosperms (e.g., *Gnetum* shows elevated levels of retrotransposon elimination rather than accumulation[204]). Generalities about genome evolution across gymnosperms—to a large extent based on the remarkably large genomes of *Picea abies* and other Pinaceae (e.g., Nystedt et al.[201]), and under the assumption of no polyploidy and stable chromosome numbers—therefore are in need of reconsideration or at least refinement in light of our results and other recent studies[22,204].

**Diversification shifts and patterns**

As noted in the main text, diversification shifts in gymnosperms appear more closely connected to rate shifts in climatic evolution, with 14/17 of the inferred diversification shifts occurring on either the same branch as a climatic shift or on a branch only one or two nodes away from a climatic shift. However, there were several diversification shifts that coincided with genomic changes. Putative WGDs and diversification shifts coincided on the branches subtending *Cupressophyta* and Pinaceae. Concerted rate shifts in diversification and genome size (C-value) evolution were detected in the branches subtending *Pinus* subsection *Ponderosae*, *Pinus* subgenus *Strobus*, and an *Ephedra* subclade; a C-value rate shift in *Juniperus* was also one node away from a diversification shift (Extended Data Fig. 3). Concerted C-value jumps and diversification shifts were detected in the branches subtending *Picea*, *Gnetum*, *Ceratozamia*, and *Dioon*, with *Cycas* showing a C-value jump one node removed from a diversification shift (Extended Data Fig. 3).

Our results show that a considerable portion of gymnosperm species diversity is a product of recent Cenozoic radiations, consistent with several previous studies[20,21], and as we also show here that these diversification shifts appear connected to increased rates of climatic evolution particularly in more cool/arid conditions, which became more prevalent around the globe after the Eocene and after the mid-Miocene[46]. We argue that these patterns generally make sense in light of the competitive dynamics stemming from the basic functional morphology of gymnosperms vs. angiosperms, which would in many cases restrict gymnosperms to more stressful growing conditions[44].

Of course, gymnosperms have experienced major extinctions throughout their long history[205–207], and the Cretaceous and early Cenozoic rise of angiosperms likely drove the decline or extinction of various gymnosperm lineages[44]. But clearly there is heterogeneity in patterns of extinction across gymnosperms both phylogenetically[206] and spatially[21]. For example, cycads and gnetophytes showed more pronounced declines throughout the Cretaceous, while conifer diversity and abundance appeared relatively stable[208]. Unfortunately, however, quantitative examinations of gymnosperm diversity and abundance (in comparison to that of angiosperms), based on the entirety of the known seed plant fossil record, have not been undertaken across both the Cretaceous and Cenozoic (but see below, with respect to comments on Condamine et al.[54]). With respect to conifers, it has been suggested that the K-Pg mass extinction event played a large role in their displacement (as the dominant canopy trees) in many plant communities[47], and this possibility clearly deserves closer consideration based on the fossil record.

Accurately inferring diversification dynamics from phylogenies of only extant taxa is notoriously difficult, if not impossible, especially regarding the inference of extinction rates[43,209]. Branch-length variations across gymnosperms clearly show that there are heterogeneous patterns of diversification, but different interpretations of extinction history might be made from the same phylogeny. For example, Leslie et al.[21] interpreted the long, isolated branches of Southern Hemisphere conifers (SHC) to reflect greater survival (and less turnover) compared to Northern Hemisphere conifers (NHC). However, based on the fossil record of SHC, others have argued that SHC were subjected to considerable Cenozoic extinctions[210,211], which suggests that their long/isolated branches might also be a product of recent 'pruning'.

In contrast, in NHC and in cycads there is a prevalence of recent radiations, with many subtended by long, otherwise depauperate branches. With respect to NHC, Leslie et al.[21] attributed this pattern to higher rates of species turnover, perhaps in response to climatic cycles over the latter Neogene. We suggest that, in addition to, or prior to, high recent turnover, climatic changes over the latter part of the Cenozoic might have generally facilitated the expansion and diversification of multiple gymnosperm lineages that, previously, had experienced high rates of extinction, as is suggested by cases where a long, depauperate branch subtends a Neogene 'comb' of numerous young species. Alternatively, this type of branching pattern might be interpreted as a 'fuse'[212], but the high historical diversity of gymnosperms shown by the fossil record suggests a 'fuse' model might generally be implausible.

Although here we argue that multiple areas of gymnosperm phylogeny have undergone recent increases in diversification, a recent study by Condamine et al.[54] suggested the opposite—high rates of recent conifer extinction (and generally negative diversification rates) precipitated by climatic cooling—based largely on analyses of fossil data. However, there are several fundamental problems with the Condamine et al.[54] study that undermine their results. One is that they only analyzed fossil occurrence data at the 'generic' level. Not only is it generally inappropriate to explicitly use ranks as a basis for such analyses, but also it is clear that most of the recent instances of gymnosperm diversification (as shown by molecular phylogenies) have occurred *within* the clades we call genera. Thus, it is no surprise that fossil occurrences collapsed to the generic level would should minimal recent origination in the context of a diversification analysis. Another major problem is their unvetted use of plant fossil data from the Paleobiology Database (PBDB). While the PBDB is an excellent data resource for various clades across the Tree of Life (e.g., Hsieh et al.[213]), the plant data are notoriously incomplete and poorly curated[214] and should generally be viewed as highly unreliable for direct use in any type of analysis. In addition to ubiquitous problems of identification, only a small portion of the paleobotanical data accumulated over the history of the field is represented in the PBDB. None of this is to say that there have not been recent, climate-driven extinctions in various gymnosperm lineages (e.g., SHC[215]), exacerbated by competition with angiosperms, but the analyses of Condamine et al.[54] are not able to speak to this question.

### 4. Supplementary Figures

**Supplementary Fig. 1 *Ks* plots for within-taxon paralog pairs (black lines) and between-taxon ortholog pairs (blue lines)**. The taxa capture the root nodes of (A) Cupressaceae, (B) Podocarpaceae, (C) Taxaceae, and (D) Pinaceae, respectively.

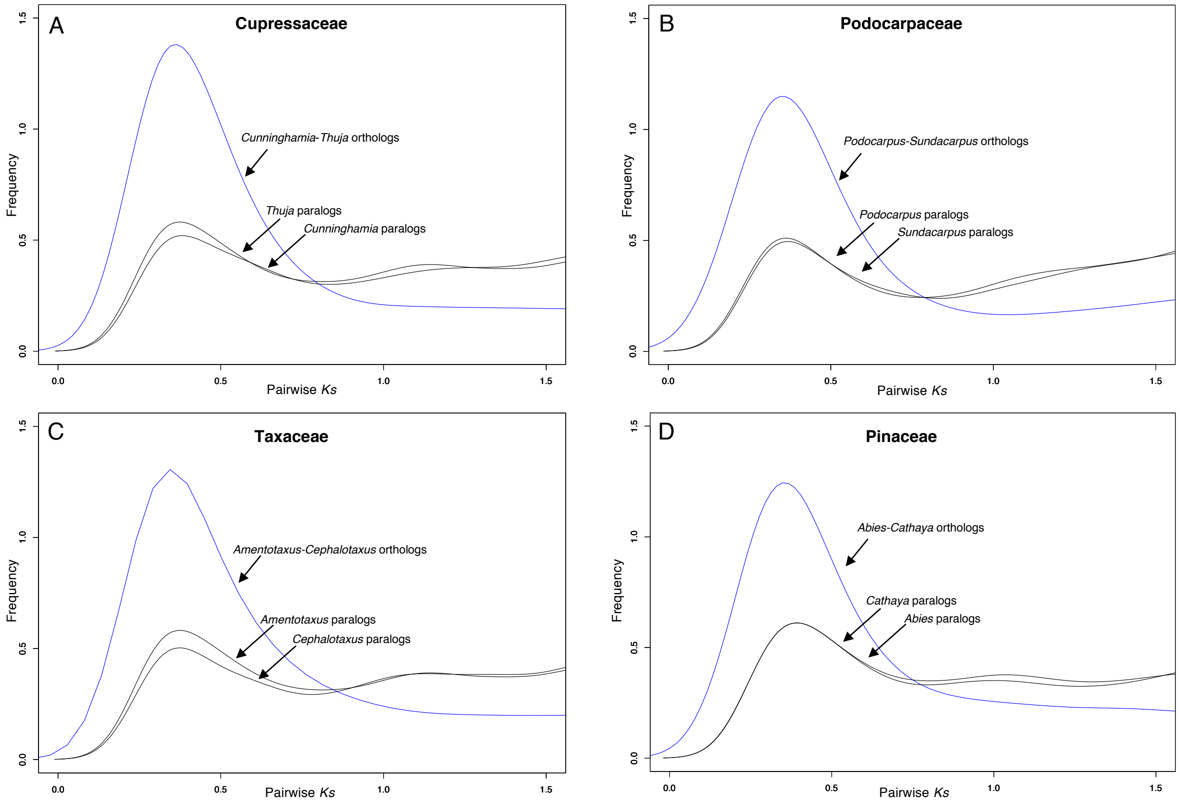

**Supplementary Fig. 2 Ancestral reconstruction of chromosome number**. Reconstructions of chromosome number were performed in ChromEvol using the dated supermatrix phylogeny and chromosome data obtained from the Chromosome Count Database (http://ccdb.tau.ac.il/).

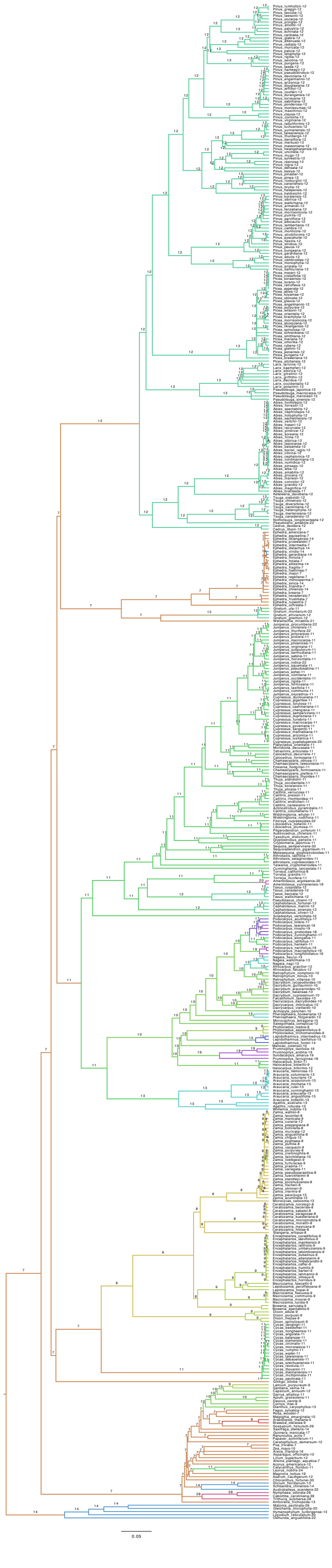

**Supplementary Fig. 3 Gymnosperm diversification**. Dated gymnosperm supermatrix phylogeny showing BAMM diversification rate shifts from the best shift configuration (i.e., the shift configuration with the maximum a posteriori [MAP] probability).

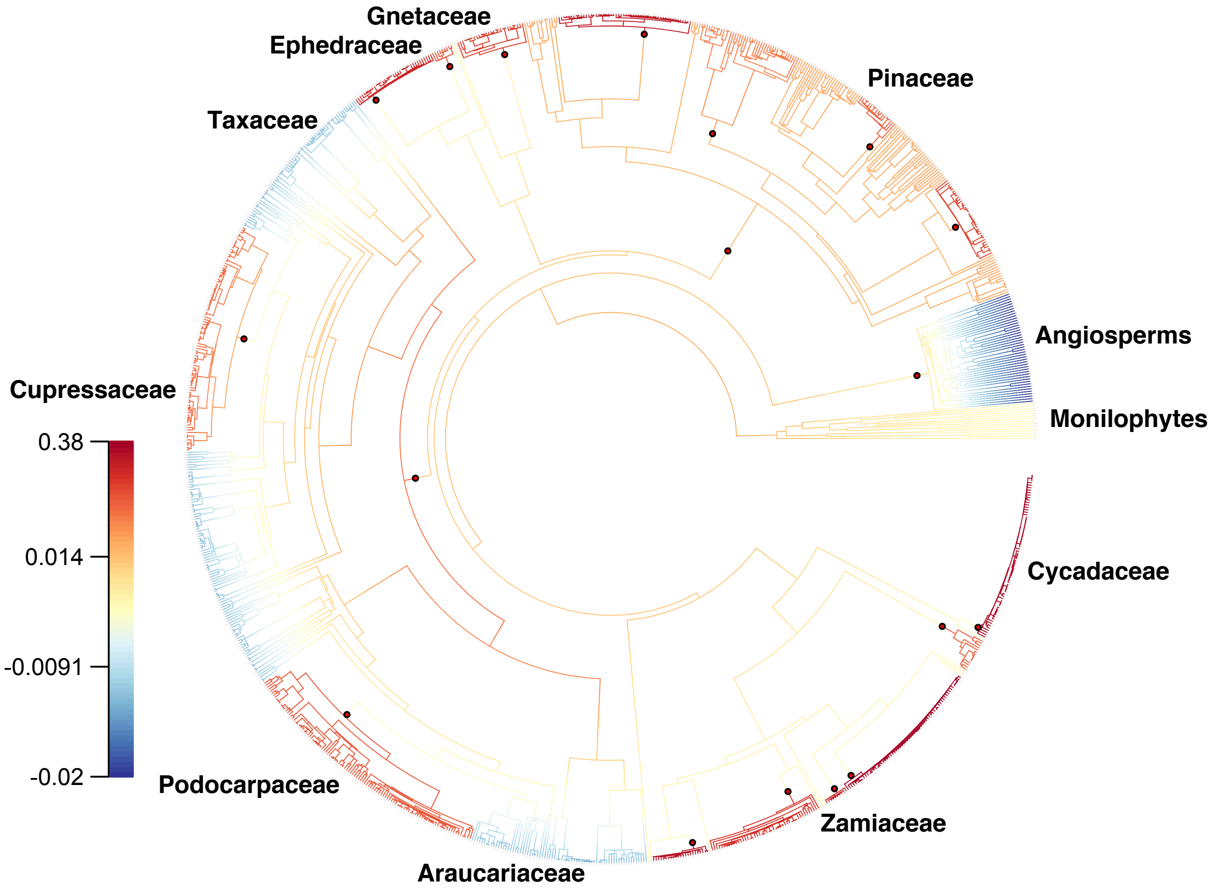

**Supplementary Fig. 4 Linear regression analysis of phenotypic innovation vs. gene duplications**. In this analysis ($n$ = 117), two outlier nodes (Gnetales and *Coniferae*) were excluded from the analysis. Justification for their exclusion is presented in the Main Text and Methods. We also conducted an analysis including all nodes (see Supplementary Fig. 21).

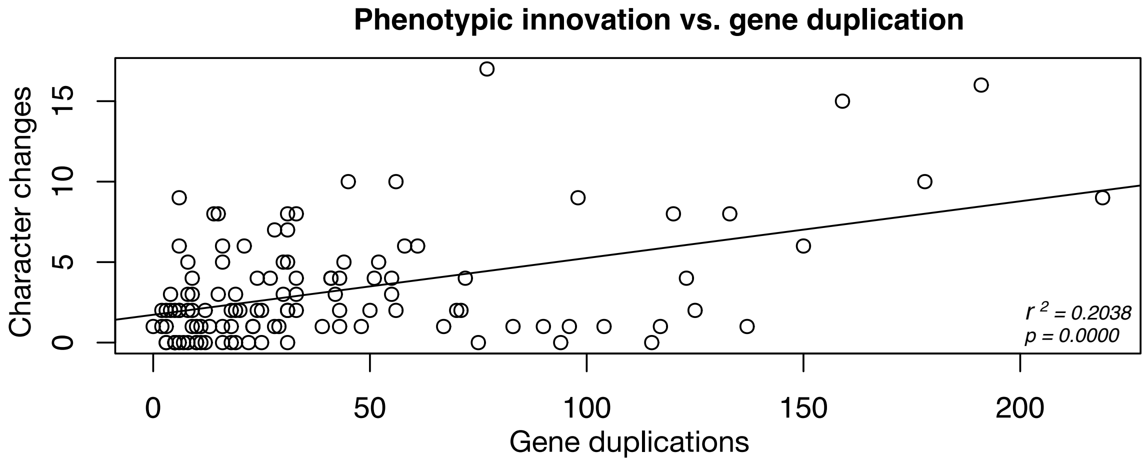

**Supplementary Fig. 5 Permutation test of phenotypic innovation vs. gene duplication**. The grey bars represent the null distribution of expected values (i.e., the expected frequency of co-occurrence of significant innovation and duplication values across all nodes); the dotted line represents the observed value (*n* = 119).

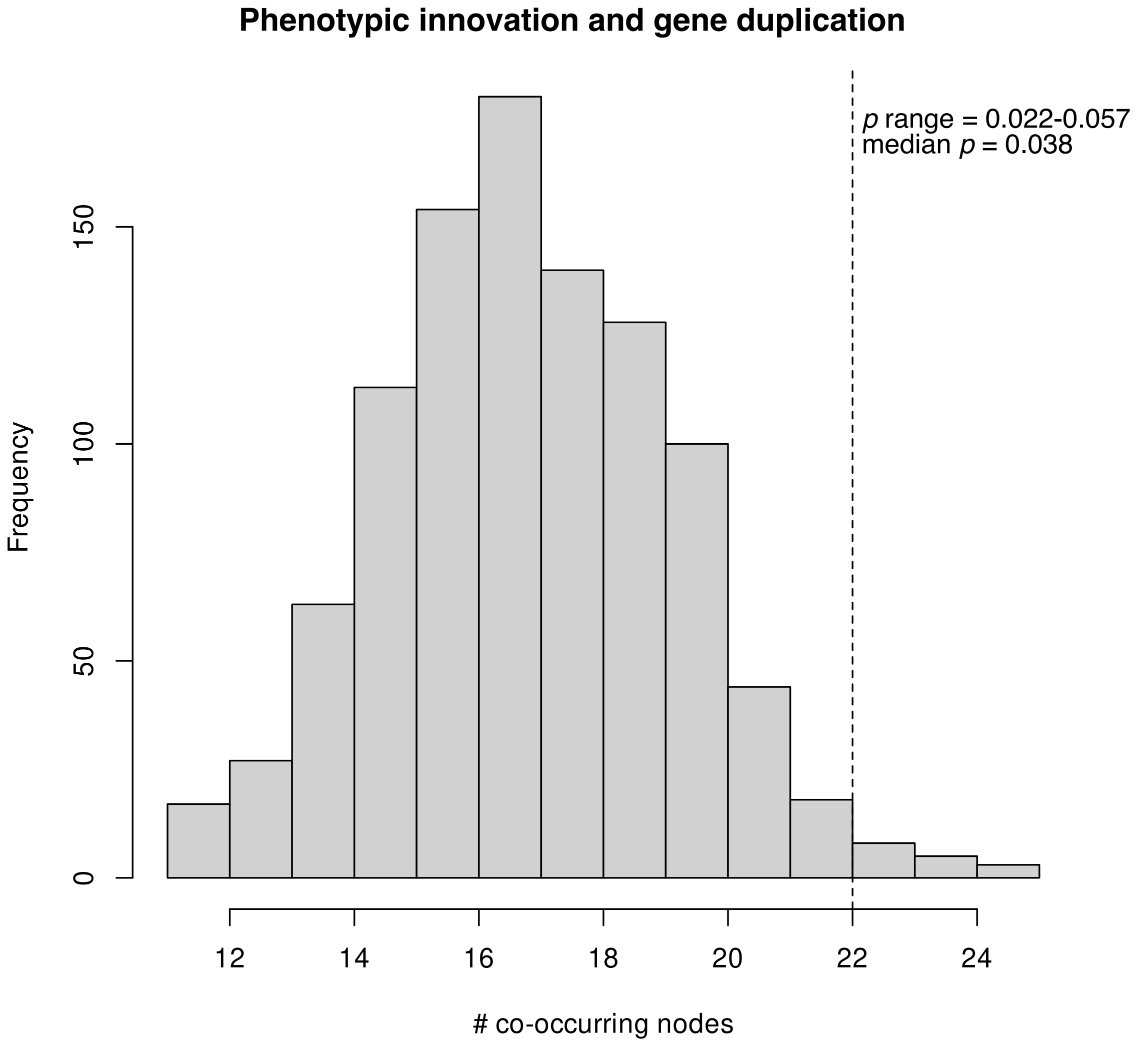

**Supplementary Fig. 6 Impact of sampling on phenotypic reconstructions**. Plots juxtaposing levels of gene duplication with levels of phenotypic innovation. The phenotypic plot shows values for each node when Gnetales were included in the analysis (dot without asterisk; $n = 119$) and when Gnetales were excluded from the analysis (dot with asterisk; $n = 118$).

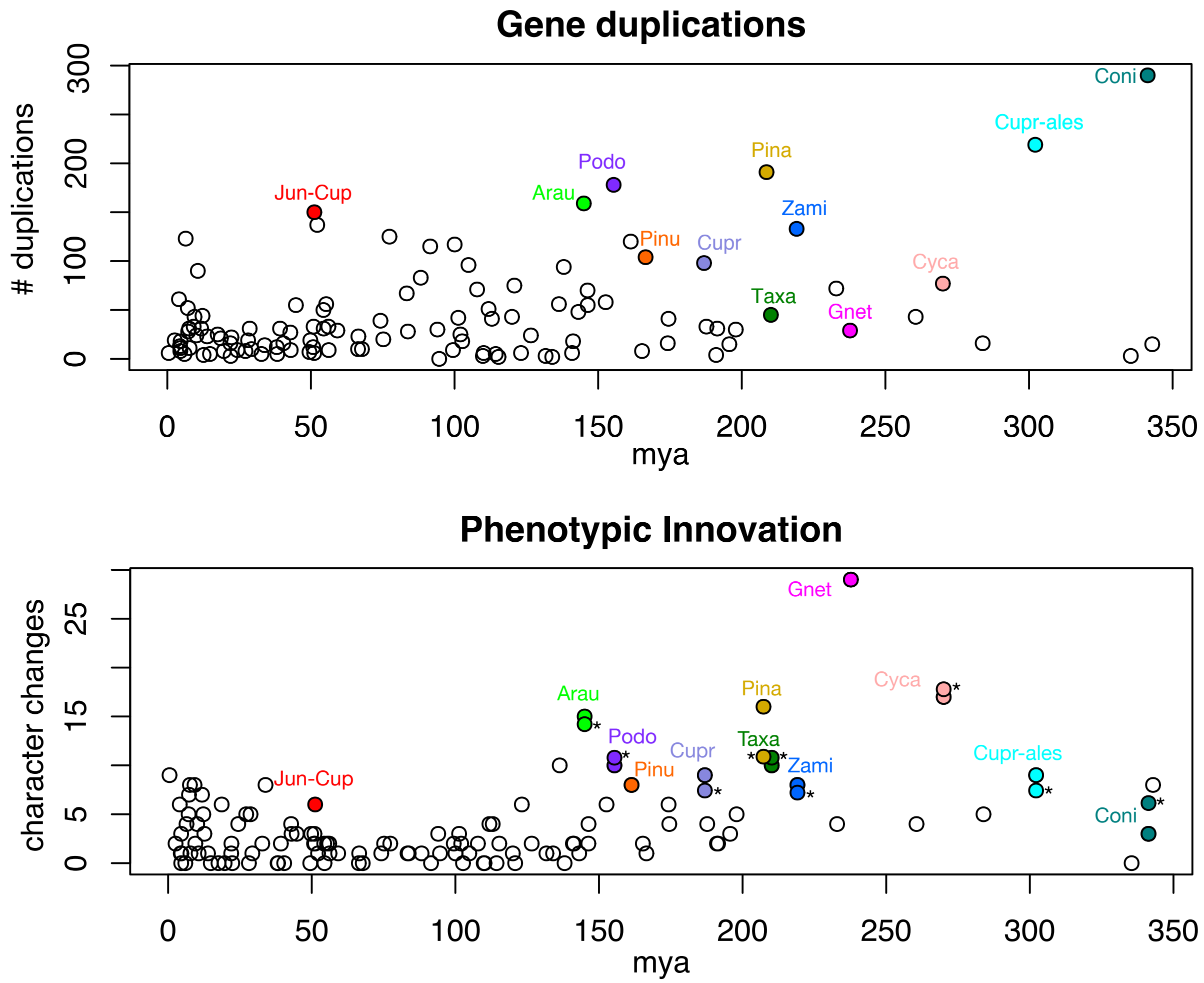

**Supplementary Fig. 7 Linear regression analysis of phenotypic rates vs. gene-tree conflict levels**. All gymnosperm nodes were included in this analysis (*n* = 119).

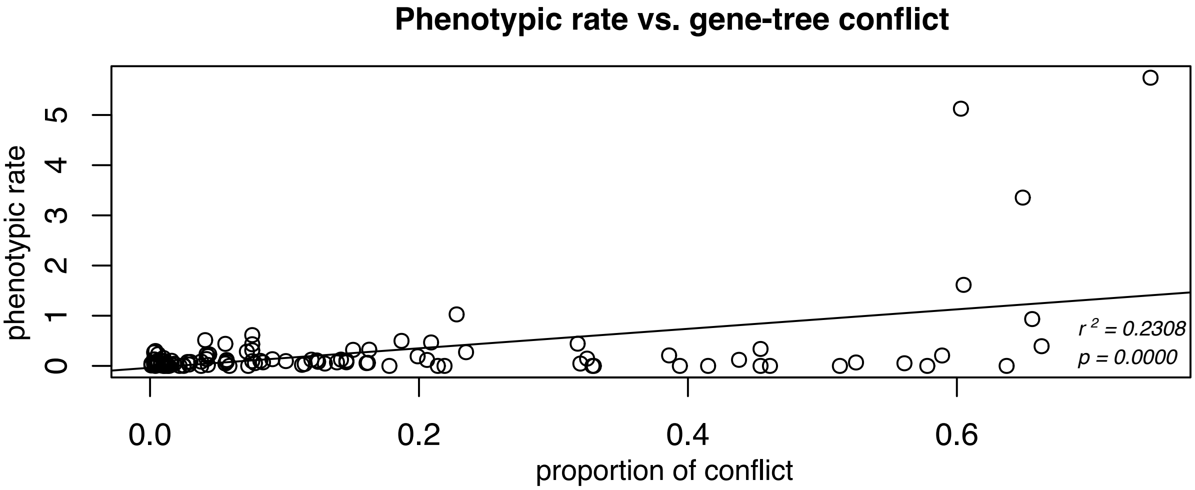

**Supplementary Fig. 8 Permutation test of phenotypic rate vs. gene-tree conflict**. The grey bars represent the null distribution of expected values (i.e., the expected frequency of co-occurrence of significant rate and conflict values across all nodes); the dotted line represents the observed value (*n* = 119).

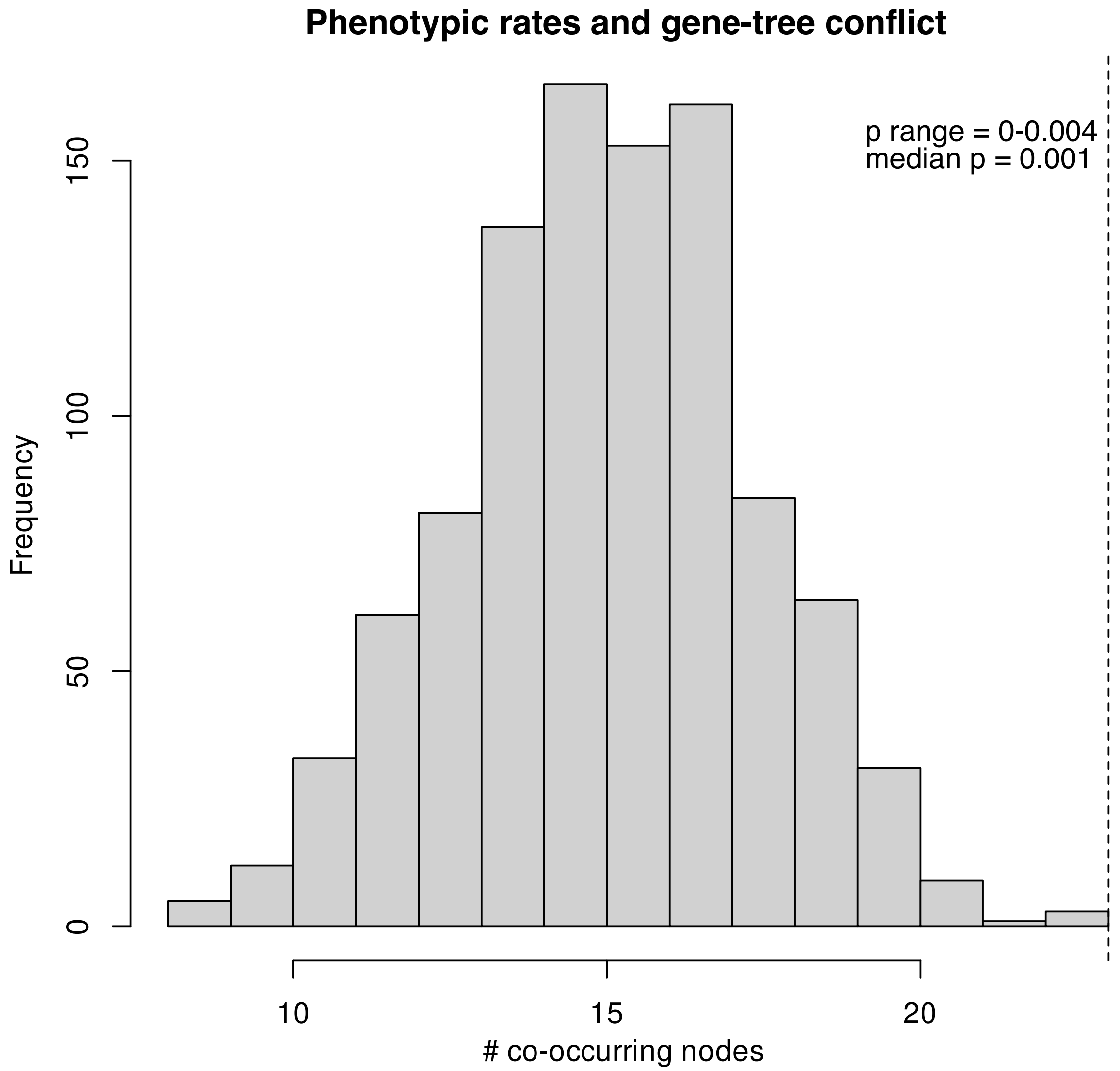

**Supplementary Fig. 9 Rate shifts in climatic evolution (Bio 1, mean annual temperature)**. The analysis was conducted using BAMM. The shifts plotted are those from the MAP shift configuration.

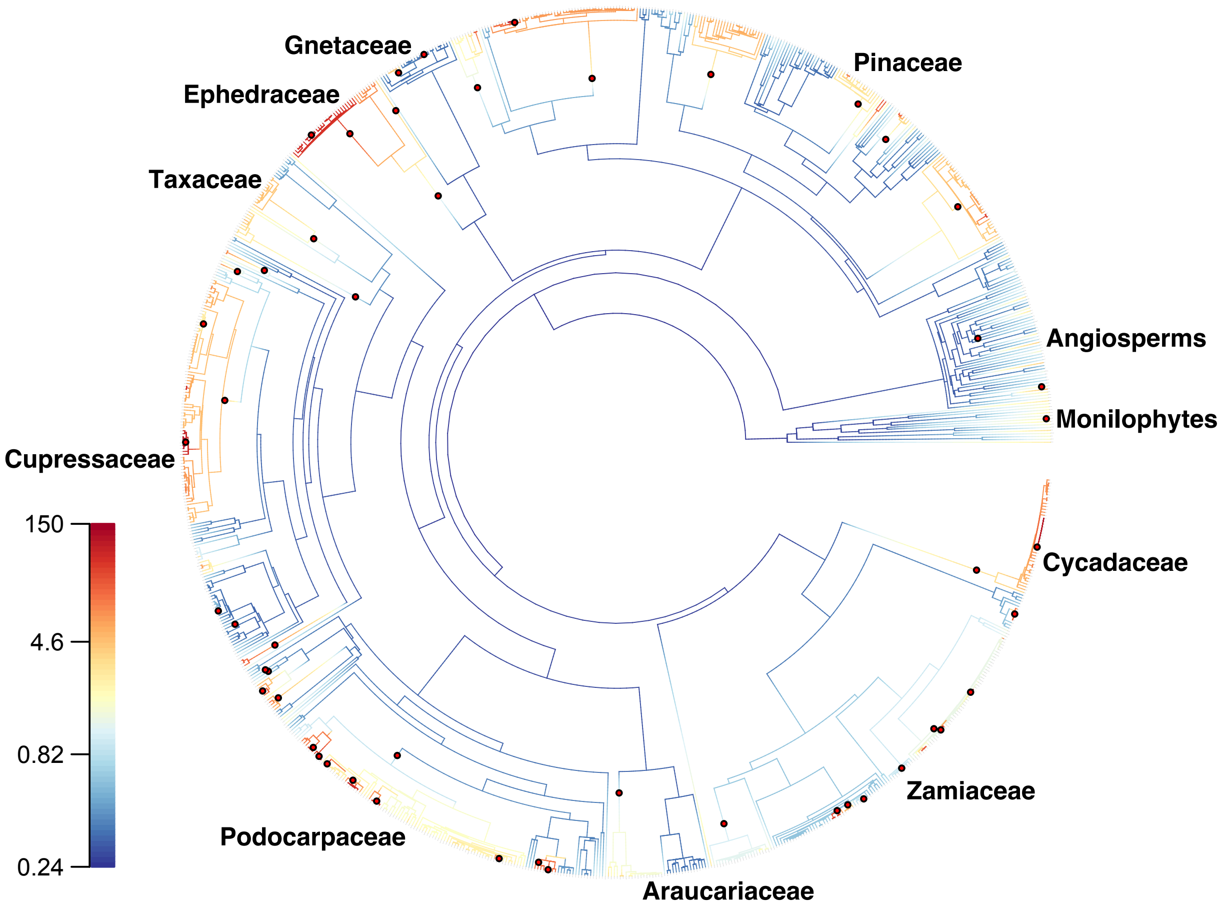

**Supplementary Fig. 10 Rate shifts in climatic evolution (Bio 12, annual precipitation)**. The analysis was conducted using BAMM. The shifts plotted are those from the MAP shift configuration.

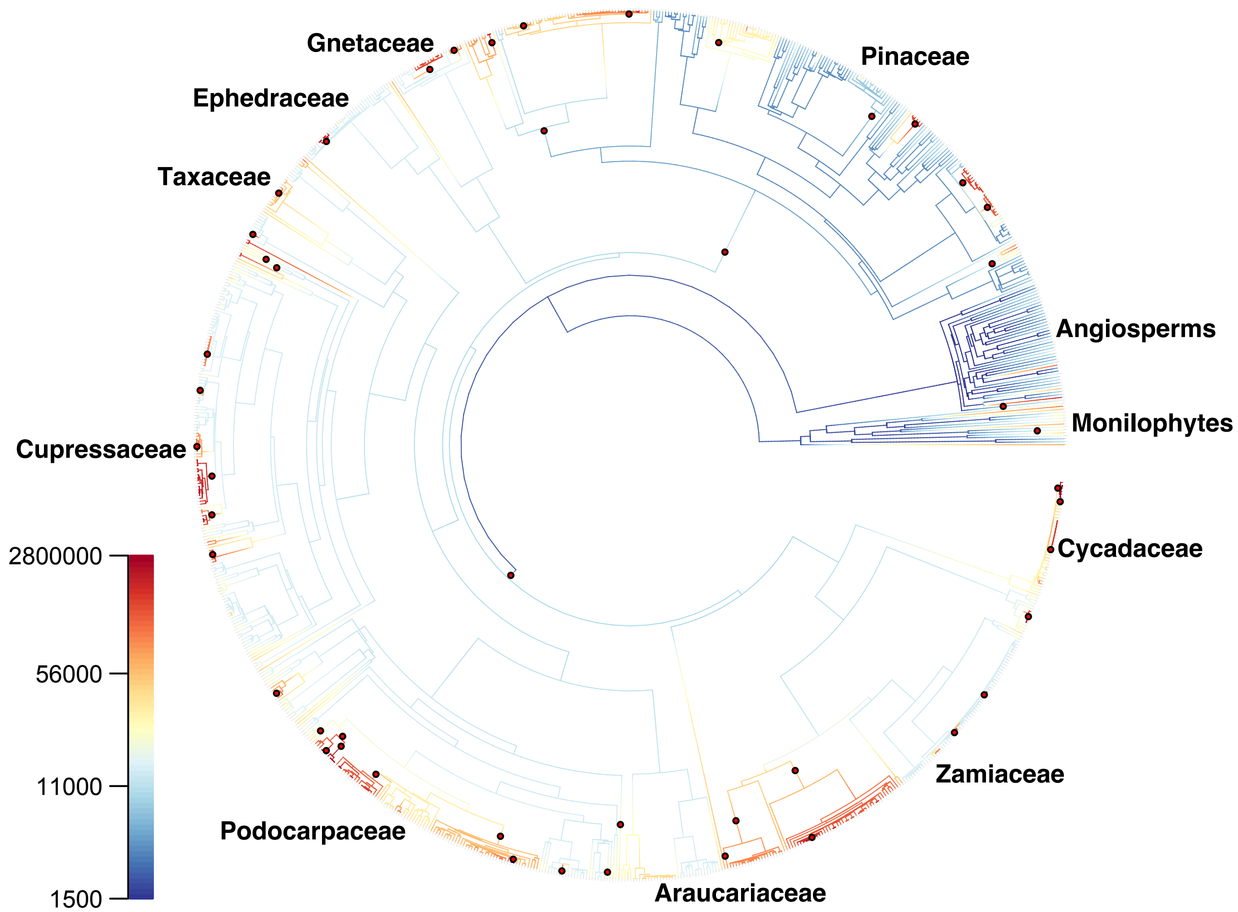

**Supplementary Fig. 11 Rate shifts in climatic evolution (principle component one of the ordinated Bioclimatic variables)**. The analysis was conducted using BAMM. The shifts plotted are those from the MAP shift configuration.

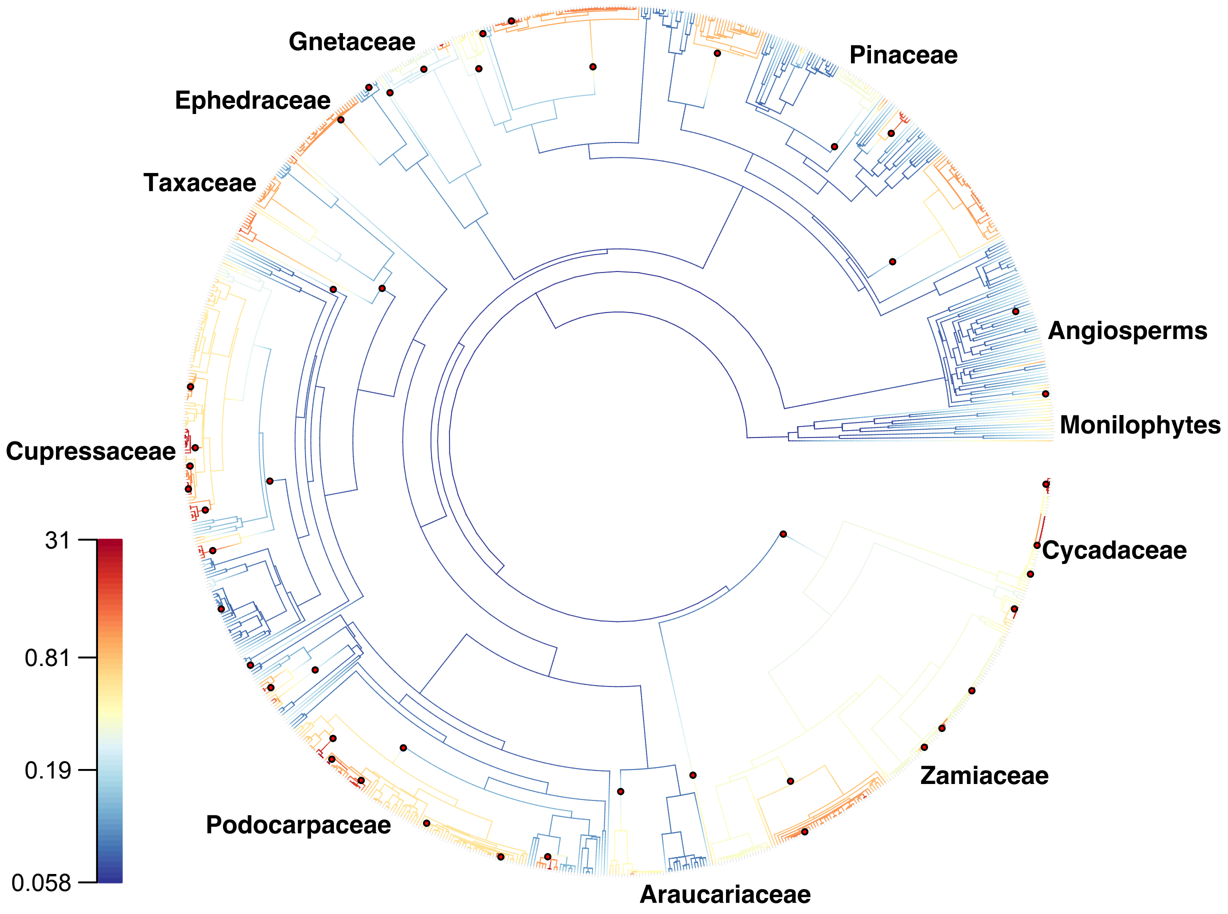

**Supplementary Fig. 12 Rate shifts in climatic evolution (principle component two of the ordinated Bioclimatic variables)**. The analysis was conducted using BAMM. The shifts plotted are those from the MAP shift configuration.

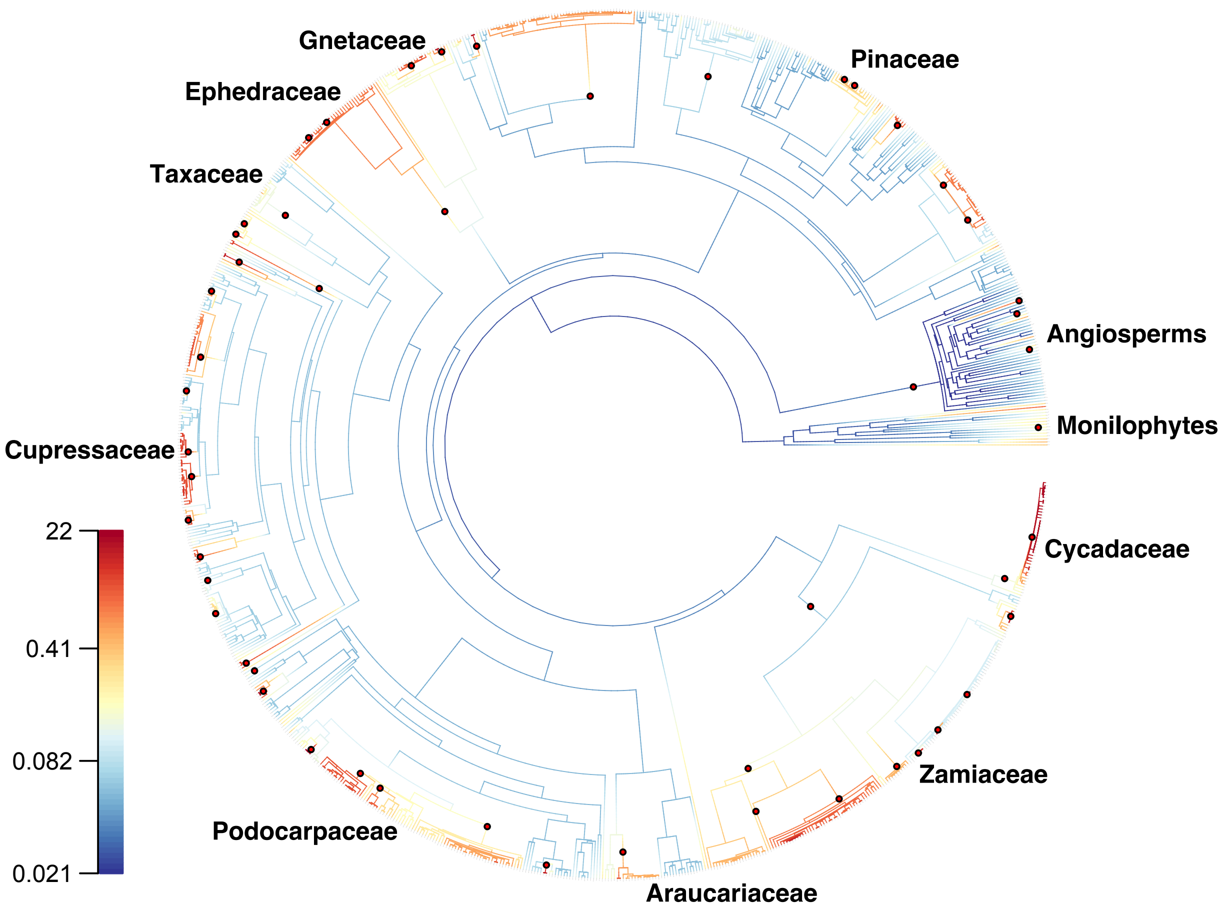

**Supplementary Fig. 13 Major jumps in climatic evolution (Bio 1 and 12)**. Ancestral reconstruction of mean annual temperature (Bio 1) on the dated supermatrix phylogeny, showing climatic jumps (i.e., extreme parent node-child node differences) in mean annual temperature (Bio 1, solid circles) and annual precipitation (Bio 12, triangles), as well as diversification rate shifts inferred from BAMM (open circles).

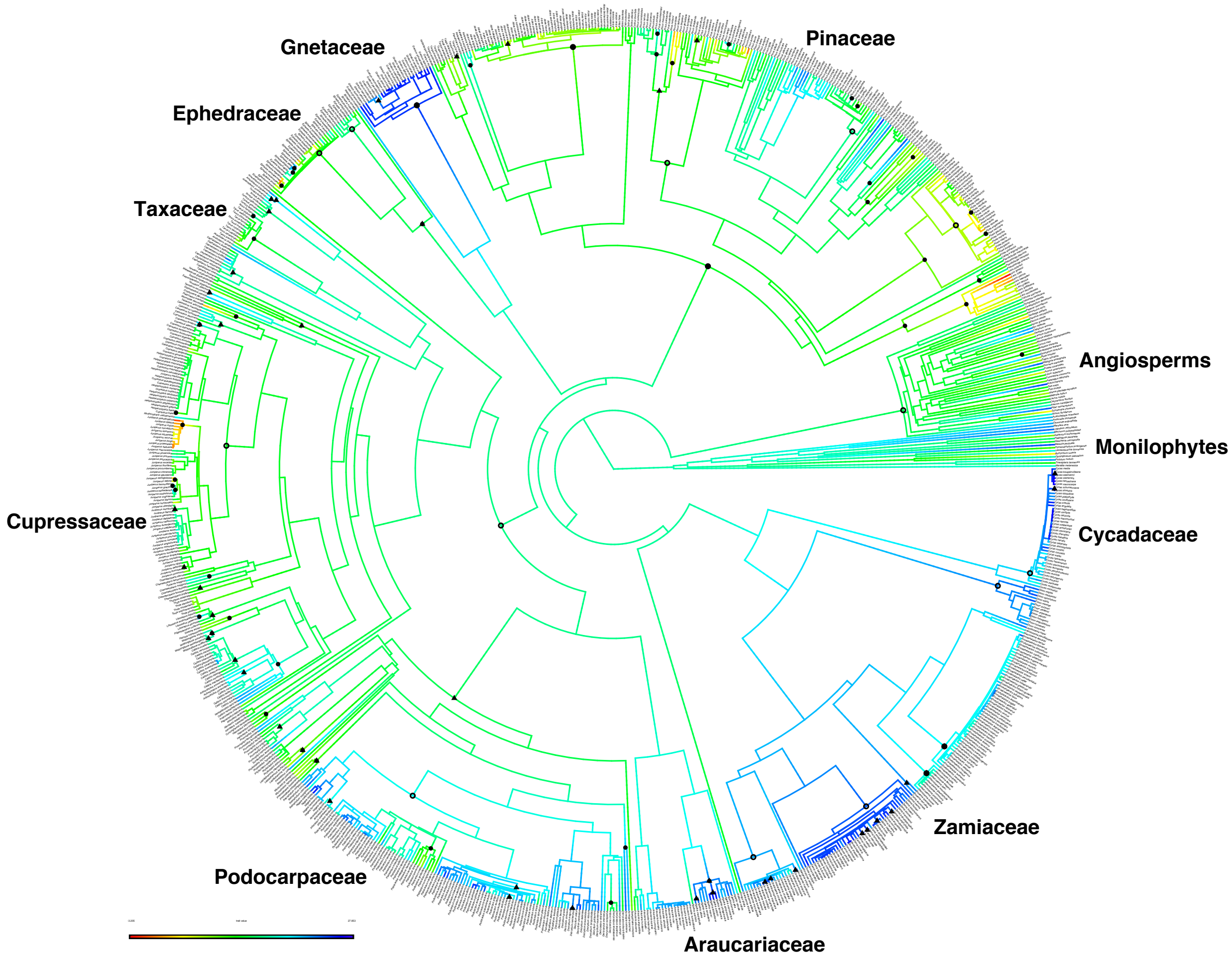

**Supplementary Fig. 14 Major jumps in climatic evolution (PC1 and PC2)**. Ancestral reconstruction of mean annual temperature (Bio 1) on the dated supermatrix phylogeny, showing climatic jumps (i.e., extreme parent node-child node differences) in principle component one (solid circles) and principle component two (triangles), as well as diversification rate shifts inferred from BAMM (open circles).

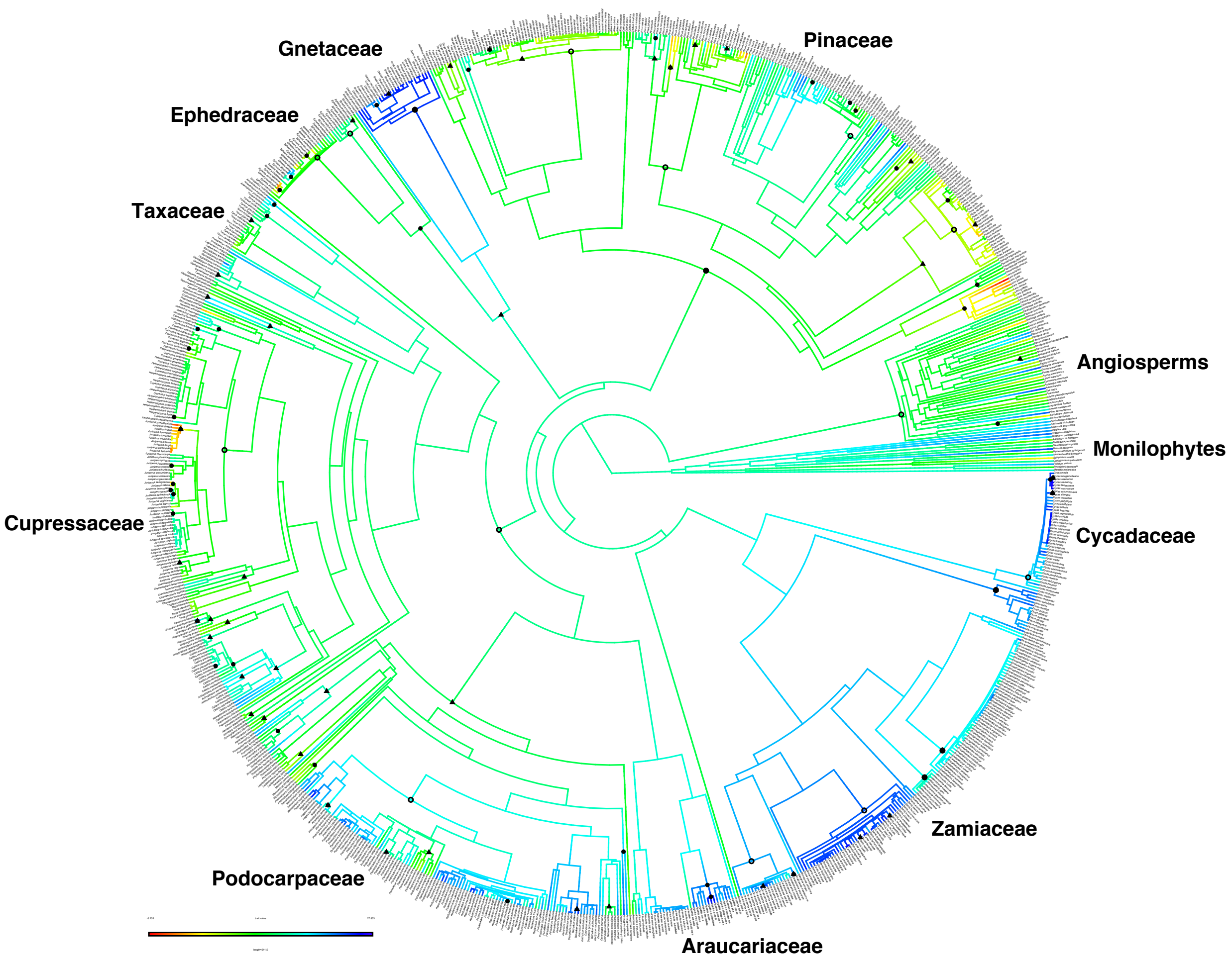

**Supplementary Fig. 15 Genome size evolution**. This is the same tree shown in Extended Data Fig. 3, but with the tips labeled, for more detailed reference.

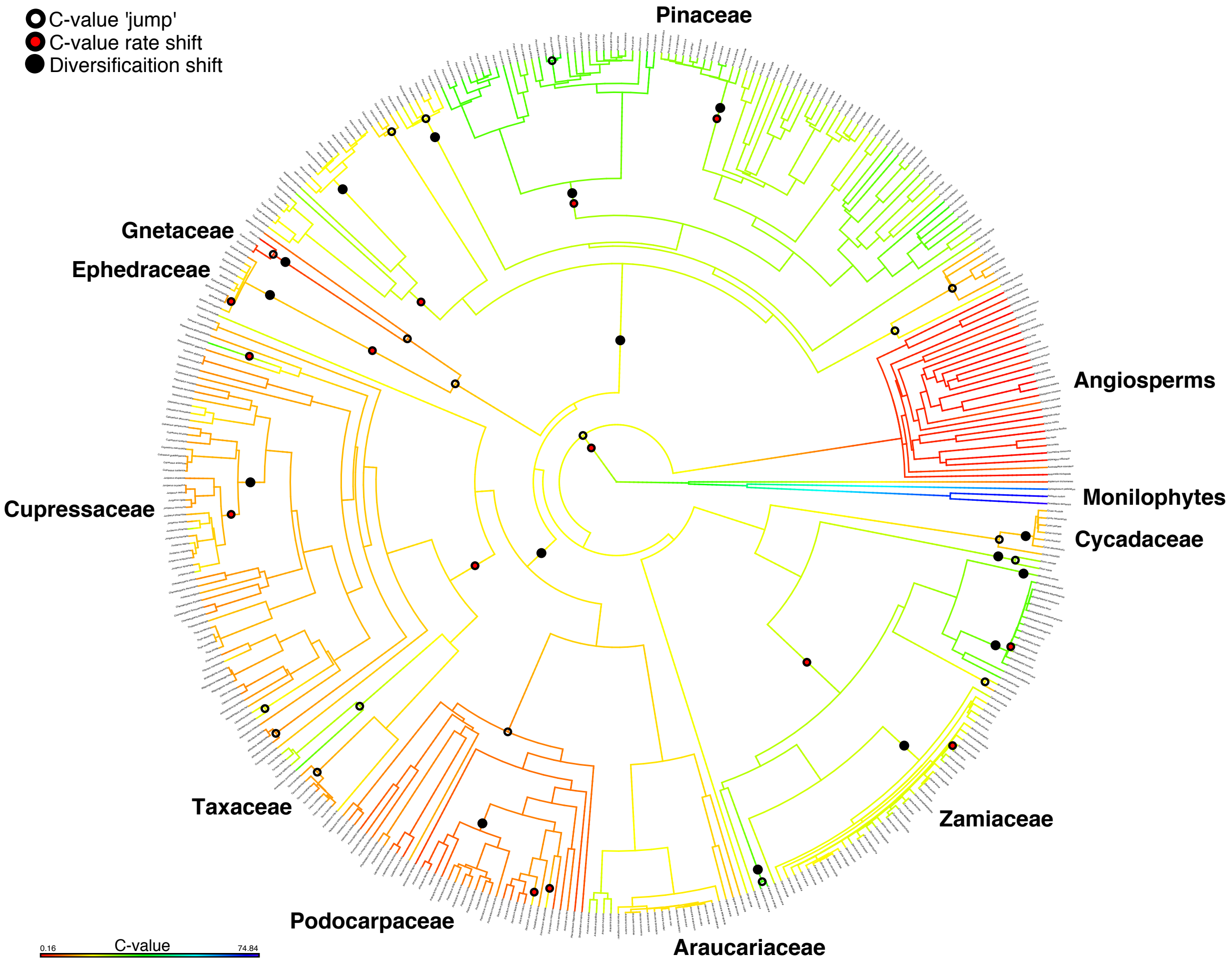

**Supplementary Fig. 16 ASTRAL phylogeny**. The ASTRAL analysis was based on the 790 nuclear loci from the transcriptomic and genomic dataset. All branches are fully supported unless otherwise indicated.

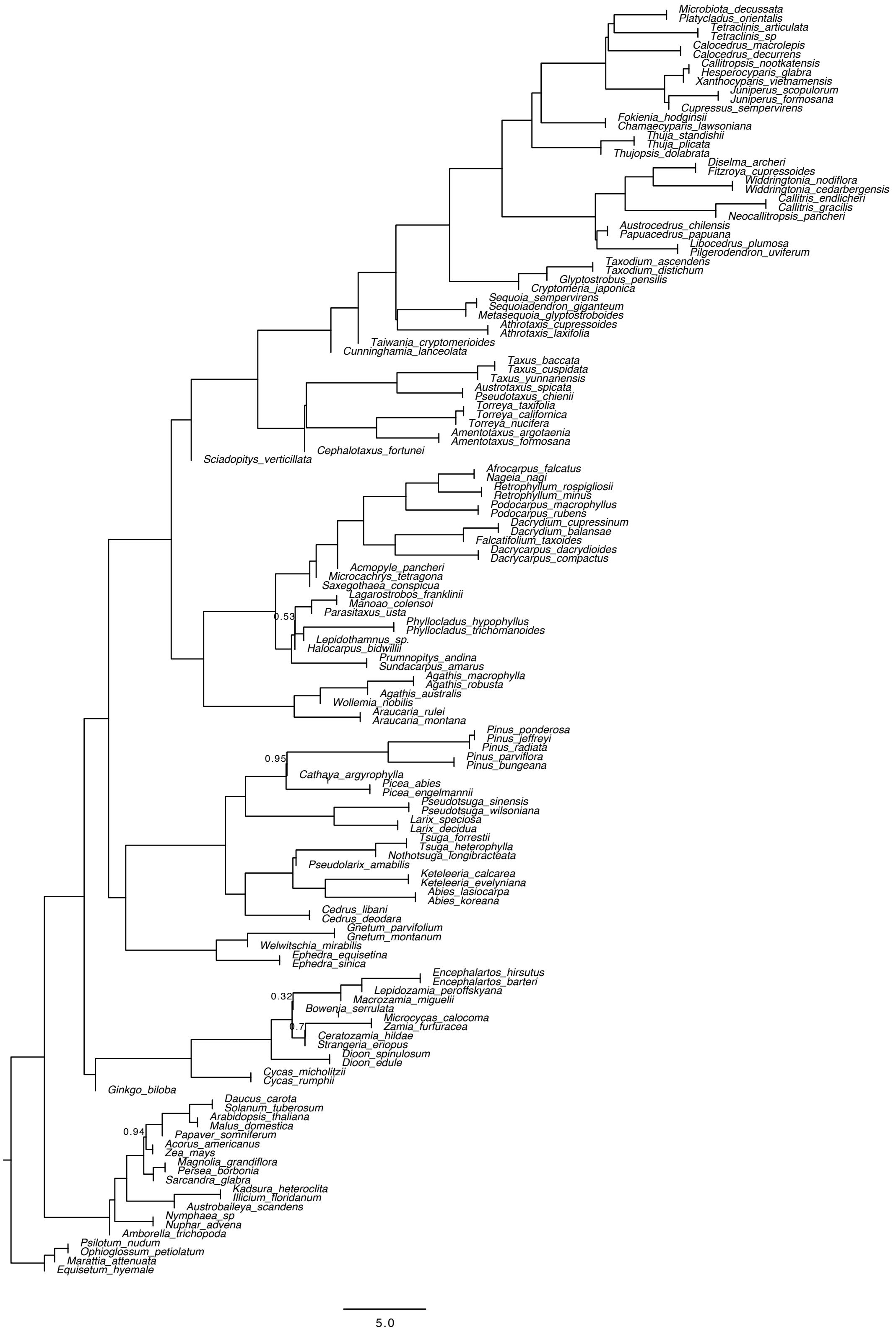

**Supplementary Fig. 17 Concatenation phylogeny**. Phylogeny inferred from the concatenated nuclear loci using IQTREE, without model partitioning. Branches have maximal support unless otherwise indicated.

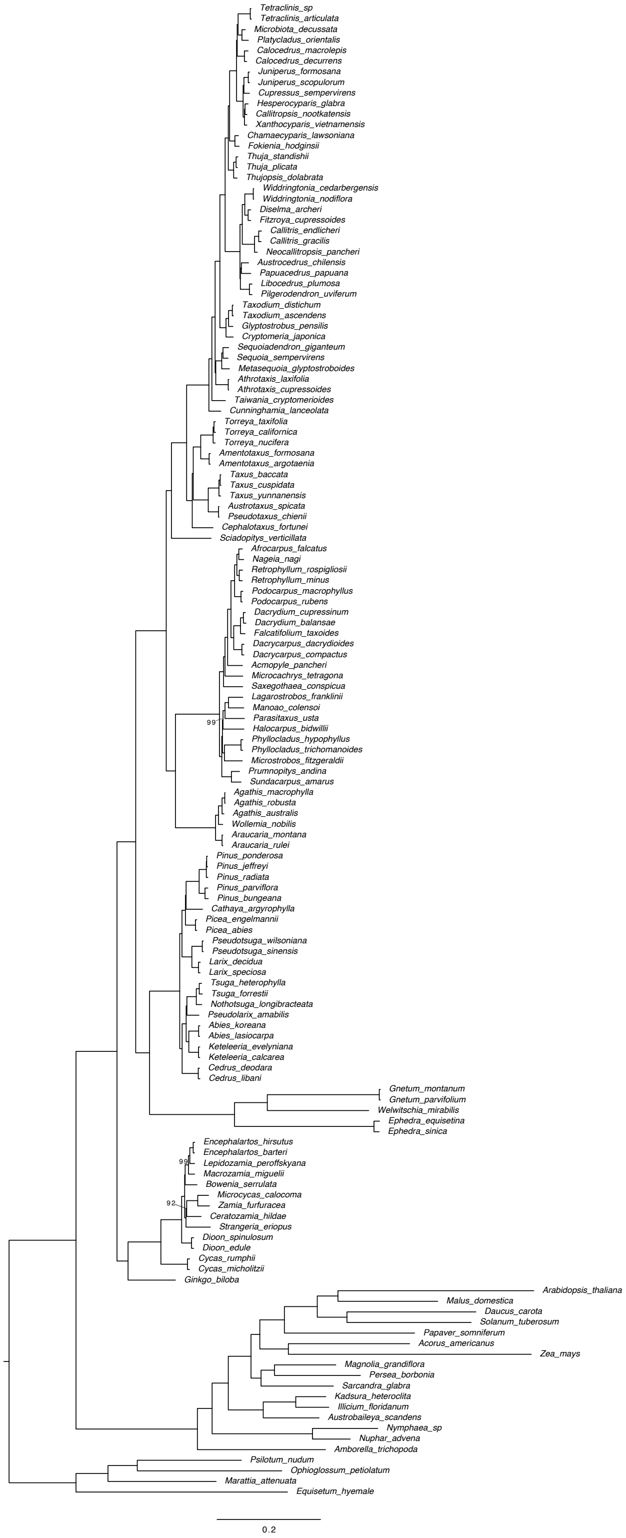

**Supplementary Fig. 18 Gene-duplication mapping using the MRCA method**. Gymnosperm phylogeny (inferred using ASTRAL) showing percentages of gene-duplication at each node, inferred using the MRCA method of Yang et al.[28]

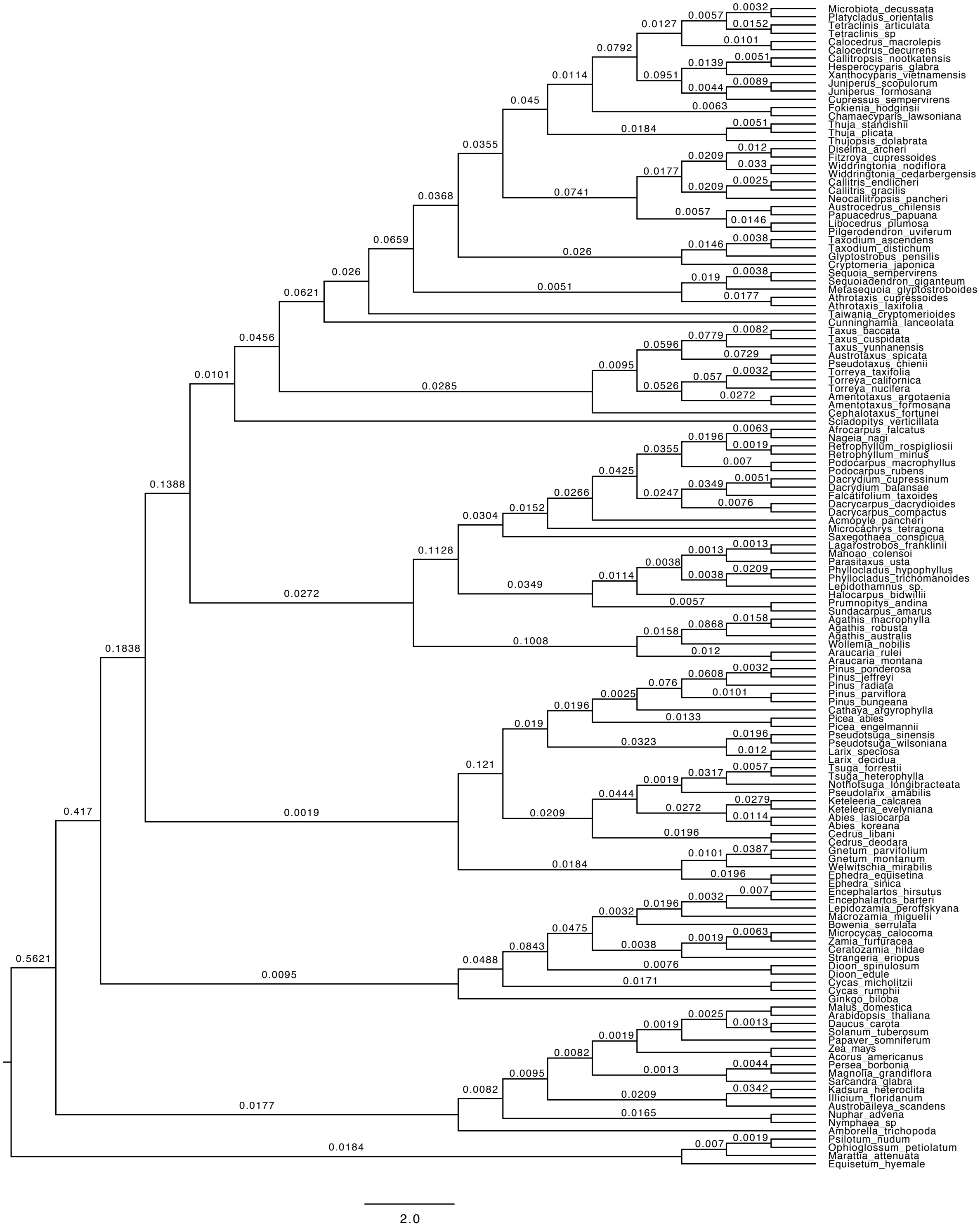

**Supplementary Fig. 19 Gene-duplication mapping using the concordance method**. Gymnosperm phylogeny (inferred using ASTRAL) showing percentages of gene-duplication at each node, inferred using the concordance method of Yang et al.[28]

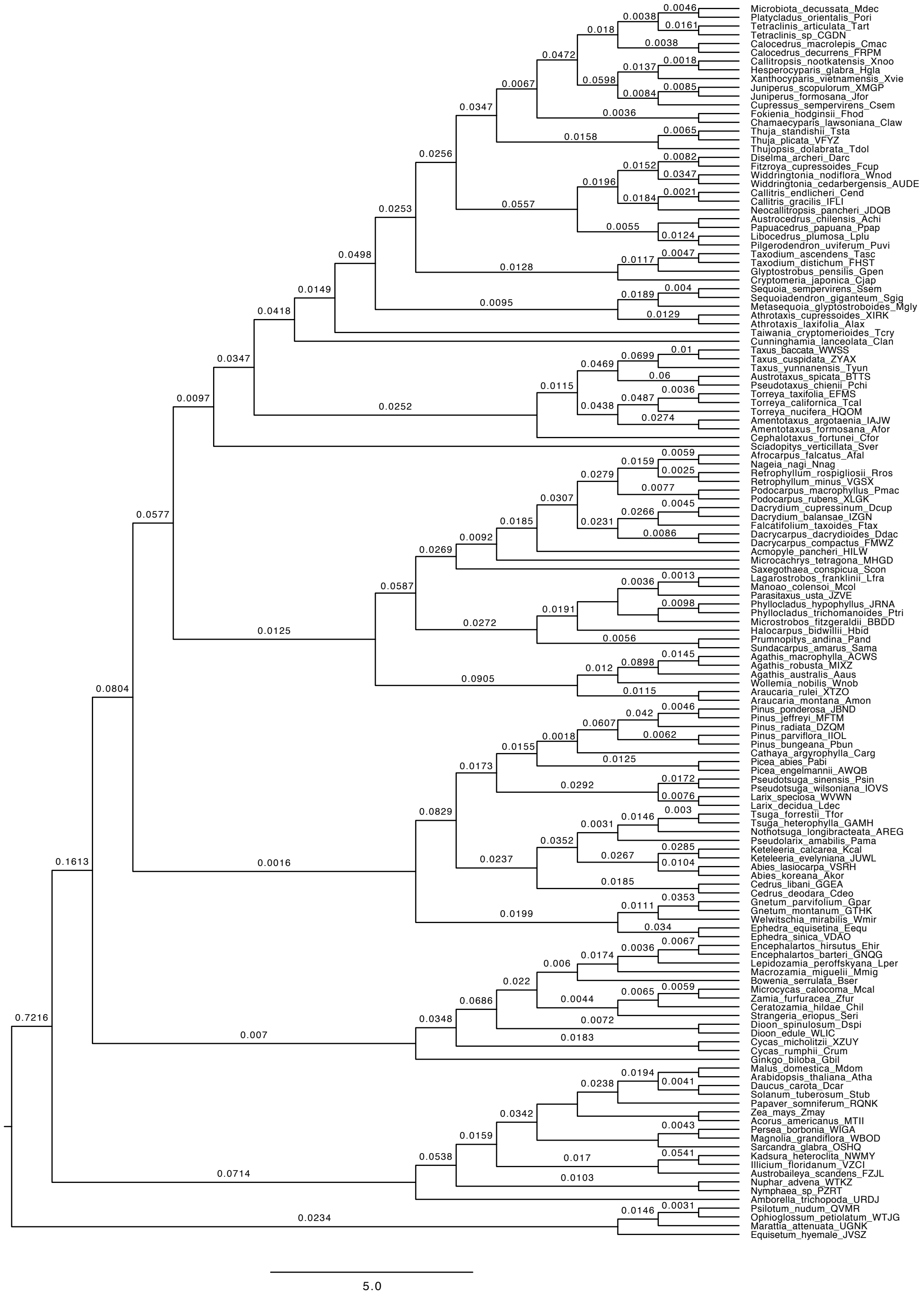

**Supplementary Fig. 20 Variation in divergence-time estimates**. The trees shown are the 25 independently dated ML trees, plotted using the 'densiTree' function in phangorn[216]. The individual ML trees were dated using treePL.

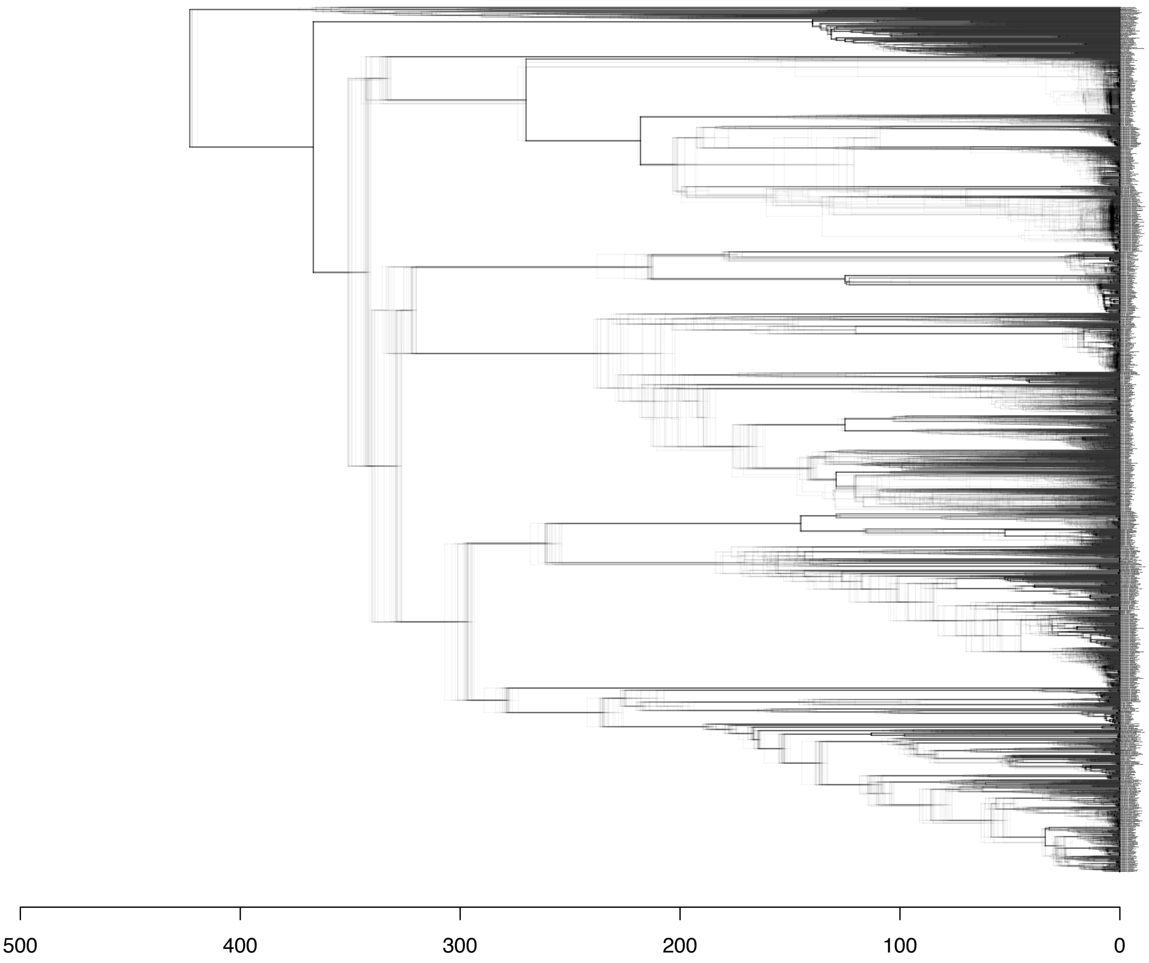

**Supplementary Fig. 21 Linear regression analysis of phenotypic innovation vs. gene duplications**. In this analysis ($n$ = 119), all gymnosperm nodes were included (compare with Supplementary Fig. 4)

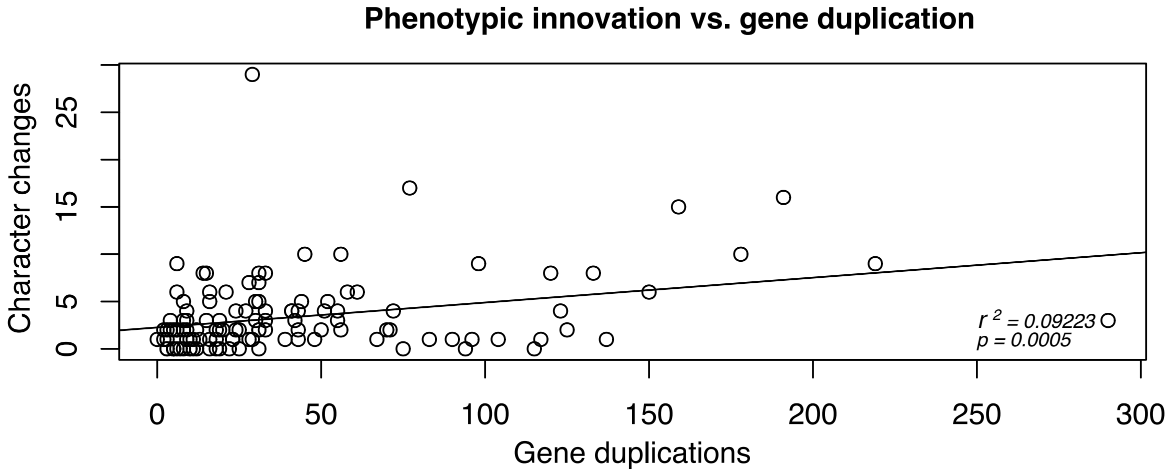

**Supplementary Fig. 22 Permutation test of diversification vs. phenotypic innovation**. The grey bars represent the null distribution of expected values (i.e., the expected frequency of co-occurrence of diversification shifts and significant innovation values across all nodes); the dotted line represents the observed value (*n* = 119).

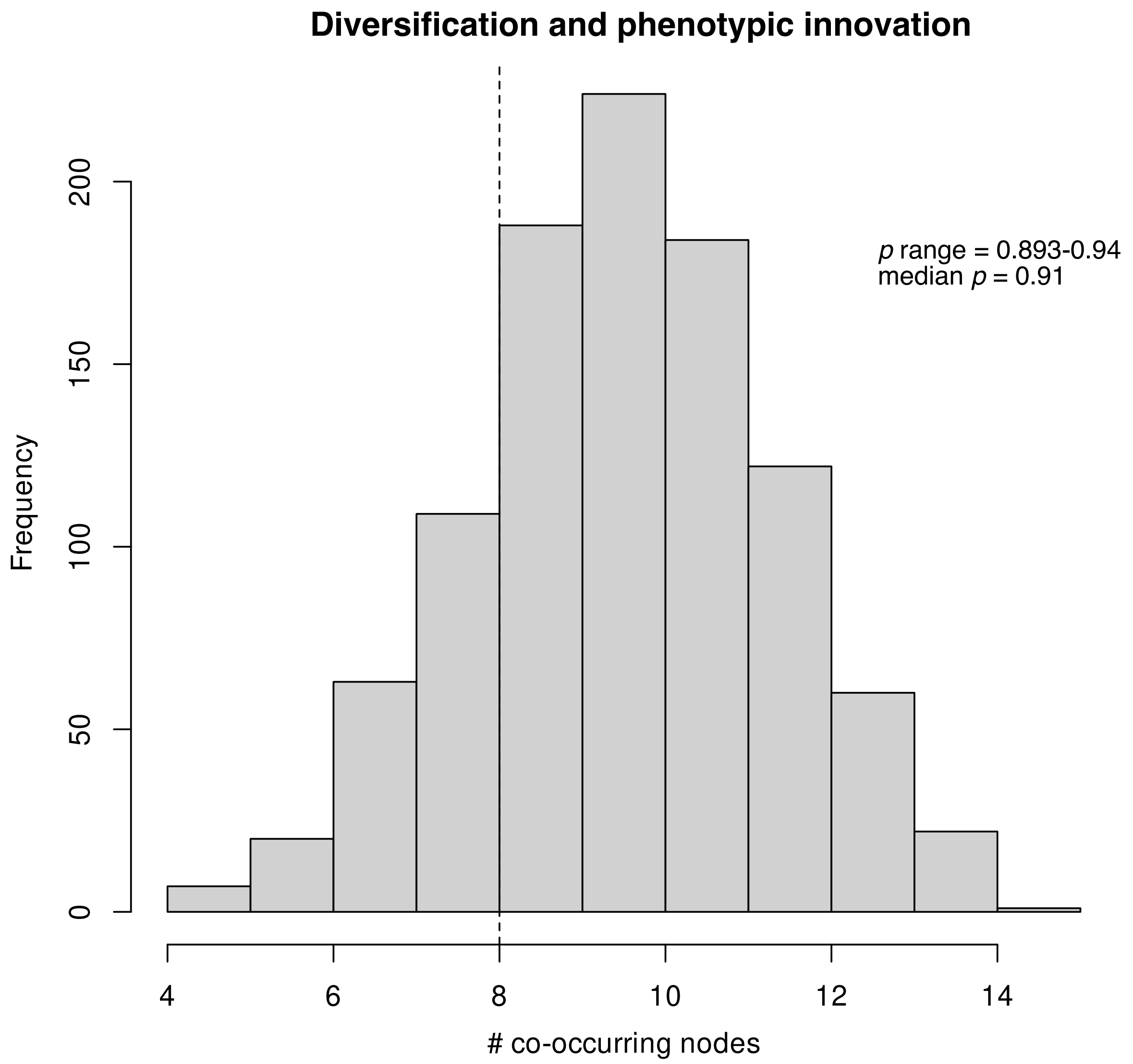
